# GSNOR-dependent nitric oxide homeostasis promotes recovery from repeated climate stress across generations in *Arabidopsis thaliana*

**DOI:** 10.64898/2026.08.05.742973

**Authors:** R Behl, A Ghirardo, JB Winkler, S Jafarian, L Frungillo, SH Spoel, A Fischback, G Huber, R Koller, A Albert, E Mattes, L Gößl, F.X. Mayer K, F Johannes, C Lindermayr, JP Schnitzler

## Abstract

Climate change exposes plants to recurrent and interacting stresses, yet the extent to which these effects persist across generations, and the mechanisms involved, remain unclear. We propagated *Arabidopsis thaliana* wild type (WT, Col-0) and nitric oxide homeostasis mutant *gsnor1-3* (hereafter, *gsnor-ko*) for five successive generations. Plants were grown under control, drought, elevated CO_2_, O_3_, warm temperature, and combined treatment scenarios for the first three generations (G1-G3), followed by two recovery generations under control conditions (G4-G5). We quantified rosette growth, photosynthetic traits, seed production, and transcriptome dynamics by RNA-seq. Across environments, *gsnor-ko* showed reduced vegetative growth and reproductive output relative to WT. Transcriptomic responses were strongly scenario- and generation-dependent, with the largest differential expression shifts observed under warm-climate and combined-treatment conditions. Compared with WT, *gsnor-ko* displayed broader gene overlap across generations and stronger retention or reconfiguration of stress-responsive states after stress withdrawal. Functional enrichment and candidate-gene analyses identified pathways/components linked to DNA methylation, heterochromatin maintenance, histone ubiquitination, m^6^A RNA regulation, and methyl-donor metabolism. Together, these results support a model in which GSNOR activity promotes transcriptomic recovery after repeated climate stress, whereas impaired GSNOR function shifts responses toward multi-generational persistence and epigenetically associated regulatory reconfiguration.

**Highlight:** GSNOR-dependent nitric oxide homeostasis promotes transcriptomic resetting after repeated climate stress, whereas impaired NO homeostasis favours multigenerational persistence and chromatin- and RNA-linked regulatory reconfiguration.

**Graphical Abstract:** 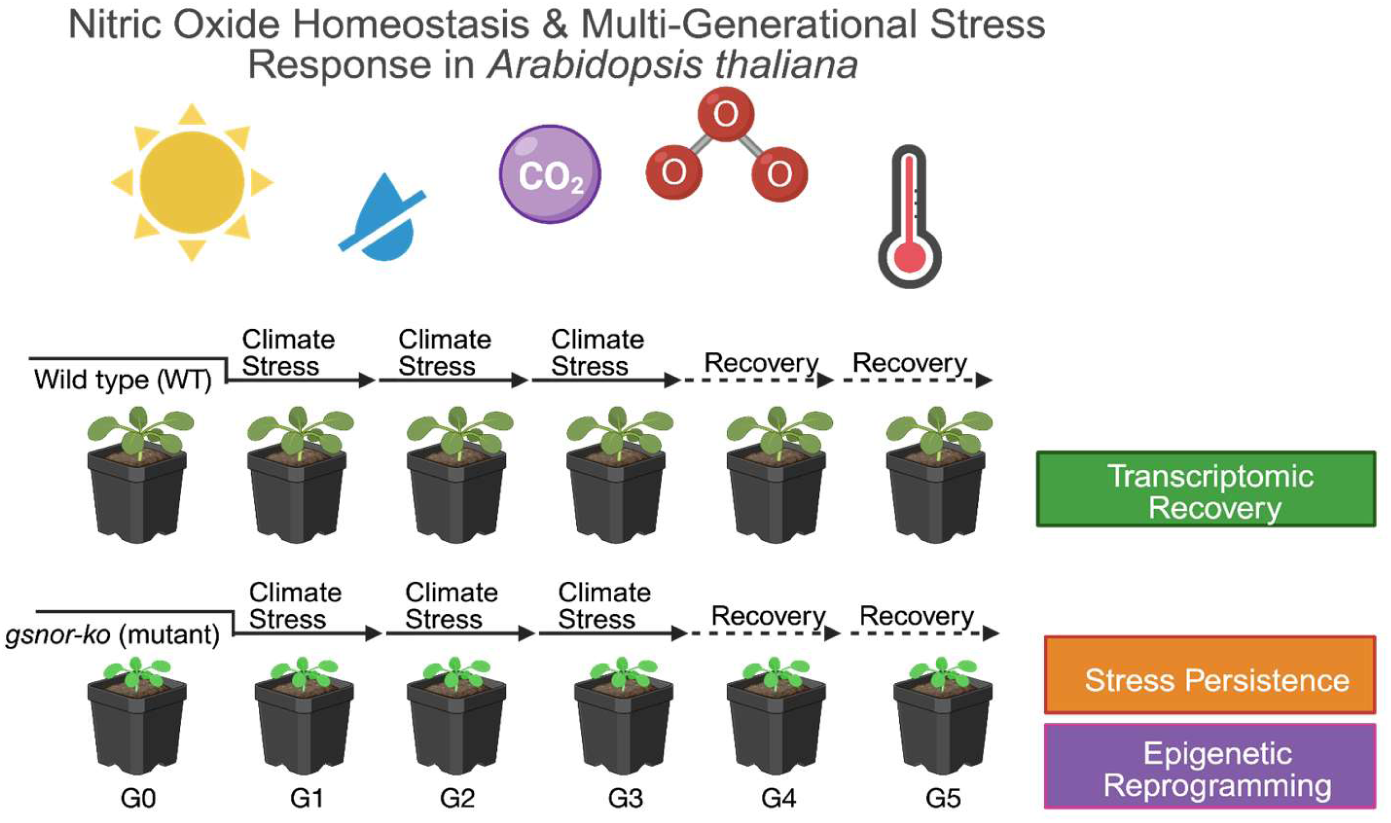

## Introduction

Climate change is reshaping the plant environment through simultaneous shifts in temperature, water availability, and atmospheric composition (IPCC, 2023a). In addition to warming and drought, plants are increasingly exposed to rising concentrations of carbon dioxide (CO_2_) and tropospheric ozone (O_3_), which can often occur together rather than as isolated stresses (IPCC, 2023b). Because multiple stressors can trigger non-additive physiological and molecular responses, understanding plant adaptation under future climates requires experimental frameworks that integrate multiple environmental drivers across time (De Kauwe *et al*., 2021; Montes *et al*., 2022; Pascual *et al*., 2022).

Plants can retain information from prior stress exposure, leading to physiological, transcriptional, or developmental stress memory that shapes subsequent responses (Zheng *et al*., 2024). However, the stability and mechanism of this memory vary widely, and a critical distinction must be made between short-term somatic memory, intergenerational effects, and transgenerational inheritance. Foundational Arabidopsis studies reported stress-associated transgenerational effects under distinct conditions (Molinier *et al*., 2006; Boyko *et al*., 2010), whereas later work also showed that stable heritable methylome changes are not a general outcome under repeated drought stress (Ganguly *et al*., 2017). Current evidence therefore strongly supports stress memory and priming in plants, but whether stress-induced states are stably transmitted through meiosis remains context-dependent and is often unresolved (Crisp *et al*., 2016; Erdmann and Picard, 2020; Pratx *et al*., 2024). Broadening this perspective, work in poplar under future-climate-relevant drought-heat scenarios showed that transcriptomic responses were similar during stress but diverged strongly after recovery, supporting the concept of stress-related molecular memory and altered post-stress steady states in a climate-change context (Georgii *et al*., 2019).

Epigenetic regulation provides a plausible molecular framework for the maintenance or resetting of stress-responsive states. In plants, this includes DNA methylation pathways mediated by methylation writers such as METHYLTRANSFERASE 1 (MET1), CHROMOMETHYLASE 3 (CMT3), and DOMAINS REARRANGED METHYLTRANSFERASE 2 (DRM2); active demethylases such as REPRESSOR OF SILENCING 1 (ROS1), DEMETER (DME), DEMETER-LIKE 2 (DML2), and DEMETER-LIKE 3 (DML3); histone-modifying enzymes that deposit or remove activating and repressive chromatin marks and histone variants that can shape nucleosome accessibility (Liu and Lang, 2020; Ueda and Seki, 2020; Nunez-Vazquez *et al*., 2022). In parallel, RNA modifications such as N^6^-methyladenosine (m^6^A) add a post-transcriptional regulatory layer through writer, reader, and eraser proteins that influence RNA processing, stability, and translation (Reichel *et al*., 2019). Together, these systems are well-positioned to connect environmental sensing with sustained changes in gene expression. Heat-stress memory, for example, is associated with prolonged transcription of memory-associated genes including APX2, HSP18.2, HSP21, HSP22, and HSA32 and chromatin-based maintenance mechanisms such as HSFA2/HSFA3-mediated H3K4 hypermethylation, while dehydration and osmotic stress can also involve DNA methylation changes and histone-associated regulation (Brzezinka *et al*., 2016; Liu *et al*., 2018; Korotko *et al*., 2021; Friedrich *et al*., 2021; Pratx *et al*., 2023; Nishio *et al*., 2024; Rao *et al*., 2024; Behl *et al*., 2025, Preprint). These findings suggest that recurring abiotic stress can leave a regulatory imprint beyond the initial exposure period, although the extent to which such states are maintained or reset remains strongly context-dependent; recent studies emphasise that resilience depends on the balance between stress memory and resetting (Oberkofler *et al*., 2021; Staacke *et al*., 2025).

A hallmark of plant stress responses is the rapid production and signalling of reactive oxygen and nitrogen species (ROS and RNS, respectively) (Delledonne *et al*., 1998; Durner *et al*., 1998; Miller *et al*., 2009). These transient redox molecules transduce environmental cues into changes in cellular metabolism via post-translational modifications (PTMs) of proteins. Nitric oxide (NO), a pivotal RNS, has emerged as a modulator of the transcriptome and epigenetic landscape (Skelly *et al*., 2016; Lindermayr *et al*., 2020). NO mainly signals through protein *S-*nitrosation (protein-SNO), a post-translational modification that targets regulatory thiol groups in proteins. *S*-nitrosation of the master immune transcriptional co-activator NONEXPRESSOR OF PATHOGENESIS-RELATED GENES 1 (NPR1) regulates its subcellular localisation and interactions with transcription factors, thereby driving the expression of thousands of defence-related genes (Tada *et al*., 2008; Lindermayr *et al*., 2010). Conversely, nucleosome repositioning mediated by the chromatin remodelling ATPase EMBRYO SAC DEVELOPMENT ARREST 16 (EDA16) dampens immune responses by selectively repressing a transcriptional programme involved in the biosynthesis of the redox-active, immune activator glutathione (GSH) (Pardal *et al*., 2021). Taken together, these findings suggest redox signalling confers dynamic plasticity to transcriptional responses. GSH is a major cellular antioxidant that may react with NO to form the potent *trans*-nitrosylating agent, *S-* nitrosoglutathione (GSNO). Cellular GSNO pool is largely controlled by the evolutionarily conserved GSNO reductase 1 (GSNOR1) via the NADH-dependent reduction of GSNO to oxidized GSH and ammonium. GSNOR1-deficient *Arabidopsis* (*gsnor1-3,* hereafter *gsnor-ko*) plants exhibit overaccumulation of protein-SNO, resulting in reduced growth, delayed development, sensitivity to higher temperature and biotic stress as well as tolerance to oxidative stress (Feechan *et al*., 2005; Lee *et al*., 2008; Chen *et al.*, 2009, Holzmeister *et al*., 2011). This demonstrates that GSNOR1 indirectly controls NO signalling (Jensen *et al*., 1998; Liu *et al*., 2001). Our previous work demonstrated that *gsnor-ko* exhibits altered chromatin accessibility and, therefore, transcription of stress-responsive genes. Mechanistically, this epigenetic reprogramming is underpinned by the accumulation of the primary methyl donor for DNA and histone methylation, S-adenosylmethionine, and the inhibitory *S*-nitrosation of the histone modifier, HISTONE DEACETYLASE 6 (HDA6) (Ageeva-Kieferle et al., 2021; Rudolf et al., 2021; Lindermayr et al., 2010). It remains unknown, however, if GSNOR1 drives stress-specific transcriptional programmes across generations and affects subsequent transcriptional recovery.

Here, we used a five-generation Arabidopsis experiment to test how repeated exposure to future-climate scenarios shapes phenotypic and transcriptomic responses in wild type and the nitric oxide homeostasis mutant *gsnor-ko*. Specifically, we tested three linked hypotheses: First, repeated future-climate conditions would produce not only immediate phenotypic and transcriptional responses, but also multigenerational carryover effects, with the strongest shifts expected under the warming climate (hereafter, heat), CO_2_, ozone, drought, and combined (hereafter, combination) treatments. Second, if GSNOR-dependent NO/redox homeostasis contributes to post-stress recovery, WT should reset more efficiently toward a control-like transcriptomic state after stress withdrawal, whereas *gsnor-ko* should show broader persistence or reconfiguration of stress-responsive programs across generations. Third, if these differences reflect altered regulatory resetting rather than only stress severity, the persistent states detected in *gsnor-ko* should coincide with broader engagement of candidate chromatin- and RNA-associated pathways, including DNA methylation and RdDM-related regulation, heterochromatin maintenance, histone ubiquitination, RNA methylation, and methyl-donor metabolism. In this framework, NO/redox homeostasis is considered an upstream regulator of stress-state recovery versus persistence, while epigenetic pathways are treated as candidate mediators of altered recovery dynamics rather than as direct evidence of stable transgenerational epigenetic inheritance.

## Materials and methods

### 1. Plant material and cultivation

Seeds of *A. thaliana* ecotype Col-0 (WT) were obtained from the Nottingham Arabidopsis Stock Centre (NASC). The *gsnor1-3* mutant line (also reported as *hot5-2*; GABI-Kat line 315D11) was obtained from GABI-Kat (Rudolf *et al*., 2021). Both WT and *gsnor-ko* plants were grown in soil (Floradur® Anzucht Substrat, Floragard, Germany; substrate 0.5 kg/m3 NPK fertilizer (18-10-20) and trace elements) mixed with quartz sand (0.6 to 1.2mm) (4:1) in individual standard Photon Systems Instruments (Photon Systems Instruments-PSI Brno, Czechia) pots (6cm × 6cm × 9.5cm), containing 170g soil mix. These pots were placed in trays (5 × 4 pots per tray) and registered into the PSI PlantScreen™ system. After stratification for 2–3 d at 4°C in darkness, pots were transferred to the ExpoSCREEN chambers and cultivated under long-day conditions (14 h light / 10 h dark) at a photosynthetic photon flux density (PPFD) of 300 µmol m⁻² s⁻¹. Leaf tissue was harvested from 4- to 5-week-old rosettes at 3 h (from 9:00 to 10:00) after lights-on, flash-frozen in liquid nitrogen, and stored at −70 °C until extraction.

### 2. Multi-generational experimental design

#### 2.1 Overview and generation scheme

Plants were propagated in a multi-generational experiment as described in Fig. 1A and Supplementary Fig. S1A. Seeds originating from a single WT and a single *gsnor-ko* plant (generation G0) were used to produce G1 offspring (approximately 3 months for each generation) (Rudolf *et al*., 2021). Offspring were randomly assigned to climate scenarios (approximately 40 plants per condition per genotype). For each genotype × condition, 10 plants were randomly selected and used for seed production and sequencing (two-leaf harvest), 10 for phenotyping, and the remaining ∼20 for a 4-week post-germination harvest (the same time of day as for RNA sampling) for downstream analyses.

**Fig 1.**
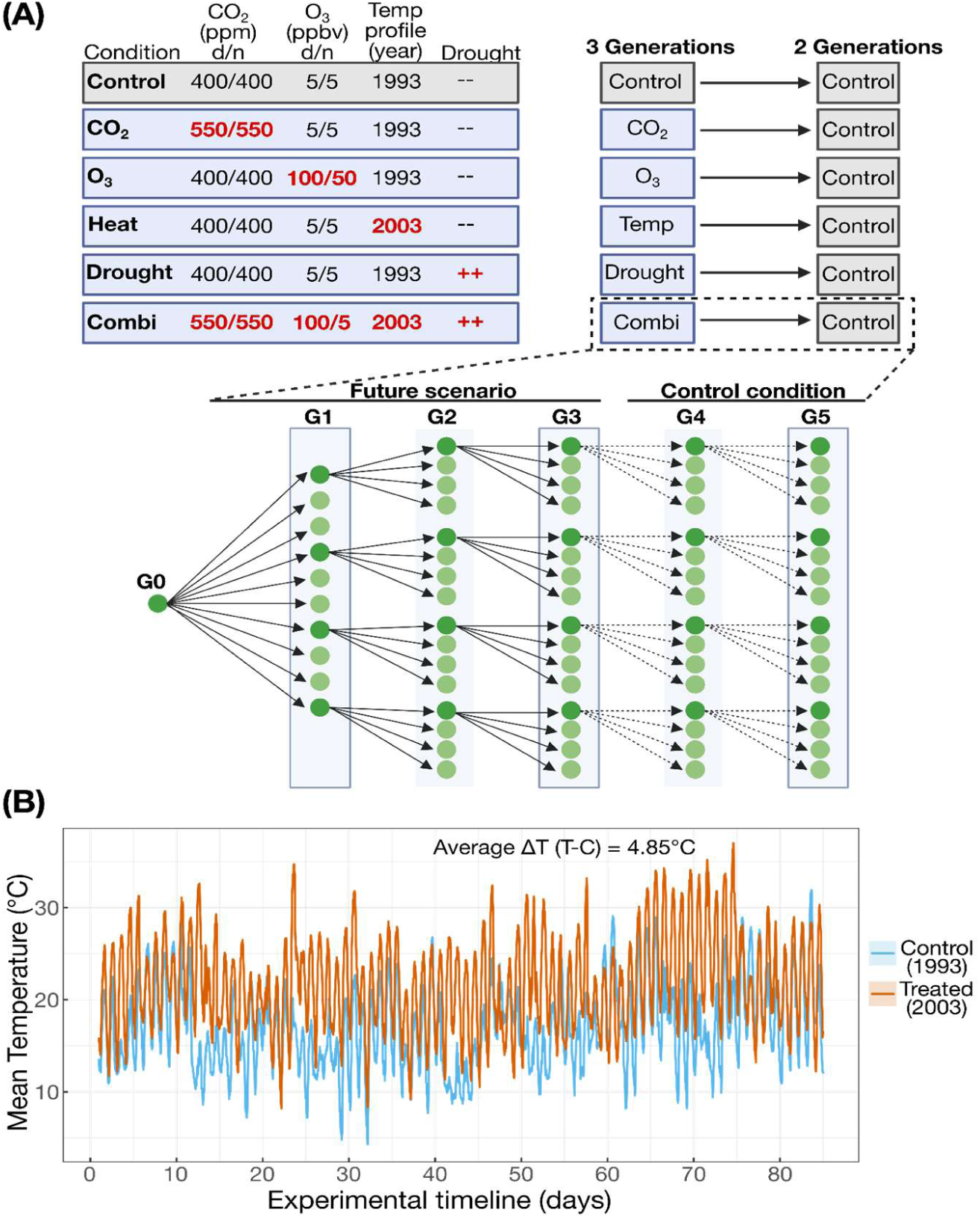
Experimental design of the multi-generational future-climate study. **(A)** Wild-type *Arabidopsis thaliana* Col-0 (WT) and the nitric oxide homeostasis mutant *gsnor1-3* (*gsnor-ko*) were propagated across five successive generations (G1-G5) starting from a founder generation (G0). Plants were grown under six climate scenarios: control, elevated CO₂, elevated O₃, warming, drought, and a combined treatment (Combi). Treatments were applied for three consecutive generations (G1-G3), followed by two generations grown under control conditions (G4-G5) to assess the persistence or recovery of stress responses. The environmental regimes for each condition are summarised, including day/night CO₂ and O₃ concentrations, the annual temperature profile, and drought treatment status. **(B)** Temperature profiles used for the ExpoSCREEN simulations. Control conditions followed the 1993 temperature profile, whereas the treated warm scenario followed the 2003 profile. The treated profile was, on average, ∼5°C warmer than the control profile over the experimental timeline.

To enable pedigree-based analyses, offspring were randomly assigned to one of four pedigrees, and each individual progenitor plant generated the subsequent generation (approximately 20 offspring per progenitor). For molecular analyses, a random subset of siblings was sampled as described below. To minimise positional effects, pots were redistributed within chambers every 10-12 days using a random placement scheme. Treatments were run in parallel chambers within each generation, and generations were grown sequentially (i.e., one generation at a time) with the same treatment programs applied in each generation.

#### 2.2 Climate scenarios/treatments

Temperature regimes were designed to mimic naturally occurring local seasonal conditions for the Munich region, based on per-minute 2 m air-temperature records monitored at the Deutscher Wetterdienst (DWD) station München-Flughafen (station ID 01262; 48°22′ N, 11°48′ E; 453 m a.s.l.), downloaded from the DWD Climate Data Centre archive (Supplementary File S1) (Wetterdienst, 2026). For control conditions, a slightly modified per-minute temperature profile derived from June to August 1993 was implemented (monthly mean temperatures: 16.5°C, 16.1°C, and 17.1°C, respectively). For the warm-temperature simulation, per-minute temperature profiles derived from June–August 2003 were used (monthly mean temperatures: 21.5°C, 20.4°C, and 22.4°C, respectively), reflecting a historically hot summer and intended to approximate conditions projected to become typical near the end of the century, roughly the 2080s, under high-emissions scenarios in Bavaria (Bayerisches Staatsministerium für Umwelt und Verbraucherschutz, 2021) (Fig. 1B; Supplementary File S1). To avoid confounding effects from the temperature treatment, the temperature during a few days in the 1993 climate program was consistently lower than in the 2003 program throughout the climate simulation. The final mean temperature differences between the warm and control programs were approximately +5.0°C (June), +4.3°C (July), and +5.3°C (August) across the temperature regimes (Fig. 1B; Supplementary Fig. S2A).

Under control conditions of the 1993 simulation scenario, atmospheric CO_2_ concentrations were 400/400 ppm (day/night), and O_3 was_ ∼5/5 ppbv (day/night; also negligible, as it was scrubbed by charcoal filters in the chambers) and plants were well-watered. Future climate scenarios included enhanced CO_2_ levels of 550/550 ppm (day/night), a concentration projected to be reached approximately by 2050 under SSP5-8.5 and by 2070 under SSP2-4.5, depending on the climate scenario (Meinshausen, 2020). Plants were exposed to 100 ppb O_3_ during the day and 50 ppb O_3_ at night to mimic an elevated summer urban/peri-urban ozone regime rather than background conditions (Fitzky *et al*., 2019; Im *et al*., 2013). Drought stress was applied by reducing soil water content to approximately 50% of the control value (Fig. 1; Supplementary Fig. S2 and S3; Supplementary Files S3-S5). Except for drought, treatments were initiated 1 week after germination; drought was imposed from the start (see Section 3.2).

### 3. Climate chamber settings and environmental monitoring

#### 3.1 Baseline growth conditions

Simulation of different environmental conditions was performed in the ExpoSCREEN chambers at Helmholtz Munich (HMGU; as described in Vanzo *et al*., 2015; Roy *et al*., 2016; Ghirardo *et al*., 2020; Roy *et al*., 2021). Each chamber contained four subchambers made of acrylic glass (about 1 m^3^ in volume). In each subchamber, different treatments were applied, such as elevated CO_2_, high O_3_, and drought. In another large chamber (containing 4 subchambers), heat and combined conditions were applied. Both WT and *gsnor-ko* plants were grown together in each subchamber. Each subchamber was equipped with combined air temperature and relative humidity sensors (ifm, Switzerland) and was flushed (80 m^3^ h^−1^) with purified air (charcoal-filtered) to adjust temperature and humidity for each ExpoSCREEN chamber, providing the same temperature and humidity in all subchambers of the respective ‘main’ chamber. To achieve irradiation regimes very close to outdoor solar conditions, from UV to near-infrared, the phytotron facility used a combination of lamps and filters that simulate the daily course of solar radiation from sunrise to sunset (Thiel *et al*., 1996). Details about climate conditions in subchambers are reported in the Supplementary Files S2-S7. Plants were watered manually by adding water directly to the trays. Watering decisions were based on pot weight and chamber conditions (temperature program) (Supplementary Files S2, S6, and S7). PPFD was set to 300 µmol m⁻² s⁻¹ at plant height (Supplementary File S4). Plants were fertilised with Compo Complete Pflanzendünger (liquid fertiliser, Brand Compo, EAN 4008398135737, 3ml dissolved in 1 litre of water) at the end of weeks 3 and 4. Due to technical issues with the chamber sensors, temperature, humidity, and fumigation data were not recorded for 13 days for G1 (control, CO_2_, drought, and O_3_) and for 3 days for G4 (Supplementary Fig. S2 and S3; Supplementary Files S2-S5).

#### 3.2 Drought treatment

Drought was imposed from the start of the experiment by allowing water to evaporate from pre-watered pots until they reached a weight of 220 g (pot + soil + water), corresponding to an estimated Soil Water Content (SWC, vol-%) of approximately 23%vol. SWC was measured using a soil moisture meter (HH2) connected to a ThetaProbe (ML2x; Delta-T Devices, Britain, UK). After sowing, drought pots were watered ∼45–50% less than control plants. Heat-treated plants were watered ∼45-50% more than control plants, and combination-treated plants were watered ∼50% less than heat-treated plants and, in some cases, were similar to controls, as combination plants also experienced heat treatment. This was done to compensate for the higher VPD and evaporative demand under a warmer climate and to prevent plants from lethal drying (Supplementary Fig. S3 and Supplementary Files S6-S7).

### 4. Experimental timeline, sampling strategy and statistical analysis

Plants were phenotyped using a PSI PlantScreen™ platform (Photosystem Instruments Brno, Czechia) at the end of week 3 and week 4, including morphological traits (e.g., rosette area) and chlorophyll fluorescence parameters (e.g., Maximum Quantum Yield of Photosystem II - QYmax and Non-Photochemical Quenching - NPQ_Lss). QYmax (Fv/Fm) was used as an estimate of the maximum efficiency of Photosystem II (PSII) photochemistry in dark-adapted tissue, whereas NPQ_Lss was used as a measure of non-photochemical quenching at light steady phase and photoprotective heat dissipation during illumination (Baker, 2008; Murchie and Lawson, 2013).

Exported PSI data were analysed in R (using packages car, emmeans, multcomp, and tidyverse) (Supplementary Files S8-S12). Phenotypic traits were analysed using Generalised Linear Models (GLM) with a Gaussian error distribution. For week 3 and week 4 PSI measurements, genotype, treatment, and generation were included as fixed factors together with their interactions. Overall model effects were evaluated by Type III analysis of variance, and estimated marginal means were calculated for post hoc comparisons. Pairwise contrasts were used to compare genotypes within treatment × generation combinations and treatments within genotype × generation combinations, with p-values adjusted by the Benjamini–Hochberg method. For treatment-specific plots, separate Gaussian GLMs were fitted within each treatment using genotype, generation, and their interaction, followed by Tukey-adjusted post hoc tests and compact letter displays. Descriptive statistics are reported as mean ± SE. No additional dark-adaptation time was needed for the fluorescence measurements in the PSI system, as plants were transferred to the PSI platform the evening before measurement and measured the next morning, before lights-on. However, due to an error in the PSI software, plant trays in G3 (all treatments at week 4) and G4 (drought and heat at week 4) were re-measured after a 30-minute dark-adaptation period.

For sequencing, two random fully elongated (mature) leaves per individual plant were harvested at week 4.5 (3 h after lights-on; 9:00-10:00). Leaves for RNA-seq were harvested individually for each plant and not pooled. Seeds from each plant were collected and weighed to quantify reproductive output across treatments. They were harvested at approximately week 10 (WT) and week 11 (*gsnor-ko*), corresponding to comparable reproductive/developmental stages rather than identical chronological ages, as the mutant generally developed more slowly than WT. At harvest, siliques had turned brown/dry, and seeds were fully matured. Seeds were collected per individual plant (and were not pooled) and stored at 4°C (Supplementary Fig. S1A; Supplementary File S10). Additionally, seed phenotyping was performed using the phenoSeeder platform to complement seed weight measurements (Jahnke *et al*., 2016; Klasen *et al*., 2025). Individual WT seeds were analysed under standardised conditions to compare seed trait distributions across treatments and generations. Because *gsnor-ko* produced insufficient seeds in several treatment × generation combinations, distribution-based seed phenotyping was restricted to WT, while seed weight per plant was quantified in both genotypes. This approach enabled assessment of both total reproductive output and variation in individual seed phenotypes within seed batches. Seed weight and mass were analysed analogously, with separate Gaussian GLMs fitted for each treatment using genotype, generation, and their interaction as explanatory variables (as described above for PSI data).

### 5. RNA extraction, library preparation, and sequencing

RNA-seq was performed from snap-frozen leaves (two leaves per plant). For each genotype and condition, five biological replicates were analysed for controls (except for G1 WT Control, where 6 biological replicates were used) and four biological replicates for treated samples (drought, CO_2_, O_3_, heat, and combination).

Total RNA was extracted from ∼50 mg ground tissue using an in-house protocol based on the Sigma TRI Reagent method (Chomczynski and Sacchi, 1987). RNA quantity and integrity were assessed using a Nanodrop and a Bioanalyzer.

Sequencing libraries were prepared by BMK Gene GmbH (Münster, Germany) using poly(A)-enriched RNA and sequenced on a DNBSEQ-T7 platform as 150 bp paired-end reads, generating ∼9 GB per sample (approximately 30 million reads per sample) (https://www.bmkgene.com/eukaryotic-mrna-sequencing-illumina-product/). Reads were mapped to the *A. thaliana* TAIR10 reference genome using HISAT2 (v2.2.1; https://daehwankimlab.github.io/hisat2/main/). Read counting was performed with featureCounts using TAIR10 annotation files (access date-19 June 2023) (Liao *et al*., 2013). Differential gene expression analysis was conducted in R using DESeq2 (DESeq2_1.42.1) (Supplementary Fig. S1B). All the calculations and visualizations were carried out using R programming and Rstudio, with plot adjustments performed in Inkscape.

### 6. Pathogen infection

For disease assays, plants were grown in soil in a controlled environment chamber at 21°C and 100 µmol m-2 s-1 light, 50-65% relative humidity, and a photoperiod of 16/18 h light/dark. *Pseudomonas syringae* pv. maculicola (Psm) ES4326 were cultivated overnight in LB medium containing 10 mM MgSO_4_ and kanamycin (50µg/mL) at 28°C, harvested by centrifugation and resuspended to OD600 = 0.0005 in 10 mM MgSO_4_. Leaves of comparable age and size were infiltrated with the bacterial suspension on the abaxial surface using a 1 mL syringe. For each experimental condition, 16–18 leaves were infiltrated (≤2 per plant), and plants were maintained in the growth chambers for 5 days after inoculation. Leaf extracts were serially diluted in 10 mM MgSO_4_ and streaked in parallel on LB agar supplemented with kanamycin. After incubation at 28°C for 2 days, colonies were counted and bacterial titre calculated per leaf disc (Frungillo *et al*., 2023, Preprint).

### 7. AI usage in Data analysis

For data analysis, AI tool (ChatGPT 5.4) was used for programming in R (2024.12.1+563). After using this tool/service, the author(s) reviewed and edited the codes as needed and take(s) full responsibility for the content of the published article.

## Results

*Arabidopsis thaliana* WT and the NO-homeostasis mutant *gsnor-ko* were propagated for five successive generations in controlled-climate chambers under current and projected future climate scenarios (Fig. 1; Supplementary Fig. S1A; see Materials and Methods).

### 1. Phenotypic output in response to climate scenarios are genotype and generation specific

#### 1.1 Rosette growth responses are stress- and generation-dependent

Using the PSI platform, phenotyping of plants was performed at week 3 and week 4 to measure growth parameters. At week 4, WT rosette area varied strongly with both treatment and generation (Fig. 2A-B). The largest WT rosettes were observed under warmer climates, especially in the treated generations, with a peak in G2 (4842.5 mm^2^), followed by a decline in later generations (2207.0 mm^2^ in G3, 1559.9 mm^2^ in G4, and 1309.5 mm^2^ in G5). A similar but smaller transient increase was observed under the combination treatment, where WT exhibited the highest rosette area in G2 (3097.1 mm²). In contrast, the rosette area was lower in G3 (1611.1 mm²), G4 (1591.5 mm²), and G5 (1390.4 mm²), and these later generations did not differ significantly from one another. Under drought, WT rosette area increased from 1217.0 mm^2^ in G1 to 2143.0 mm^2^ in G2, decreased again in G3-G4, and partially recovered in G5 (1816.5 mm^2^). O_3_-treated WT plants showed a marked reduction in G3 (618.2 mm^2^) relative to G1-G2, followed by recovery in G4-G5 (1517.1-1392.6 mm^2^). Elevated CO_2_ also showed a pronounced G2 increase (2344.7 mm^2^), followed by a reduction in G3 (783.3 mm^2^) and partial recovery in G4-G5 (1312.1-1554.6 mm^2^) (Fig. 2A-B; Supplementary File S13).

**Fig 2.**
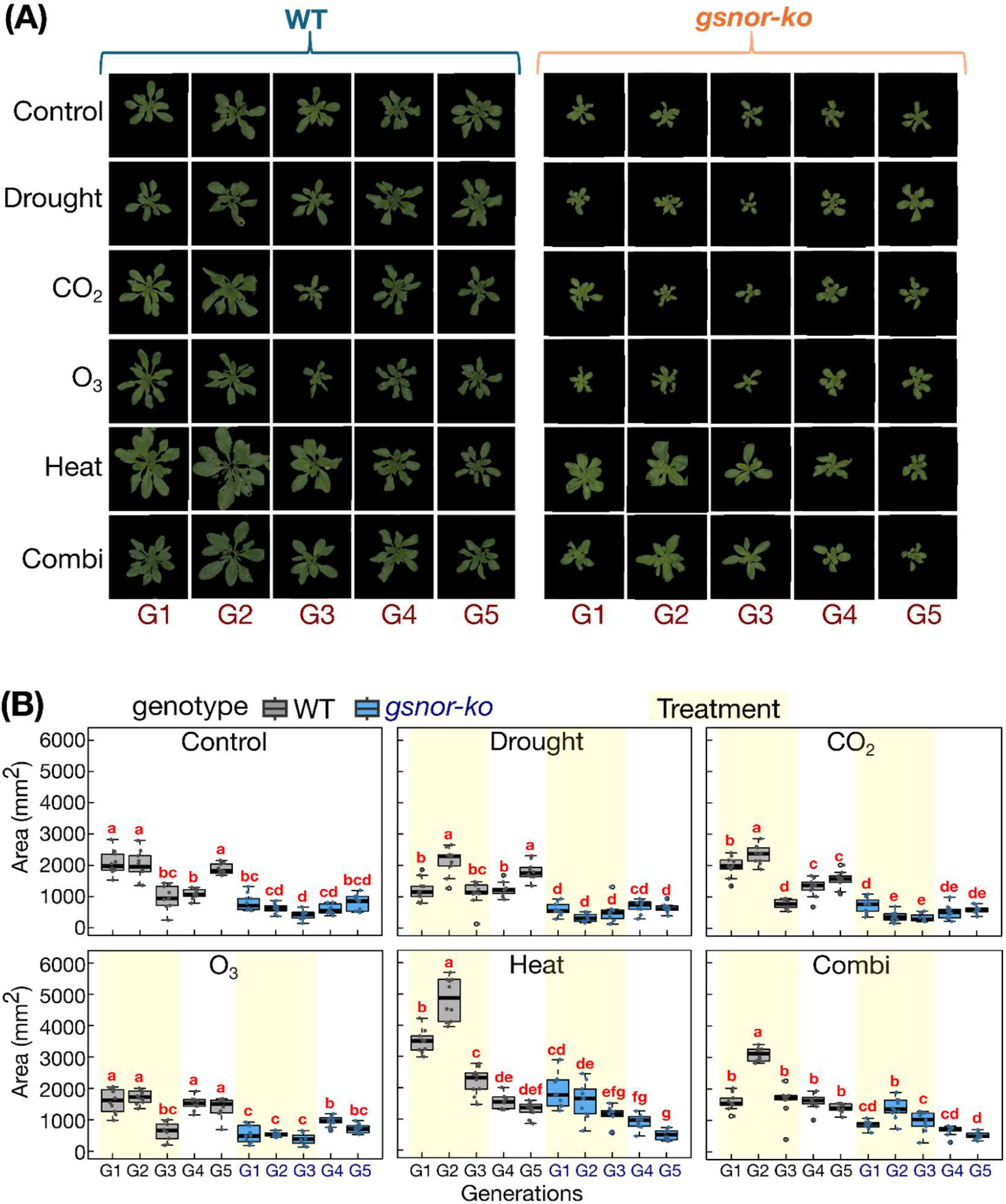
Multi-generational phenotypic responses of WT and *gsnor-ko* at week 4. **(A)** Representative rosette images of WT and *gsnor-ko* plants grown under Control, Drought, elevated CO_2_, elevated O_3_, Heat, and Combination treatments across generations G1-G5. **(B)** Rosette area at week 4 for WT and *gsnor-ko* across treatments and generations. Boxplots show the distribution of individual values, with points representing biological replicates. Beige shading indicates generations exposed to stress treatment; G4 and G5 were grown under control conditions after stress withdrawal. Different letters indicate significant differences among genotype × generation groups within each treatment (GLM followed by post hoc multiple comparison, *P* < 0.05).

Across all week-4 environments, *gsnor-ko* plants remained substantially smaller than WT. Under control conditions, for example, WT reached 2072.3 mm^2^ in G1 and 2000.7 mm^2^ in G2, whereas *gsnor-ko* reached only 807.2 mm^2^ and 632.4 mm^2^, respectively. The difference was especially pronounced under heat in G2, where WT reached 4842.5 mm^2^ compared with 1599.6 mm^2^ in *gsnor-ko* (adjusted p < 0.001). A similar contrast was observed under the combination treatment in G2 (3097.1 mm^2^ in WT versus 1377.2 mm^2^ in *gsnor-ko*, adjusted p < 0.001). Even after stress removal, descendants of the *gsnor-ko* line remained smaller than WT; under the combination lineage in G5, WT averaged 1390.4 mm^2^ whereas *gsnor-ko* averaged 513.1 mm^2^ (adjusted p < 0.001), and under heat in G5, WT averaged 1309.5 mm^2^ versus 492.5 mm^2^ in *gsnor-ko* (adjusted p < 0.001) (Supplementary Files S12-S13). Overall, WT lineages often showed partial recovery across generations, whereas *gsnor-ko* showed lower growth capacity and weaker recovery across most stress histories.

To distinguish absolute growth differences from treatment-specific resilience, rosette area was additionally normalized to the corresponding control within each genotype and generation (Supplementary Fig. S8). This analysis showed that some WT stress lineages performed closer to their generation-matched controls than was apparent from the absolute leaf-area plots. In particular, WT drought remained near control-equivalent levels across later generations, whereas heat- and combination-derived lineages showed elevated relative performance through G1–G4 but declined again by G5. In *gsnor-ko*, normalized responses also varied across generations, but the mutant remained strongly growth-limited in absolute terms, indicating that relative within-genotype resilience did not translate into WT-like rosette size.

The same overall pattern was already evident at week 3 (Supplementary Fig. S9), although absolute rosette sizes were smaller. For example, under heat in G2, WT plants averaged 1654.3 mm^2^ whereas *gsnor-ko* averaged 618.2 mm^2^ (adjusted p < 0.001), and under the combination treatment in G2, WT reached 1177.7 mm^2^ compared with 667.6 mm^2^ in *gsnor-ko* (adjusted p < 0.001) (Supplementary Files S14-S15).

#### 1.2 Photosynthetic performance is comparatively stable, with genotype- and generation-specific shifts

In contrast to the rosette area, QYmax varied less across treatments and generations (Supplementary Fig. S10-S11). At week-4, genotype, treatment, and generation effects were statistically significant (adjusted p < 0.001), but the fitted means remained within a relatively narrow range, from 0.737 to 0.852 across all genotype × treatment × generation combinations. The highest week-4 QYmax value was observed in WT under the combination treatment in G2 (0.852), whereas the lowest was observed in *gsnor-ko* under control conditions in G5 (0.737). Thus, the pronounced growth penalty in *gsnor-ko* was not associated with a broad collapse of maximum PSII efficiency (Supplementary Files S12-S15).

By contrast, NPQ_Lss showed stronger condition- and generation-dependent variation than QYmax (Supplementary Fig. S12-S13). At week 4, NPQ_Lss was significantly affected by genotype, treatment, and generation (adjusted p < 0.001). Across week-4 means, NPQ_Lss ranged from 1.12 to 1.71, indicating greater relative variation than QYmax. In G5, *gsnor-ko* also showed altered energy dissipation relative to WT (Supplementary Files S12-S15). Together, these results indicate that differences between WT and *gsnor-ko* are expressed more strongly in growth and photoprotective regulation than in maximum PSII quantum efficiency.

#### 1.3 Reproductive output across climate scenarios in *gsnor-ko* and WT

Seed weight per plant was markedly lower in *gsnor-ko* than in WT across all treatments and generations (Supplementary Fig. S14). Depending on treatment and generation, mutant seed yield was typically about 7-27% of WT values. Under control conditions in G1, for example, WT plants produced 642.6 mg seeds per plant compared with 91.2 mg in *gsnor-ko*. Under the combination treatment in G5, WT reached 595.3 mg whereas *gsnor-ko* reached only 70.1 mg; under heat in G5, WT produced 515.0 mg compared with 70.9 mg in *gsnor-ko* (Supplementary File S16).

In WT, seed production varied strongly across generations within each treatment. Under drought, WT seed weight decreased from 618.0 mg in G1 to 195.5 mg in G2, then increased to 243.1 mg in G3, and recovered to 475.4 mg in G4 and 527.4 mg in G5 (adjusted p < 0.001). Under CO_2_, WT seed weight was highest in G1 (753.3 mg), decreased in G2 and was comparable in G2-G3 (388.6 and 327.1 mg), and partially recovered in G4-G5 (482.5 and 477.8 mg). Under O_3_ and heat, WT again showed strong generation-dependent shifts, with lower seed yield in intermediate generations and recovery under control conditions in G4 and G5. Under the combination treatment, WT seed weight was lowest in G2-G3 (208.9 and 191.7 mg) but increased sharply after stress removal, reaching 535.3 mg in G4 and 595.3 mg in G5 (adjusted p <0.001) (Supplementary Fig. S14A; Supplementary Files S16-S17).

In addition to seed yield, phenoSeeder-based seed phenotyping showed that repeated climate exposure altered the distribution of seed traits in WT. Compared with controls, heat and combined treatments produced broader distributions of seed traits and/or shifts in their distributions across generations (Supplementary Fig. S14B-C; Supplementary Files S18-S19). Thus, repeated climate exposure influenced both the quantity and heterogeneity of seeds produced.

By contrast, *gsnor-ko* seed weight remained consistently low across all generations and showed significantly weaker generational modulation. Across all treatments, no generation-to-generation contrast within *gsnor-ko* reached significance in this analysis, indicating that mutant reproductive output remained uniformly reduced regardless of stress history (Supplementary Fig. S14; Supplementary Files S16-S17). Thus, whereas WT often showed substantial recovery of seed production in G4 and G5 (treatment history lineages), particularly under heat, drought, and combination-exposed plants (G1-G3), *gsnor-ko* maintained persistently low reproductive output throughout the experiment.

### 2. Pattern of differential expression across climate scenarios and generations

#### 2.1 Wild type exhibits scenario- and generation-dependent DEG responses

Differential expression analysis revealed condition-specific transcriptional responses in WT across the multigenerational experiment (Fig. 3A & 3B). The number of DEGs varied substantially among climate scenarios and generations (adjusted p < 0.05), ranging from only 4 genes in the weakest contrasts to approximately 6000 in the strongest. The strongest response was detected in G1 under the combination scenario, whereas the weakest response was observed in G1 under elevated CO_2_. The relative proportions of up- and down-regulated DEGs also differed among scenarios, indicating that each climate condition elicited a distinct transcriptional program.

**Fig 3.**
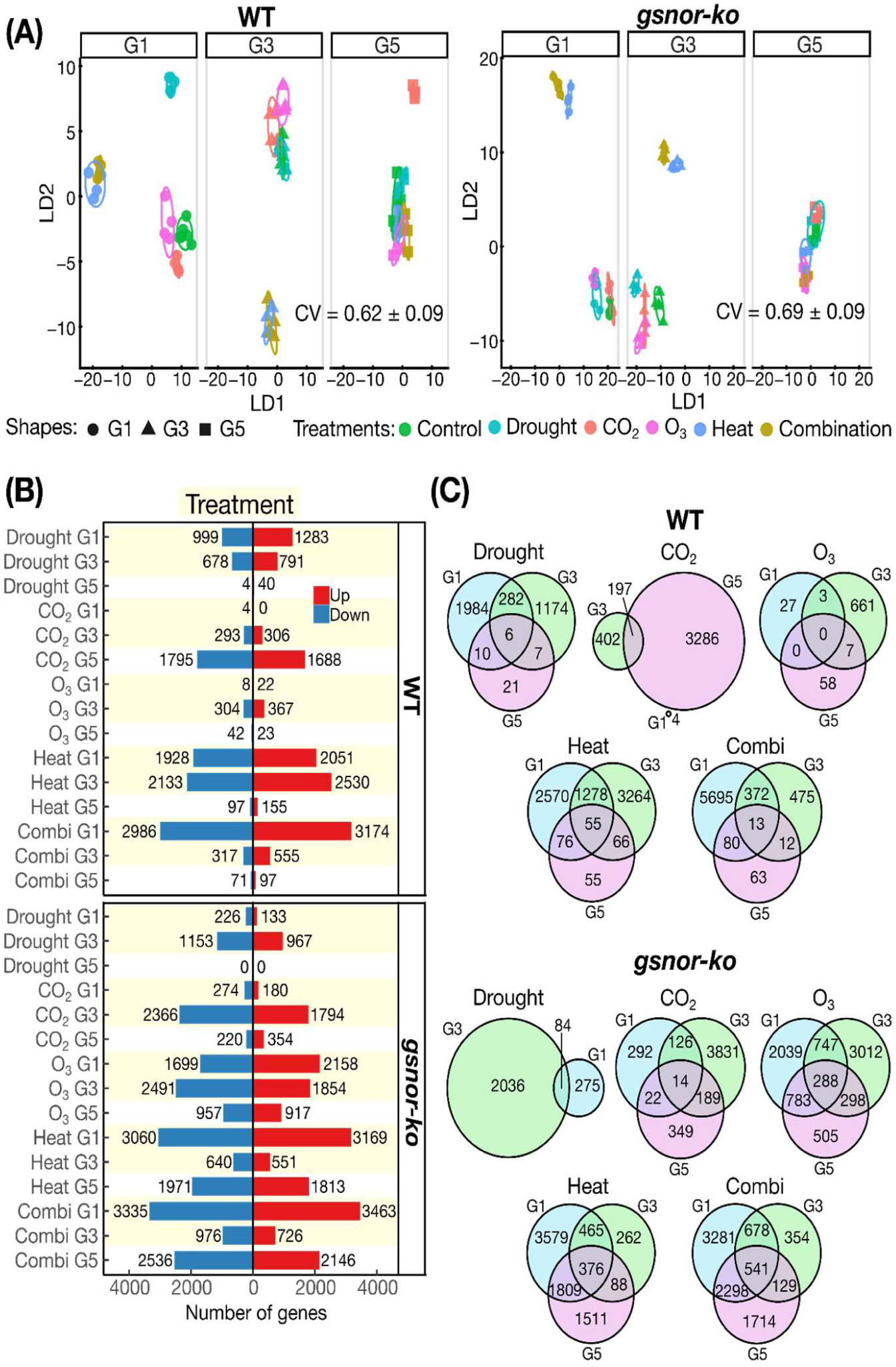
Global transcriptome structure, differential expression, and intergenerational overlap in WT and *gsnor-ko*. **(A)** Discriminant analysis of principal components (DAPC) of VST-transformed RNA-seq data for WT and *gsnor-ko* across generations G1, G3, and G5. Symbols indicate generations and colours indicate treatments (Control, Drought, CO_2_, O_3_, Heat, and Combination). LD1 and LD2 represent the first two discriminant axes. CV values indicate cross-validation performance. **(B)** Numbers of up-regulated and down-regulated differentially expressed genes (DEGs) for each treatment and generation in WT and *gsnor-ko*. Up-regulated genes are shown in red and down-regulated genes in blue. **(C)** Venn diagrams showing overlap of DEG sets among G1, G3, and G5 within each treatment for WT and *gsnor-ko*. Numbers indicate the number of unique and shared DEGs across generations. Combi, combination treatment.

Across generations, drought and the combination scenario showed a general decline in DEG numbers from G1 to G3 and then from G3 to G5, suggesting attenuation of the transcriptional response or reduced responsiveness after stress removal. By contrast, under elevated CO_2_ and heat DEG numbers increased from G1 to G3 and from G3 to G5. Responses to elevated O_3_ were intermediate, with relatively few DEGs in G1, a stronger response in G3, and a subsequent decline by G5-treatment history lineage (Fig. 3B; Supplementary Fig. S15A).

#### 2.2 *gsnor-ko* shows altered DEG magnitude and directionality

In *gsnor-ko*, both the numbers and direction (up- or down-regulation) of DEGs differed from WT across environmental conditions and generations (Fig. 3B). DEG numbers ranged from about 130 to 3500 (adjusted p < 0.05), with heat and the combination scenario eliciting the strongest responses, whereas drought generally induced fewer changes. Relative to WT, *gsnor-ko* showed weaker responses under drought-history (in G5) and elevated CO_2_-history (in G5), but stronger responses under elevated CO_2_ (in G3), elevated O_3_ (in G1, G3, and G5), heat (in G1 and G5-treatment history), and the combination scenario (in G1, G3 and G5-treatment history), consistent with altered transcriptional responsiveness under impaired NO homeostasis.

Under drought, CO_2_, and O_3_, *gsnor-ko* showed its strongest DEG response in G3 relative to G1 and G5-treatment history. By contrast, under heat and the combination scenario, DEG numbers were highest in G1, decreased in G3, and increased again in G5-treatment history (Fig. 3B; Supplementary Fig. S15B). These patterns indicate that impaired NO homeostasis changes not only DEG magnitude but also the temporal profile of transcriptional responses across generations.

To assess intergenerational consistency, we compared DEG sets within each environmental scenario across generations and, separately, within each generation across factors (Fig. 3C; Supplementary Fig. S16A-B). Only a subset of DEGs overlapped across generations within each climate scenario. In WT, the heat scenario showed the greatest overlap across G1, G3, and G5 (55 shared DEGs), whereas elevated O_3_ showed no overlap. In *gsnor-ko*, the combination scenario showed the highest overlap (541 shared DEGs), whereas drought (0) and elevated CO_2_ (14) showed the lowest overlap among the three analyzed generations. Overall, intergenerational DEG overlap was greater in *gsnor-ko* than in WT, consistent with a broader persistent core response in the mutant background.

### 3. Functional signatures identify chromatin- and RNA-linked regulation as candidate layers of the multi-generational stress response

#### 3.1 Enrichment analysis identifies recurrent regulatory themes across treatments and generations

To interpret stress-responsive transcriptome changes, we performed functional enrichment analysis on up- and down-regulated gene sets for each treatment x generation x genotype combination and summarised the results using Mercator categories. To emphasise reproducible biology rather than isolated contrasts, we further grouped enriched terms into shared between genotypes and epigenetic-associated categories and visualised significant hits across the dataset (Fig. 4). Recurrent functional signatures extended beyond metabolism and photosynthesis to include categories linked to chromatin organisation, histone-related structure, RNA processing, and regulatory reprogramming.

**Fig 4.**
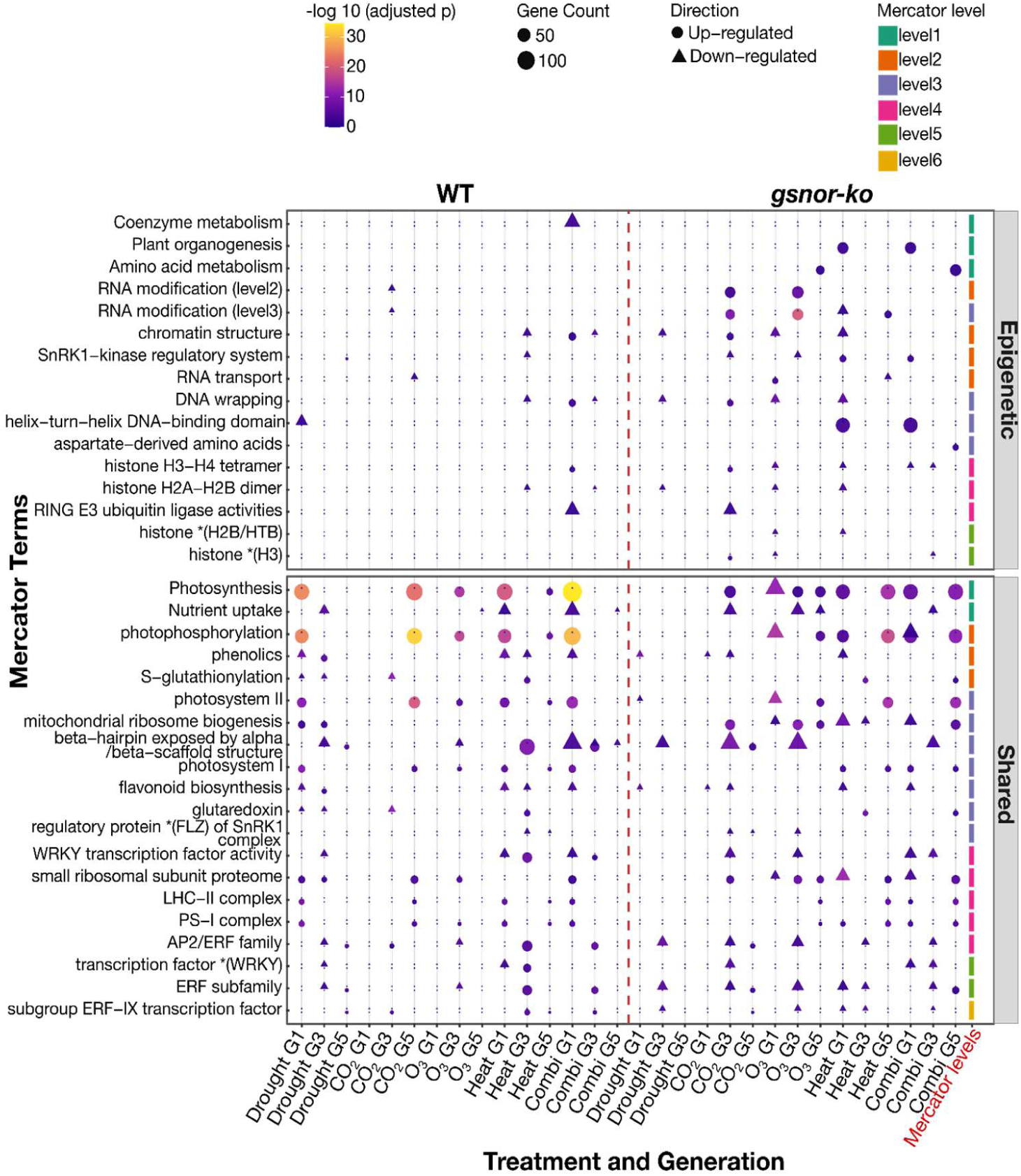
Recurrent functional enrichment signatures across treatments and generations in WT and *gsnor-ko*. Bubble plot summarising filtered Mercator enrichment terms identified across treatment × generation contrasts in WT and *gsnor-ko*. The upper panel shows epigenetic- and chromatin-associated categories; the lower panel shows recurrent shared functional categories. Circle size indicates the number of genes assigned to each term, and colour indicates enrichment significance, as -log_10_-adjusted *P*-value. Circles denote enrichment among up-regulated genes and triangles denote enrichment among down-regulated genes. The coloured bar at the right indicates Mercator levels 1–6.

Among the terms shared across WT and *gsnor-ko*, recurrent enrichment was observed for photosynthesis, photophosphorylation, photosystem I/II, glutaredoxin, S-glutathionylation, nutrient uptake, FLZ/SnRK1-related regulation, pre-RNA splicing, and several stress-associated transcription-factor families. These shared categories indicate that both genotypes engage overlapping energy, redox, and regulatory programs during multi-generational stress exposure.

#### 3.2 Explicit chromatin- and RNA-linked enrichment terms support a mechanistic regulatory hypothesis

The enrichment analysis also recovered a distinct set of terms directly relevant to chromatin- and RNA-associated regulation. These included histone H3, histone H2B/HTB, histone H2A-H2B dimer, histone H3-H4 tetramer, DNA wrapping, chromatin structure, RNA modification, RNA transport, and RING E3 ubiquitin ligase and CULLIN-based E3 ubiquitin ligase activities. Together, these categories indicate that the stress-responsive transcriptome repeatedly intersects with pathways involved in nucleosome composition, chromatin packaging, post-transcriptional regulation, and protein turnover.

The genotype-specific (’Different’) enrichment panel further showed that these regulatory signatures were not equally distributed between genotypes. In particular, *gsnor-ko* displayed broader and denser enrichment across multiple regulatory categories, including chromatin structure, RNA modification, DNA damage response, RNA editing, proteasome-associated terms, and additional redox- and signalling-related functions, especially under O_3_, heat, and combination conditions. In contrast, WT responses were generally more restricted. This broader regulatory footprint in *gsnor-ko* is consistent with the DEG-overlap and state-flow analyses, which point to reduced transcriptomic resetting in the redox mutant after stress removal (Fig. 4; Supplementary Fig. S17-S18).

Taken together, the enrichment analyses indicate that multi-generational stress responses in this system involve not only metabolic and photosynthetic remodelling but also repeated engagement of chromatin-, histone-, RNA-, and redox-linked regulatory functions. We therefore next examined candidate genes with known or proposed roles in DNA methylation, heterochromatin maintenance, Polycomb-related repression, histone ubiquitination, RNA methylation, and methyl-donor metabolism.

### 4. Persistence and turnover of environmental factor-responsive genes across generations

#### 4.1 Gene-state flow reveals fading, emergent, persistent, and switcher behaviour

To move beyond DEG overlap and trace how transcriptional states changed across generations, each gene within each treatment was classified according to its log fold change relative to the corresponding control as up (log2 fold change > 0), down (log_2_ fold change < 0), or none (not differentially expressed in that generation) across G1, G3, and G5. These state transitions were visualised with alluvial plots (Supplementary Fig. S18). This framework partitions genes into four intuitive response modes: (i) persistent, in which the same differential-expression state is maintained across generations; (ii) fading, in which genes are differentially expressed under stress in earlier generations but return to a non-differential expressed state after transferring to control conditions (G5); (iii) emergent, in which no differential expression is detected in early generations but appears later, often in G5; and (iv) switcher behaviour, in which genes change direction between up- and down-regulation across generations.

In WT, the dominant trajectory under drought, heat, and the combination scenario was fading, with a substantial fraction of DEGs detected in G1 and G3 returning to a non-differentially expressed state when exposed to control conditions (G5). This pattern is consistent with broad transcriptomic resetting after stress removal in G4. By contrast, WT under elevated CO_2_ displayed a pronounced emergent component toward G5, consistent with the increase in DEG numbers during the recovery phase. WT under elevated O_3_ showed only a weak persistent signal, indicating limited long-term retention of ozone-responsive transcriptional states (Supplementary Fig. S18).

In *gsnor-ko*, O_3_, heat and combined conditions showed larger fractions of persistent and/or switcher trajectories than WT, most notably under the combination, heat, and O_3_ scenarios, in agreement with the higher intergenerational DEG overlap (Fig. 3; Supplementary Fig. S18). By contrast, *gsnor-ko* plants exposed to drought were dominated by fading trajectories and showed little evidence of maintained differential expression after two generations under control conditions. Overall, these state-flow patterns indicate that the mutant does not simply amplify stress responses globally; instead, it shows stress-specific differences in the retention, reversal, or delayed reconfiguration of transcriptional states across generations.

To validate the state transitions inferred from the alluvial plots at the level of individual genes, we examined heatmaps of persistent and switcher genes across generations for each environmental scenario and genotype (Supplementary Fig. S19-S22). These heatmaps confirmed that WT generally contained fewer genes maintained across the analysed generations, with the strongest persistent set observed under heat, whereas elevated CO_2_ and O_3_ showed little or no persistent multigenerational signal in WT. Switcher genes were also detected in WT, but were comparatively limited and were most evident under heat and the combination scenario, supporting the view that WT largely resets its transcriptome after stress withdrawal while retaining only a restricted core of long-lived responses.

In contrast, *gsnor-ko* displayed broader persistent signatures across multiple environmental scenarios, particularly elevated CO_2_, O_3_, heat, and the combination scenario, as well as larger switcher gene sets than WT. These patterns indicate that, relative to WT, the mutant retains more stress-responsive transcriptional activity after stress removal, especially under conditions that strongly perturb cellular homeostasis. Thus, the heatmap analysis reinforces the alluvial plots by showing that WT more often reverts toward a non-stress transcriptional state, whereas *gsnor-ko* more frequently maintains or reconfigures stress-associated gene expression into later generations (Supplementary Fig. S19-S22).

### 4.2 Recurrently responsive transcription factors identify candidate regulatory nodes

To determine whether stable transcriptome responses were accompanied by recurrent regulation of upstream regulators, we extracted transcription factors (TFs) that were differentially expressed in all three analysed generations within each treatment and visualised their expression patterns across genotypes and environments (Supplementary Fig. S23). This analysis showed that persistent responses were not limited to downstream functional genes but also involved multiple TF families, suggesting that part of the multigenerational signal is retained at the regulatory level.

Across treatments, recurrently responsive TFs belonged to several well-established stress-associated families, including basic helix-loop-helix (bHLH), basic leucine zipper (bZIP), ERF, MYB, WRKY, homeodomain-leucine zipper (HD-ZIP), GAI-RGA-SCR (GRAS), nuclear factor Y (NF-Y), and CONSTANS-like (CO-like) groups. Representative candidates included BRI1-EMS-SUPPRESSOR 1 (BES1), bHLH and ERF family members, MYB-related genes, WRKY transcription factors, NF-YB subunits, and TEOSINTE BRANCHED1/CYCLOIDEA/PROLIFERATING CELL FACTOR (TCP) factors, highlighting regulatory programs potentially associated with stable stress adaptation across generations (references/PubMed IDs in Supplementary File S20).

The strongest and broadest recurrent TF signatures were observed under heat and the combination scenario in WT, and under O_3_, heat, and the combination scenario in *gsnor-ko*, whereas fewer stable TF responses were evident in some WT elevated-CO_2_ and O_3_ contrasts. Although these data do not establish direct regulatory causality, they identify a compact set of candidate TFs that may contribute to persistent or reconfigured stress responses. The broader recurrent TF response in *gsnor-ko* is consistent with reduced transcriptional resetting after stress withdrawal.

### 4.3 Candidate epigenetic regulators are embedded within persistent and switching responses

Because persistent and switcher trajectories are compatible with altered transcriptional memory or incomplete resetting, we next screened stress-responsive genes for candidates with known or proposed roles in DNA methylation, chromatin remodelling, histone modification, histone variants/chaperones, RNA-directed silencing, RNA methylation, and one-carbon metabolism. To organise these candidates, we grouped them into five regulatory models (Model 1-5) and visualised their expression patterns across treatments, generations, and genotypes (Fig. 5-6; Supplementary File S21 includes references and PubMed IDs).

**Fig 5.**
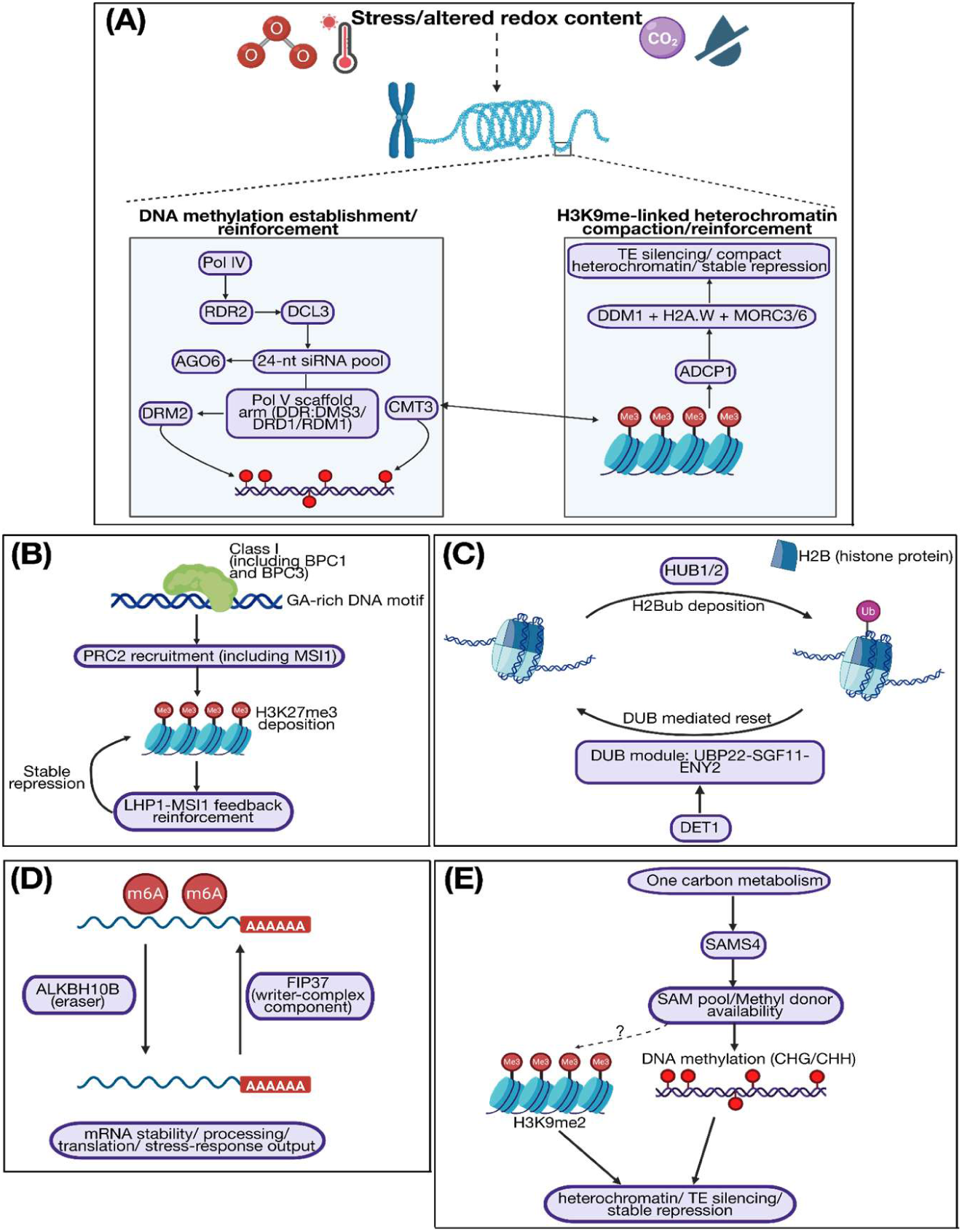
Proposed epigenetic models linking repeated climate stress and altered redox status to multi-generational transcriptional persistence. Schematic representation of five non-mutually exclusive mechanistic models derived from the transcriptomic analyses and the known functions of candidate regulators. **(A)** DNA methylation and heterochromatin reinforcement via RNA-directed DNA methylation (RdDM) and chromatin compaction pathways, including Pol IV, RDR2, DCL3, AGO6, DRM2, CMT3, DMS3/DRD1/RDM1, DDM1, H2A.W, MORC3/6, and ADCP1. **(B)** Polycomb-linked repression and chromatin stabilisation, in which BPC proteins may contribute to PRC2 recruitment (including MSI1), leading to H3K27me3-associated silencing and reinforcement through LHP1-MSI1.**b(C)** Dynamic regulation of histone H2B ubiquitination through HUB1/2-mediated deposition and deubiquitination via the UBP22-SGF11-ENY2 module, potentially linked to DET1-dependent chromatin resetting. **(D)** RNA methylation-based regulation, in which FIP37 and ALKBH10B may influence mRNA stability, processing, translation, and stress-response output. **(E)** One-carbon and methyl-donor metabolism, where SAMS4 and related pathways may affect S-adenosylmethionine availability and thereby influence DNA/histone methylation and stable repression. These models represent working hypotheses and, by themselves, do not demonstrate direct epigenetic inheritance.

**Fig 6.**
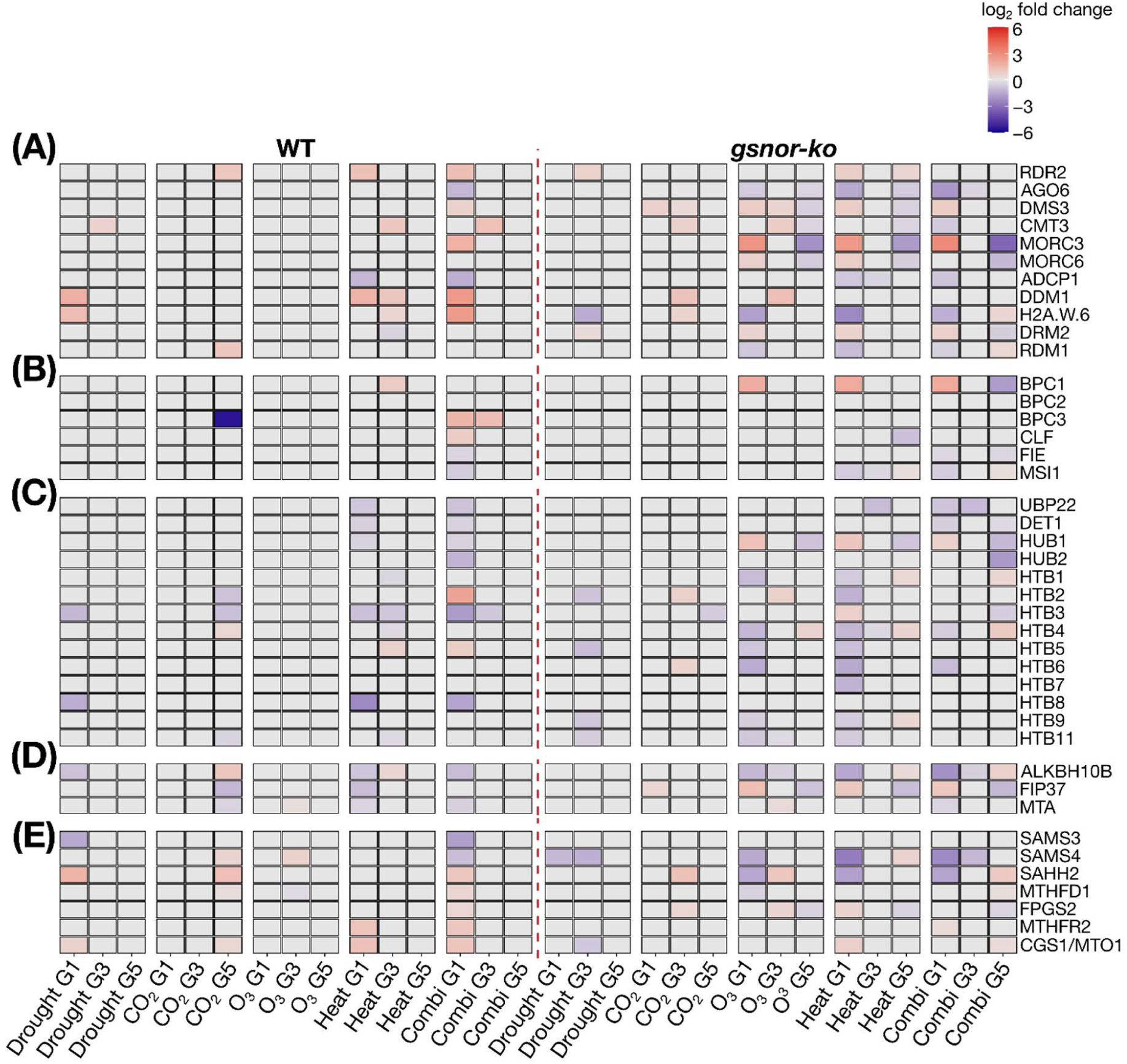
Expression profiles of candidate epigenetic regulators across treatments and generations in WT and *gsnor-ko*. Heatmaps showing log_2_ fold-change values of candidate genes associated with the five proposed epigenetic models across Drought, elevated CO_2_, elevated O_3_, Heat, and Combination treatments in generations G1, G3, and G5 for WT and *gsnor-ko*. **(A)** DNA methylation/RdDM and heterochromatin-associated genes. **(B)** Polycomb- and chromatin repression-associated genes. **(C)** Histone H2B ubiquitination and chromatin resetting-associated genes. **(D)** RNA methylation-associated genes. **(E)** One-carbon and methyl-donor metabolism-associated genes. The dashed vertical line separates WT and *gsnor-ko*. Red indicates higher expression and blue indicates lower expression relative to the corresponding control comparison, with colour intensity proportional to log_2_ fold change.

The strongest and broadest transcriptional changes were observed among genes associated with DNA-methylation establishment/reinforcement and heterochromatin compaction (Model 1). This group included RNA-DEPENDENT RNA POLYMERASE 2 (RDR2), ARGONAUTE 6 (AGO6), DEFECTIVE IN MERISTEM SILENCING 3 (DMS3), CHROMOMETHYLASE 3 (CMT3), MICRORCHIDIA 3/6 (MORC3/6), AGENET DOMAIN CONTAINING PROTEIN 1 (ADCP1), DECREASE IN DNA METHYLATION 1 (DDM1), H2A.W histone variants, DOMAINS REARRANGED METHYLTRANSFERASE 2 (DRM2), and RNA-DIRECTED DNA METHYLATION 1 (RDM1) (Fig. 5A and 6A). These candidates were differentially expressed across multiple stress trajectories, with particularly prominent changes under heat and combination in WT, and under elevated O_3_, heat, and combination in *gsnor-ko*. The occurrence of both positive and negative changes within this group suggests dynamic remodelling rather than a uniform directional response.

Genes associated with Polycomb-mediated repression and H3K27me3-linked regulation (Model 2), including BASIC PENTACYSTEINE 1/2/3 (BPC1/2/3), CURLY LEAF (CLF), FERTILIZATION-INDEPENDENT ENDOSPERM (FIE), and MULTICOPY SUPPRESSOR OF IRA 1 (MSI1), showed a more selective and generally weaker transcriptional response (Fig. 5B and 6B). Nevertheless, recurrent changes in this group, particularly under the combination scenario in WT and under O_3_, heat, and combination in *gsnor-ko*, indicate that Polycomb-related pathways may also contribute to stable or reconfigured transcriptional states across generations.

A third group comprised genes involved in H2B ubiquitination and chromatin resetting (Model 3), including UBIQUITIN-SPECIFIC PROTEASE 22 (UBP22), DE-ETIOLATED 1 (DET1), HISTONE MONOUBIQUITINATION 1 (HUB1), and HISTONE MONOUBIQUITINATION 2 (HUB2), together with several HISTONE H2B genes (Fig. 5C and 6C). These genes again showed broader perturbation in *gsnor-ko* than in WT, especially under O_3_, heat, and combination, suggesting an altered chromatin-resetting capacity in the mutant background.

We also identified stress-responsive genes linked to m^6^A RNA-methylation-related regulation (Model 4), including ALKB HOMOLOG 10B (ALKBH10B), FKBP12-INTERACTING PROTEIN 37 (FIP37), and mRNA ADENOSINE METHYLASE A (MTA), as well as genes associated with one-carbon metabolism and methyl-donor availability (Model 5), including S-ADENOSYLMETHIONINE SYNTHETASE 4 (SAMS4), S-ADENOSYL-L-HOMOCYSTEINE HYDROLASE 2 (SAHH2), METHYLENETETRAHYDROFOLATE DEHYDROGENASE/METHENYLTETRAHYDROFOLATE CYCLOHYDROLASE 1 (MTHFD1), FOLYLPOLYGLUTAMATE SYNTHETASE 2 (FPGS2), METHYLENETETRAHYDROFOLATE REDUCTASE 2 (MTHFR2), and CYSTATHIONINE GAMMA-SYNTHASE 1 / METHIONINE OVERACCUMULATOR 1 (CGS1/MTO1) (Fig. 5D-E and 6D-E). Although these groups were smaller, they recurred across several stress trajectories, again most clearly under heat and combination in WT and under O_3_, heat, and combination in *gsnor-ko*.

Overall, these candidate genes were not confined to a single environmental scenario or regulatory class but recurred across multiple trajectories and mechanistic categories. Importantly, the response was globally broader and more persistent in *gsnor-ko* than in WT, particularly after return to control conditions for two generations, consistent with the alluvial and heatmap analyses indicating reduced transcriptomic resetting in the mutant. However, differential expression alone does not establish epigenetic causality, but the repeated appearance of genes linked to DNA methylation, heterochromatin maintenance, Polycomb repression, histone ubiquitination, RNA methylation, and methyl-donor metabolism supports the idea that chromatin- and RNA-associated regulatory pathways contribute to the maintenance or resetting of environmental responses across generations.

### 4.4 Co-expression network analysis identifies genotype- and generation-associated modules

WGCNA of the combined RNA-seq dataset indicated that module eigengene patterns were influenced more strongly by genotype and generation than by treatment (Supplementary Fig. S24-S29). In particular, the modules darkgreen and lightcyan1 primarily distinguished WT from *gsnor-ko*, whereas magenta was mainly associated with generation and ivory showed the clearest treatment-related signal. Functional enrichment linked darkgreen to plant development and cell cycle, lightcyan1 to photosynthesis and biosynthesis, magenta to cellular respiration and oxidative phosphorylation, and ivory to external stimulus response, defence responses, and protein kinase activities (Supplementary Fig. S26 and S29).

Exploratory module-trait analyses incorporating rosette area and seed weight were heterogeneous and did not reveal robust module-phenotype relationships across the dataset (Supplementary Fig. S28B-S29A-B). Accordingly, WGCNA is presented here as a supportive systems-level analysis that reinforces the DEG-based conclusion that the major transcriptomic structure of the experiment is shaped primarily by genotype background and generational state, with more modest treatment-specific effects.

Because external stimuli response, pathogen and systemic acquired resistance (SAR) terms were identified among the enriched WGCNA modules (ivory), we tested whether these transcriptomic signatures were accompanied by altered pathogen responses (Supplementary Fig. S29-S30). The leaf disease assay showed that disease outcome differed across genotypes, treatment histories, and generations rather than following a single uniform pattern. Across generations, *gsnor-ko* generally displayed higher bacterial growth than WT under control conditions, consistent with increased basal susceptibility (as published by Feechan *et al*., 2005). However, when plants had experienced the combination-stress history, *gsnor-ko* showed reduced bacterial growth relative to its corresponding control history in some generations, indicating a stress-history-associated reduction in susceptibility (Supplementary Fig. S30A-B; Supplementary file S22). Thus, the disease assay supports the conclusion that repeated climate history influenced immune output, consistent with the immunity-associated transcriptomic signatures identified in WGCNA. At the same time, the response was generation-dependent, suggesting that the relationship between prior climate exposure and defence is dynamic rather than constitutively fixed.

## Discussion

*Arabidopsis thaliana* was repeatedly exposed to climate conditions for 3 generations, followed by 2 generations grown under control conditions, and phenotypic, transcriptomic, and network-level analyses were performed. A central outcome of this study is that both genotype background and stress context shaped the extent of recovery after repeated perturbations. In WT, many responses returned to a control-like state after stress withdrawal, whereas *gsnor-ko* more often retained reduced growth, low seed production, and altered transcriptional states. We therefore interpret impaired NO homeostasis as a factor that shifts repeated climate responses away from efficient resetting and toward more persistent stress-associated reprogramming.

### 1. Impaired NO homeostasis reduces recovery across repeated climate stress

Across the phenotypic datasets, repeated climate stress revealed a clear genotype-dependent difference in recovery across generations rather than a single uniform stress effect. In nearly all environments, the size of *gsnor-ko* remained smaller than WT and produced substantially fewer seeds, indicating that impaired NO homeostasis imposes a persistent penalty on both vegetative growth and reproductive output. This is consistent with earlier work showing that GSNOR is required for normal growth/development and thermotolerance, and with later evidence linking NO/GSNOR signalling to chromatin-associated regulation of stress responses (Ageeva-Kieferle *et al*., 2021; Rudolf *et al*., 2021; Lee *et al*., 2008).

The measured traits also differed in their degree of plasticity. Rosette area showed the clearest genotype × generation × treatment structure, whereas QYmax remained comparatively buffered, and NPQ_Lss captured subtler shifts in energy dissipation. Thus, the strongest multigenerational effects were observed at the level of growth and allocation rather than in core PSII efficiency. Seed production sharpened this contrast: WT often showed partial recovery after stress withdrawal, whereas *gsnor-ko* remained uniformly low across treatments and generations. This suggests that the mutant phenotype reflects not only stronger stress sensitivity but also a reduced ability to regain productive performance after repeated perturbations. Similar phenotypic responses for *gsnor-ko* have been reported before in response to biotic stress (Holzmeister *et al*., 2011).

Among the tested scenarios, O_3_, heat, and the combination treatment were especially informative because they are closely linked to redox and nitrosative signalling. O_3_ activates ROS-, NO-, salicylic acid-, and ethylene-linked pathways in Arabidopsis (Ahlfors *et al*., 2009), while heat is the context in which GSNOR function has been most clearly linked to stress tolerance and thermomemory-related processes (Lee *et al*., 2008; Naaz *et al*., 2025). The strong genotype dependence of these scenarios therefore fits a model in which NO/GSNO balance contributes not only to immediate stress responses, but also to the maintenance or clearance of their downstream consequences.

### 2. Multi-generational transcriptome responses distinguish resetting from persistence

The RNA-seq data showed high mapping quality and reproducible multivariate structure, supporting the interpretation that the observed transcriptional patterns are biological rather than technical. More importantly, DAPC and PCA indicated that genotype background and generation contributed more strongly to global transcriptome organisation than any single treatment snapshot. This suggests that a history of repeated stress becomes embedded in the transcriptome as a state variable, rather than appearing only as a transient response in individual contrasts. This systems-level structure is compatible with the broader molecular-memory literature, such as work in poplar under future-climate-relevant drought and heat scenarios, which showed that transcriptomes can differ most strongly during recovery rather than during the stress exposure itself, highlighting post-stress state transitions as a major component of climate resilience (Georgii *et al*., 2019).

In WT, drought, heat, and the combination treatment generally showed declining DEG numbers across generations, and the gene-state analysis indicated fading as a dominant trajectory in several of these conditions. Together, these patterns are consistent with broad transcriptomic resetting after stress withdrawal. By contrast, *gsnor-ko* showed altered DEG magnitude, altered temporal profiles, and broader persistent or switcher behaviour, particularly under O_3_, heat, and combination scenarios, consistent with the view that NO is a central node in plant responses to both elevated greenhouse-gas-related environments and ozone-linked oxidative signalling (Kabange *et al*., 2022). Thus, the mutant differed from WT not only in response strength but also in the duration of stress-responsive states.

Importantly, the combined treatment should not be viewed simply as a stronger version of the single stresses. In Arabidopsis, combined stresses frequently generate transcriptomic and metabolic responses that are not predictable from single-stress experiments alone (Rasmussen *et al*., 2013; Zinta *et al*., 2018). In our dataset, the single treatments helped resolve pathway-specific behaviour—for example, drought- and CO_2_-related shifts versus the more strongly redox-linked O_3_ and heat responses—whereas the combination treatment provided a test of how plants integrate overlapping stress cues and how effectively they reset once stress is removed. The fact that *gsnor-ko* remained more altered than WT under the combination regime, therefore, supports the idea that NO/GSNO balance is particularly important when multiple stress signals converge.

Recent studies of plant stress memory emphasise that recurring stress can leave a measurable molecular imprint without necessarily implying stable meiotic inheritance of a defined epigenetic mark, and that recovery is itself a major component of the memory process (Crisp *et al*., 2016; Harris *et al*., 2023; Hemenway and Gehring, 2023; Oberkofler *et al*., 2021). Work on heat-stress memory and chromatin resetting similarly highlights the role of active recovery mechanisms, including DDM1/MOM1-associated resetting, in determining whether stress states persist or are cleared (Iwasaki and Paszkowski, 2014; Pratx *et al*., 2024; Staacke *et al*., 2025). Our results fit that framework well: WT more often returned toward a control-like transcriptomic configuration, whereas *gsnor-ko* more often remained in a stress-responsive or reconfigured state across generations.

### 3. Chromatin-, RNA-, and methylation-linked signatures nominate candidate mechanisms

The enrichment analyses move the study from descriptive transcriptomics toward mechanism. Both genotypes shared a core set of responses involving photosynthesis, glutaredoxin, S-glutathionylation, nutrient uptake, SnRK1-linked regulation, and stress-associated transcription factors, indicating that repeated climate stress engages overlapping energetic, redox, and regulatory programs. However, the more informative result is that *gsnor-ko* showed broader or stronger recurrence of terms associated with DNA methylation, heterochromatin maintenance, RdDM-linked regulation, histone organisation, ubiquitin-mediated chromatin control, RNA modification, and methyl-donor metabolism.

These enrichments do not demonstrate that a specific chromatin mark or methylation state was inherited across meiosis. They do, however, support the idea that repeated stress engages regulatory layers capable of stabilising or clearing transcriptional states. This interpretation is consistent with work showing that NO can influence chromatin regulation through histone modifications and altered DNA accessibility, and that GSNOR contributes to demethylation-associated regulation of transposable elements and stress-responsive genes (Ageeva-Kieferle *et al*., 2021; Rudolf *et al*., 2021). It is also compatible with current models of abiotic-stress memory in which DNA methylation, histone dynamics, non-coding RNAs, RNA turnover, and RNA-directed silencing act together to shape memory intensity and duration (Cao and Chen, 2024; Crisp *et al*., 2016; Xu *et al*., 2024).

In this context, the candidate regulators identified in our dataset, including factors linked to RdDM and heterochromatin reinforcement, histone ubiquitination, Polycomb-related repression, RNA methylation, and one-carbon metabolism, provide a plausible mechanistic bridge between altered redox signalling and reduced transcriptomic resetting in *gsnor-ko*.

### 4. Conclusions and Outlook

Taken together, the most interesting conclusion is that GSNOR-dependent NO/redox homeostasis acts upstream of the balance between stress-state recovery and stress-state persistence. In WT, this balance more often favours resetting after stress withdrawal, especially after drought, heat, or combined treatments. In *gsnor-ko*, the same balance shifts toward continued engagement or the reconfiguration of stress-responsive programs across generations. Mechanistically, this suggests that NO/GSNO status influences the efficiency with which plants exit a stress-conditioned transcriptional state once the external trigger is removed. Our data therefore support a model of redox-dependent transcriptomic resetting rather than direct evidence of stable meiotic epigenetic inheritance.

This model is relevant in the context of future climate change, where recurrent and overlapping stresses are expected to become more common, and resilience will depend not only on acute tolerance but also on how efficiently plants recover between stress episodes. In that sense, our study complements work on future-climate molecular memory in poplar and current models of plant stress memory by placing NO homeostasis upstream of the resetting problem itself (Fejes *et al*., 2025; Georgii *et al*., 2019;Peer, 2026).

At the same time, the present dataset does not provide direct proof of transgenerational epigenetic inheritance. Rather, it supports multi-generational persistence and incomplete resetting of stress-responsive states, together with repeated engagement of chromatin-, RNA-, and methylation-associated pathways. Direct testing of this model will require profiling DNA methylation, chromatin accessibility, histone modifications, and small RNAs within the same multigenerational framework. Future work should also test whether the candidate regulators identified here, including components linked to RdDM, heterochromatin maintenance, histone ubiquitination, RNA methylation, and methyl-donor metabolism, play causal roles in redox-dependent stress-state recovery. Framed in this way, the study provides a mechanistic foundation for understanding how NO homeostasis shapes plant recovery under recurrent climate stress without overextending beyond the current evidence (Supplementary Fig. S31).

## Supplementary data

Supplementary data are available at Journal of Experimental Botany online and comprise Supplementary Fig. S1-S31 and Supplementary Files S1-S22.

## Supplementary methods and results: RNA-seq computational analysis

### 1. Preprocessing, quantification, alignment, and multivariate statistics

Primary QC and trimming were performed by the sequencing provider (BMK Gene GmbH). Reads were aligned to the TAIR10 *Arabidopsis thaliana* reference genome using HISAT2 (v2.2.1; https://daehwankimlab.github.io/hisat2/main/). RNA-seq libraries showed consistent quality library distribution across conditions and generations, with ∼97% of reads mapped to the TAIR10 reference (Supplementary Fig. S4A-B). Replicate consistency was assessed using hierarchical clustering, and batch terms were included where appropriate (because samples from G1, G3, and G5 were sequenced at different times) (Supplementary Fig. S5). These QC results support robust downstream differential expression analyses.

Reads were assigned to annotated features using featureCounts (Subread v2.0.2) with the following parameters: -T 12 -p -t exon -g Parent –primary. TAIR GFF3 annotation was converted to GTF format using R (Liao *et al*., 2013).

To assess whether global expression profiles contain treatment- and generation-specific signatures, Discriminant Analysis of Principal Components (DAPC; shown as LD1/LD2) was applied to VST-transformed expression data using treatment × generation classes within each genotype (Fig. 3A). In both WT and *gsnor-ko*, samples form compact class-wise clusters in LD space, indicating that stress conditions encode reproducible multigene expression patterns across generations. However, cross-validation performance was moderate (WT CV = 0.62 ± 0.09, *gsnor-ko* CV = 0.69 ± 0.09; Fig. 3A), and the CV confusion matrices show that a subset of samples is mis-assigned to related classes rather than being perfectly separable (Supplementary Fig. S6A–B).

In addition, unsupervised Principal Component Analysis (PCA) shows substantial overlap among treatments within each generation for both genotypes (WT PC1/PC2 = 27.9%/22.6%; gsnor-ko PC1/PC2 = 29.9%/19.7%; Supplementary Fig. S6C-D), consistent with treatment effects being distributed across multiple axes rather than captured by one dominant global component. Therefore, treatment responses were analysed using within-generation differential expression analysis and interpreted condition effects primarily through Differential Expressed Gene (DEG) sets, persistence patterns, and functional enrichment.

### 2. Differential expression analysis

Differential gene expression analysis was performed using DESeq2 in R. Statistical modelling used a design formula based on treatment/condition (treatment vs control), with additional batch terms included as described in section 6.1. Genes were considered differentially expressed at an adjusted p-value < 0.05. No log2 fold-change cutoff was applied because some treatments yielded small DEG sets.

Variance-Stabilising Transformation (VST) was applied for visualisation and clustering. Co-expression network analysis was performed using Weighted Gene Co-expression Network Analysis (WGCNA; soft-threshold network type – unsigned with power of 11, minimum module size – 40, deepSplit – 2, and Pearson correlation was used) (Langfelder *et al*., 2008; Yip and Horvath, 2007). Functional enrichment analyses were performed using Mercator annotation (https://www.plabipd.de/mercator_main.html), applying multi-level categorisation to identify enriched functional categories within gene sets (Supplementary Fig. S1B; Supplementary File 11).

### 3. Circadian marker genes support consistent sampling time across generations

To verify that transcriptome comparisons were not confounded by differences in harvest timing, we examined the expression of canonical morning- and evening-phased clock genes across genotypes, treatments, and generations (Supplementary Fig. S7A-D). Clock-gene profiles showed the expected separation between morning- and evening-associated markers, while overall expression patterns remained broadly comparable across generations within each genotype among Controls, indicating that samples were collected at a similar circadian phase. However, treatment-specific divergence persisted, as shown in PCA and log fold changes in the DEG set (Supplementary Fig. S7A-B and D). Together, these data support that the observed stress-responsive transcriptome changes are unlikely to be driven by differences in sampling time.

## Abbreviations

CO_2_: carbon dioxide
DAPC: discriminant analysis of principal components
DEG: differentially expressed gene
GLM: generalized linear model
*gsnor1-3*: *gsnor-ko*
GSNO: S-nitrosoglutathione
GSNOR: S-nitrosoglutathione reductase
m^6^A: N^6^-methyladenosine
NO: nitric oxide
NPQ_Lss: steady-state non-photochemical quenching
O_3_: ozone
PPFD: photosynthetic photon flux density
PCA: principal component analysis
PSII: photosystem II
QYmax: maximum quantum yield
RdDM: RNA-directed DNA methylation
RNA-seq: RNA sequencing
VST: variance-stabilising transformation
WGCNA: weighted gene co-expression network analysis

## Acknowledgements

This work (for RB and CL) was supported by the Initiative and Networking Fund of the Helmholtz Association under the call HIRS, the EpiCrossBorders: International Helmholtz-Edinburgh Research School for Epigenetics, Biotechnology and UK Research and Innovation (UKRI) Biological Sciences Research Council (BBSRC) grant BB/S010262/1 (to LF) and by the European Research Council (ERC) under the European Union’s Horizon 2020 research and innovation program, grant agreement no. 101001137 (to SHS.).

We thank Peiyuan Zhu, Moritz Popp, Ina Zimmer, Petra Seibel, Franz Buegger, Mustafa Oezden, Alexandros Sigalas, Georg Gerl, Ulrich Junghans, Baris Weber, and Armin Richter for their technical support during experimental work, and Tetyana Nosenko for helping with RNA-seq data pre-processing. We thank Dr. Sophie Haupt and her team at the Plant Growth Facilities of the School of Biological Sciences, University of Edinburgh. We would also like to thank Prof. Dr. Aloys Schepers for his fruitful input during thesis meetings.

During the preparation of this work, the author(s) used AI tools (such as ChatGPT 5.4 and Grammarly 1.162.2.0) in order to polish the writing framework and search relevant research articles (literature overview). After using this tool/service, the author(s) reviewed and edited the content as needed and take(s) full responsibility for the content of the published article.

## Author contributions

Experiments were designed by CL & FJ; experimental work was conducted by RB, AG, BJW, EM, LG, AA, CL & JPS. Analysis was performed by RB; statistical analysis and RNA-seq analysis support were provided by SJ, KFXM, and JPS; disease assays and analysis were performed by LF, SHS & RB; seed phenotyping and analysis was performed by AF, GH & RK; data interpretation was performed by RB, LF & JPS; and the paper was written by RB with input from JPS, AG, BJW, CL, LF, FJ & KFXM. The written paper was reviewed by all the co-authors.

## Data availability

The raw RNA-seq sequencing data supporting this study have been deposited in ArrayExpress under accession **E-MTAB-17127**. Processed data and associated supporting materials, including phenotypic data, the RNA-seq count matrix, supplementary files, and FastQC quality-control reports, have been deposited in Zenodo: https://doi.org/10.5281/zenodo.20343453.

## List of Supplementary Figures

**Supplementary Fig S1. Experimental workflow and RNA-seq analysis pipeline.** (A) Experimental timeline of the multi-generational climate study. Wild-type (Arabidopsis thaliana Col-0) and *gsnor-ko* plants were propagated across successive generations (G1-G3 under future climate conditions and G4-G5 under ambient conditions) under controlled climate conditions. The timeline indicates major experimental stages, including sowing, growth, phenotyping, leaf sampling for RNA extraction, flowering, and seed harvest. Rosette phenotyping was performed during vegetative growth, leaf tissue for RNA-seq was collected at the defined sampling stage, and seeds were harvested at maturity to initiate the next generation. (B) Overview of the RNA-seq analysis workflow. Sequencing reads were aligned to the Arabidopsis reference genome using HISAT2, and gene-level fragment counts were generated with FeatureCounts. Count tables were used for differential gene expression analysis, including variance-stabilising transformation (VST) and extraction of differentially expressed genes (DEGs) with DESeq2. Downstream analyses included weighted gene co-expression network analysis (WGCNA) and functional enrichment analysis using Mercator-based annotation.

**Supplementary Fig S2. Climate simulation profiles across the multi-generational experiment.** Environmental conditions applied in control and treated chambers during the five-generation experiment. Control conditions corresponded to G0, G4, and G5, whereas treated conditions were applied during G1- G3. (A) Simulated temperature profiles in control and treated chambers across generations. (B) Relative humidity profiles in control and treated chambers across generations. (C) Photosynthetic Photon Flux Density (PPFD) profile across the experimental timeline. (D) CO2 fumigation profile during the treated generations (G1-G3). (E) O3 fumigation profile during the treated generations (G1-G3). Blue lines indicate control conditions and orange lines indicate treated conditions. Mean differences between treated and control conditions are shown within the relevant panels. These plots document the environmental separation achieved between control and future-climate treatments across the multi-generational design.

**Supplementary Fig S3. Calibration of pot weight to soil water potential and drought profiles across generations.** (A) Calibration curve relating pot weight to soil water potential used to estimate drought progression during the experiment. The fitted linear regression and coefficient of determination (R2) are shown in the panel. (B–E) Estimated soil water potential profiles derived from pot-weight measurements for WT and *gsnor-ko* during the stressed generations (G1+G2+G3) and recovery generations (G4+G5). Separate panels are shown for WT and *gsnor-ko*, illustrating drought-treated and control conditions across the multi-generational experiment. Shaded areas indicate variability among measurements where shown. These plots document the temporal progression of water limitation and recovery in the drought treatment.

**Supplementary Fig S4. Quality assessment of RNA-seq libraries.** (A) Summary of RNA-seq library quality across all WT and gsnor-ko samples from generations G1, G3, and G5 under control and stress treatments. Bars show the percentage of bases with Phred quality scores above the indicated threshold, providing an overview of overall sequencing quality across libraries. (B) Representative per-base sequence quality profile for one RNA-seq library, shown separately for WT and *gsnor-ko*. The blue horizontal line indicates the quality threshold, and the coloured background denotes commonly used quality ranges. These plots show that the sequenced libraries were of consistently high quality and suitable for downstream alignment and differential expression analyses.

**Supplementary Fig S5. Hierarchical clustering of RNA-seq samples.** Circular dendrograms showing hierarchical clustering of RNA-seq samples from WT and gsnor-ko across generations G1, G3, and G5 under Control, Drought, elevated CO_2_, elevated O_3_, Heat, and Combination treatments. Sample labels are colour-coded by treatment, and generation is indicated in the sample names. The upper panel shows clustering for WT (A), and the lower panel (B) shows clustering for *gsnor-ko*. Red dashed lines indicate the major cluster divisions. These dendrograms illustrate the overall relatedness among samples and the extent to which transcriptomic profiles group by treatment and/or generation.

**Supplementary Fig S6. Classification performance and unsupervised transcriptome structure in WT and gsnor-ko.** (A, B) Cross-validation confusion matrices for discriminant analysis of principal components (DAPC) performed on WT (A) and *gsnor-ko* (B) RNA-seq datasets. Values are shown as row-wise percentages, with darker shading indicating lower classification frequency and lighter shading indicating higher classification frequency. Diagonal enrichment indicates correct classification of treatment × generation classes. (C, D) Principal component analysis (PCA) of VST-transformed RNA-seq data for WT (C) and *gsnor-ko* (D), shown separately for generations G1, G3, and G5. Colours indicate treatments (Control, Drought, elevated CO_2_, elevated O_3_, Heat, and Combination). Symbols distinguish generations. These plots show the unsupervised distribution of samples in transcriptome space and complement the DAPC-based class separation shown in Fig. 3A.

**Supplementary Fig S7. Clock gene analysis across genotypes, treatments, and generations.** (A) Principal component analysis (PCA) based on VST-transformed expression values of selected circadian clock genes in WT and gsnor-ko across generations G1, G3, and G5. Colours indicate treatments and symbols indicate treatment classes as shown in the legend. (B) Distribution of the expression metric used to compare morning- and evening-phased clock genes across genotype × generation × treatment combinations. Boxplots show the median, interquartile range, and individual data points for WT and gsnor-ko. (C) Heatmap of normalised expression values for representative morning- and evening-expressed clock genes across WT and *gsnor-ko* samples from control conditions in G1, G3, and G5. Morning genes are labelled in blue and evening genes in red. (D) Heatmap of log2 fold-change values for selected clock genes across treatments and generations in WT and *gsnor-ko*. Red indicates higher expression and blue indicates lower expression relative to the corresponding control comparison. Together, these analyses show that core clock gene expression patterns remained broadly consistent across generations and support comparable sampling time points for transcriptome analysis.

**Supplementary Fig. S8. Normalized growth and photoprotective responses across generations in WT and gsnor-ko.** (A) Rosette area expressed as a percentage of the corresponding same-generation control within each genotype for WT and gsnor-ko across G1–G5. (B) NPQ_Lss expressed as a percentage of the corresponding same-generation control within each genotype for WT *and gsnor-ko* across G1, G2, G3, and G5. Values are shown for Drought, CO_2_, O*_3_*, Heat, and Combination treatments; colours indicate treatment identity. The dashed horizontal line marks 100%, representing the matched control level for each genotype and generation. Points show mean normalized values, and error bars indicate SE. No fluorescence measurements were available for G4 because of technical issues during data acquisition.

**Supplementary Fig S9. Phenotypic data at week 3.** (A) Representative RGB images of WT and *gsnor-ko* plants grown under Control, Drought, elevated CO_2_, elevated O_3_, Heat, and Combination treatments across generations G1-G5 at week 3. (B) Rosette area at week 3 for WT and gsnor-ko across treatments and generations. Boxplots show the distribution of individual values, with points representing biological replicates. Beige shading indicates generations exposed to stress treatment; G4 and G5 were grown under control conditions after stress withdrawal. Different letters indicate significant differences among genotype × generation groups within each treatment (GLM followed by post hoc multiple comparison, P < 0.05).

**Supplementary Fig S10. QYmax at week 4 across treatments and generations.** (A) Representative colour images of QYmax in WT and gsnor-ko plants grown under Control, Drought, elevated CO_2_, elevated O_3_, Heat, and Combination treatments in generations G1, G2, G3, and G5. (B) Boxplots showing QYmax values for WT and *gsnor-ko* across treatments and generations at week 4. Points represent biological replicates. Beige shading indicates generations exposed to stress treatment. Different letters indicate significant differences among genotype × generation groups within each treatment (GLM followed by a post hoc multiple-comparison test). P < 0.05). QYmax measurements were unavailable for G4 due to technical issues during data acquisition.

**Supplementary Fig S11. QYmax at week 3 across treatments and generations.** (A) Representative colour images of QYmax in WT and gsnor-ko plants grown under Control, Drought, elevated CO_2_, elevated O_3_, Heat, and Combination treatments in generations G1, G2, G3, and G5. (B) Boxplots showing QYmax values for WT and *gsnor-ko* across treatments and generations at week 3. Points represent biological replicates. Beige shading indicates generations exposed to stress treatment. Different letters indicate significant differences among genotype × generation groups within each treatment (GLM followed by post hoc multiple comparison, P < 0.05). QYmax measurements were unavailable for G4 due to technical issues during data acquisition.

**Supplementary Fig S12. NPQ Lss at week 4 across treatments and generations.** (A) Representative colour images of NPQ Lss in WT and gsnor-ko plants grown under Control, Drought, elevated CO_2_, elevated O_3_, Heat, and Combination treatments in generations G1, G2, G3, and G5. (B) Boxplots showing NPQ Lss values for WT and *gsnor-ko* across treatments and generations at week 4. Points represent biological replicates. Beige shading indicates generations exposed to stress treatment. Different letters indicate significant differences among genotype × generation groups within each treatment (GLM followed by post hoc multiple comparison, P < 0.05). NPQ Lss measurements were unavailable for G4 due to technical issues during data acquisition.

**Supplementary Fig S13. NPQ Lss at week 3 across treatments and generations.** (A) Representative colour images of NPQ Lss in WT and gsnor-ko plants grown under Control, Drought, elevated CO_2_, elevated O_3_, Heat, and Combination treatments in generations G1, G2, G3, and G5. (B) Boxplots showing NPQ Lss values for WT and gsnor-ko across treatments and generations at week 3. Points represent biological replicates. Beige shading indicates generations exposed to stress treatment. Different letters indicate significant differences among genotype × generation groups within each treatment (GLM followed by post hoc multiple comparison, P < 0.05). NPQ Lss measurements were unavailable for G4 due to technical issues during data acquisition.

**Supplementary Fig S14. Seed weight and seed phenotyping across treatments and generations.** (A) Seed weight per plant in WT and gsnor-ko across Control, Drought, elevated CO_2_, elevated O_3_, Heat, and Combination treatments in generations G1-G5. Boxplots show individual biological replicates; WT is shown in grey and gsnor-ko in blue. Beige shading indicates generations exposed to stress treatment. Different letters indicate significant differences among genotype × generation groups within each treatment (GLM followed by post hoc multiple comparison, P < 0.05). (B) Seed mass distribution in WT across treatments and generations G1, G3, and G5. Boxplots show seed mass for individual seeds, with colours indicating generations and points indicating technical repetitions. (C) Density distributions of seed mass in WT across treatments and generations G1, G3, and G5. Colours indicate generations. Seed phenotyping in panels (B, C) was performed only for WT because insufficient seed material was available for comparable analysis in *gsnor-ko*.

**Supplementary Fig S15. Volcano plots of differential gene expression across treatments and generations in WT and *gsnor-ko***. Volcano plots showing differentially expressed genes in (A) WT and (B) gsnor-ko for Drought, elevated CO_2_, elevated O3, Heat, and Combination treatments in generations G1, G3, and G5. Red points indicate up-regulated genes and blue points indicate down-regulated genes. Dashed orange lines indicate the applied significance (P<0.05) and fold-change thresholds used to define differentially expressed genes. Numbers in the upper corners of each panel indicate the total numbers of down-regulated (blue) and up-regulated (red) genes for the corresponding contrast.

**Supplementary Fig S16. Overlap of differentially expressed genes across treatments, genotypes, and generations.** (A) UpSet plots showing overlap of differentially expressed gene sets between WT and gsnor-ko across generations G1, G3, and G5 for each treatment. Separate plots are shown for the elevated CO_2_, elevated O_3_, Heat, and Combination treatments. Bar plots indicate intersection sizes and set sizes for the corresponding DEG sets. (B) Multi-set Venn diagrams showing overlap of differentially expressed genes among Drought, elevated CO2, elevated O3, Heat, and Combination treatments within each genotype and generation. Separate diagrams are shown for WT G1, *gsnor-ko* G1, WT G3, *gsnor-ko* G3, WT G5, and *gsnor-ko* G5. Numbers indicate unique and shared DEGs among treatments.

**Supplementary Fig S17. Mercator-based functional enrichment analysis across treatments, generations, and genotypes.** (A-T) Bubble plots showing significantly enriched Mercator categories for differentially expressed genes (DEGs) identified in WT and *gsnor-ko* plants under Drought, elevated CO_2_, elevated O*3*, Heat, and Combination treatments across generations G1, G3, and G5. Separate panels represent individual genotype × treatment × DEG-direction subsets. The x-axis indicates generation, and the y-axis shows enriched Mercator terms. Bubble size corresponds to the number of genes assigned to each term, and bubble colour indicates enrichment significance as -log10 adjusted P-value. The coloured bars at the right indicate Mercator hierarchical levels.

**Supplementary Fig S18. Alluvial plots showing multigenerational transitions in DEG states across treatments in WT and *gsnor-ko*.** Alluvial plots showing the distribution and transitions of differential expression states across generations G1, G3, and G5 for WT and *gsnor-ko* plants under Drought, elevated CO_2_, O_3_, Heat, and Combination treatments. For each treatment, genes were classified at each generation as Up, Down, or None (not significantly differentially expressed relative to the corresponding control). The height of each stratum represents the number of genes in a given state, and the connecting flows indicate how genes changed or maintained their expression state across generations. Panels highlight treatment-specific differences in transcriptomic persistence, resetting, and state switching between WT and *gsnor-ko*.

**Supplementary Fig S19. Persistent DEG signatures across generations in WT.** Heatmaps showing persistent differentially expressed genes (DEGs) in WT across generations under the indicated climate treatments. Persistent genes were defined as genes that remained differentially expressed in the same direction across either three generations (G1, G3, and G5) or two generations. Separate panels summarise treatment-specific persistent gene sets, and the table indicates which treatment categories yielded genes in each persistence class. The large heatmap summarises the combined persistent gene set across treatments. Columns represent generations and rows represent genes. Colour scale indicates relative expression change, with red indicating higher and blue indicating lower expression.

**Supplementary Fig S20. Persistent DEG signatures across generations in *gsnor-ko*.** Heatmaps showing persistent differentially expressed genes (DEGs) in *gsnor-ko* across generations under the indicated climate treatments. Persistent genes were defined as genes that remained differentially expressed in the same direction across either three generations (G1, G3, and G5) or two generations. Separate panels show treatment-specific persistent gene sets, and the summary table indicates which treatments yielded persistent genes. The large heatmap summarises the full set of persistent genes detected in *gsnor-ko*. Columns represent generations and rows represent genes. Colour scale indicates relative expression change, with red indicating higher and blue indicating lower expression.

**Supplementary Fig S21. Switcher genes across generations in WT.** Heatmaps showing switcher genes in WT, defined as genes that were differentially expressed across generations and changed the direction of regulation (e.g., from up-regulated to down-regulated, or vice versa). Separate panels show treatment-specific switcher-gene sets, and the accompanying table summarises whether switcher genes were identified across three generations or two generations for each treatment. The large heatmap summarises the combined switcher-gene set in WT. Columns represent generations and rows represent genes. Colour scale indicates relative expression change, with red indicating higher and blue indicating lower expression.

**Supplementary Fig S22. Switcher genes across generations in *gsnor-ko*.** Heatmaps showing switcher genes in *gsnor-ko*, defined as genes that were differentially expressed across generations and reversed the direction of regulation. Separate panels show treatment-specific switcher-gene sets, and the accompanying table summarises whether switcher genes were identified across three generations or two generations for each treatment. The large heatmap summarises the combined switcher-gene set in *gsnor-ko*. Columns represent generations and rows represent genes. Colour scale indicates relative expression change, with red indicating higher and blue indicating lower expression.

**Supplementary Fig S23. Common transcription factors shared across generations and treatments in WT and *gsnor-ko*.** Heatmap showing transcription factors (TFs) identified as common across multiple generations and treatment conditions in WT and *gsnor-ko* plants. Columns represent treatment × generation combinations, arranged by genotype, with WT shown on the left and *gsnor-ko* on the right; the red dashed line indicates the separation between the two genotypes. Rows represent shared TF genes. Colour scale indicates relative expression change, with red representing higher expression and blue representing lower expression. This overview highlights recurrent TF-associated regulatory signatures that were maintained across the multigenerational climate-stress design and reveals both shared and genotype-dependent patterns of transcriptional regulation.

**Supplementary Fig S24. WGCNA network construction and module eigengene relationships.** (A) Dendrogram of all RNA-seq libraries included in the WGCNA, shown together with a trait heatmap summarising week 3 rosette area, week 4 rosette area, and seed weight across samples. (B) Scale-free topology fit index across candidate soft-thresholding powers. (C) Mean connectivity across the same range of powers. (D) Hierarchical clustering of module eigengenes; the red line indicates the threshold used for module merging.

**Supplementary Fig S25. Gene clustering and representative WGCNA modules selected for downstream analysis.** (A) Hierarchical clustering of genes used for WGCNA, with module assignment shown before and after module merging (DynamicTree cut and Merged Tree cut). (B) Summary table of representative modules selected for downstream analysis, including their putative role, hub gene, and annotated function. Highlighted modules include darkgreen and lightcyan1 as genotype-associated modules, magenta as a generation-trend module, and ivory as a treatment-responsive module.

**Supplementary Fig S26. Module eigengene correlations across combined treatment groups.** Heatmap showing Pearson correlations between WGCNA module eigengenes and treatment × generation combinations in WT and *gsnor-ko*. Columns represent Control, Drought, CO₂, O₃, Heat, and Combination at G1, G3, and G5 for each genotype; rows represent individual co-expression modules. Values in cells indicate correlation coefficients, with P-values shown in parentheses. Dashed boxes highlight representative modules predominantly associated with genotype, generation, or treatment differences.

**Supplementary Fig S27. Genotype- and treatment-stratified module eigengene correlations.** (A) Heatmap of Pearson correlations between WGCNA module eigengenes and genotype × generation classes (WT G1, WT G3, WT G5*, gsnor-ko* G1, gsnor-ko G3, *gsnor-ko* G5). (B) Heatmap of Pearson correlations between module eigengenes and genotype × treatment classes in WT and *gsnor-ko*. Cell values show correlation coefficients, with P-values given in parentheses. These plots highlight modules whose behaviour is more strongly associated with genotype background and generation than with treatment alone.

**Supplementary Fig S28. Within-group correlations between WGCNA modules and phenotypic traits.** Heatmaps showing Pearson correlations between WGCNA module eigengenes and phenotypic measurements calculated within experimental groups. Rows represent co-expression modules and columns represent treatment × generation combinations in WT and *gsnor-ko*. Cell values indicate correlation coefficients, with P-values shown in parentheses. Upper and lower panels summarise separate within-group module–trait correlation analyses across the phenotypic dataset: (A) leaf area week 3; (B) leaf area week 4.

**Supplementary Fig S29. Module–trait relationships and Mercator enrichment of selected WGCNA modules.** (A) Heatmaps showing Pearson correlations between WGCNA module eigengenes and phenotypic measurements calculated within experimental groups. Rows represent co-expression modules and columns represent treatment × generation combinations in WT and *gsnor-ko*. Cell values indicate correlation coefficients, with P-values shown in parentheses. Upper and lower panels summarise separate within-group module–trait correlation analyses across the phenotypic dataset: weight of seeds (B), Mercator-based enrichment analysis of the selected modules darkgreen, lightcyan1, magenta, and ivory. Bubble size indicates the number of genes assigned to each term, and colour indicates adjusted P-value. Side annotation bars indicate module identity and Mercator functional level.

**Supplementary Fig S30. Disease assay in WT and gsnor-ko under control and combination treatments across generations.** (A) Representative detached leaves from WT and gsnor-ko plants grown under control conditions (combination having treatment history) and assayed at G1, G3, and G5. Leaves are shown after pathogen challenge, with visible differences in lesion development and tissue collapse between genotypes and treatment histories. Scale bars indicate 12 cm. (B) Quantification of bacterial growth in infected leaves, expressed as log10(CFU) per leaf disk, for WT and *gsnor-ko* under control and combination conditions at G1, G3, and G5. Boxplots show the distribution of values across biological replicates; points represent individual samples. Different letters indicate significant differences among groups within each generation.

**Supplementary Fig S31. Outlook model summarising the proposed divergence between WT and *gsnor-ko* under repeated climate stress.**

## Supplementary Files Information

**Supplementary File S1:** Original climate records downloaded from the German Weather Service (DWD), together with the chamber-date conversion table used for climate simulation. The file includes hourly temperature and relative humidity data for the years 1993 and 2003.

**Supplementary File S2:** Simulated hourly temperature data across generations for control and treated climate scenarios, including mean temperature, standard deviation, sample size, day/hour indices, and observation status.

**Supplementary File S3**: Simulated hourly relative humidity data across generations for control and treated climate scenarios, including mean humidity, standard deviation, sample size, day/hour indices, and observation status.

**Supplementary File S4:** Hourly photosynthetically active radiation (PAR) data across generations, including mean PAR, standard deviation, source/sensor information, and sample size.

**Supplementary File S5**: Hourly CO2 and O3 fumigation data across generations, including gas type, treatment group, mean concentration, standard deviation, sample size, and day/hour indices.

**Supplementary File S6**: Weight of pots and predicted soil water potential values for WT plants across generations and treatments, including day of measurement, model output, and out-of-range flags.

**Supplementary File S7**: Weight of pots and predicted soil water potential values for *gsnor-ko* plants across generations and treatments, including day of measurement, model output, and out-of-range flags.

**Supplementary File S8:** Raw phenotyping data from week 3, including generation, genotype, treatment, rosette area, roundness, Fv/Fm-Lss, QYmax, and NPQ-Lss.

**Supplementary File S9:** Raw phenotyping data from week 4, including generation, genotype, treatment, rosette area, roundness, Fv/Fm-Lss, QYmax, and NPQ-Lss.

**Supplementary File S10**: Raw seed weight data, including generation, genotype, treatment, and individual seed weight measurements.

**Supplementary File S11**: Mercator4 functional annotation output for Arabidopsis thaliana genes used in the enrichment analysis.

**Supplementary File S12:** Overall statistical results for phenotypic traits measured at week 3 and week 4, including effects of genotype, treatment, generation, and their interactions.

**Supplementary File S13:** Estimated mean values for phenotypic traits at week 3 and week 4 across genotypes, treatments, and generations, including standard errors and confidence intervals.

**Supplementary File S14**: Pairwise statistical comparison of WT and *gsnor-ko* for phenotypic traits at week 3 and week 4 across treatments and generations.

**Supplementary File S15:** Pairwise treatment comparisons for phenotypic traits at week 3 and week 4 within each genotype and generation.

**Supplementary File S16:** Estimated mean seed weight values across genotypes, treatments, and generations, including standard errors and confidence intervals.

**Supplementary File S17**: Statistical comparisons of seed weight across generations within each treatment and genotype.

**Supplementary File S18:** Individual seed phenotyping measurements across generations and treatments.

**Supplementary File S19:** Estimated mean seed mass values across genotypes, treatments, and generations.

**Supplementary File S20:** Transcription factors discussed in this study, including gene IDs, transcription factor families, short functional notes, and supporting literature references.

**Supplementary File S21:** Candidate epigenetic regulators and associated genes identified in this study, including expression patterns, occurrence in the dataset, proposed epigenetic links, and supporting literature references.

**Supplementary File S22:** Quantitative disease assay data for WT and *gsnor-ko* plants across generations G1, G3, and G5. Separate sheets are provided for each generation. Within each sheet, values are listed by genotype and treatment history (Control and Combination) and contain the quantitative measurements used for disease-response comparisons.

