## Supplementary Figures for "GSNOR-dependent nitric oxide homeostasis promotes recovery from repeated climate stress across generations in *Arabidopsis thaliana*"

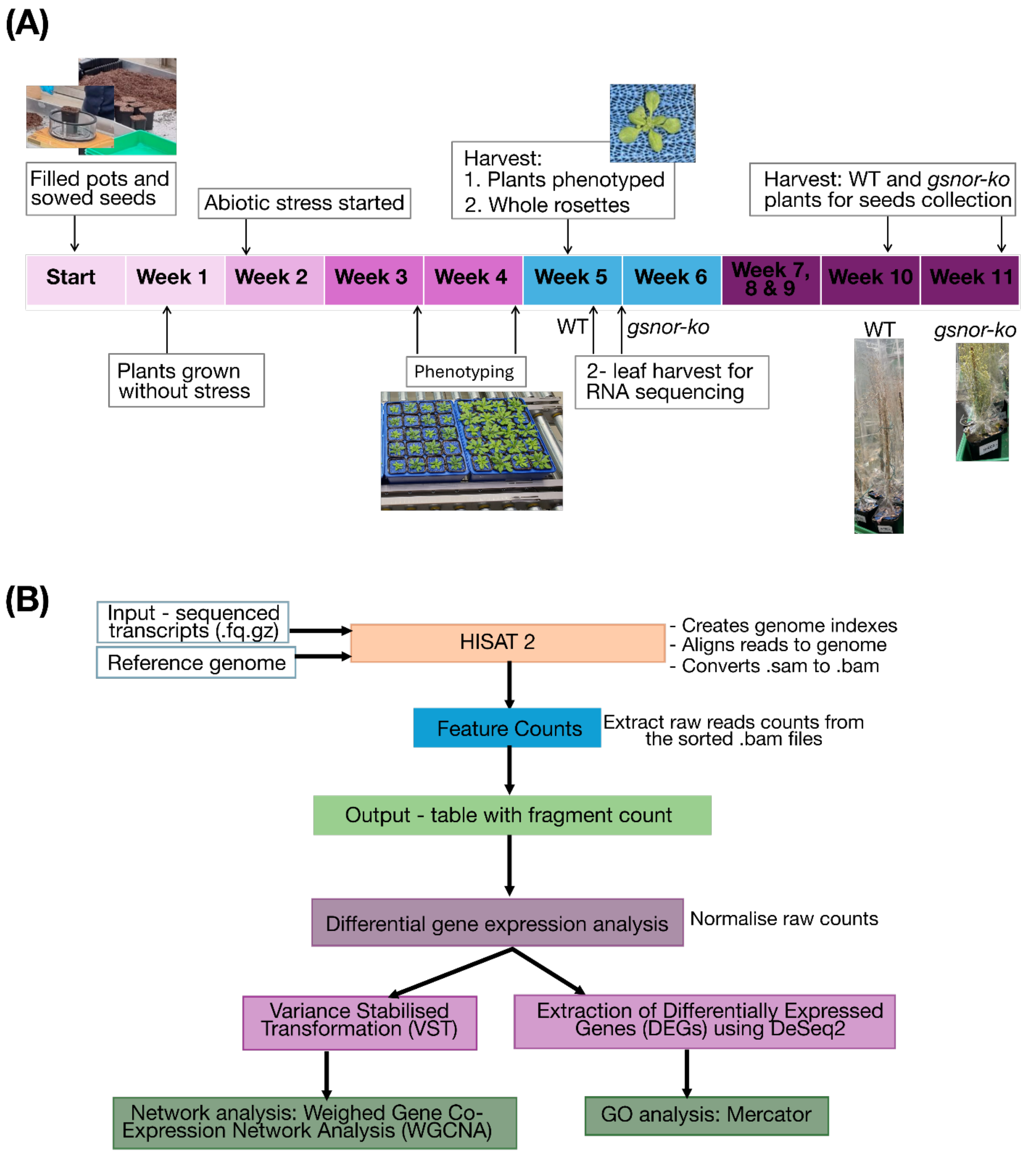


**Supplementary Fig S1. Experimental workflow and RNA-seq analysis pipeline.** **(A)** Experimental timeline of the multi-generational climate study. Wild-type (*Arabidopsis thaliana* Col-0) and *gsnor-ko* plants were propagated across successive generations (G1-G3 under future climate conditions and G4-G5 under ambient conditions) under controlled climate conditions. The timeline indicates major experimental stages, including sowing, growth, phenotyping, leaf sampling for RNA extraction, flowering, and seed harvest. Rosette phenotyping was performed during vegetative growth, leaf tissue for RNA-seq was collected at the defined sampling stage, and seeds were harvested at maturity to initiate the next generation.

**(B)** Overview of the RNA-seq analysis workflow. Sequencing reads were aligned to the Arabidopsis reference genome using HISAT2, and gene-level fragment counts were generated with FeatureCounts. Count tables were used for differential gene expression analysis, including variance-stabilising transformation (VST) and extraction of differentially expressed genes

(DEGs) with DESeq2. Downstream analyses included weighted gene co-expression network analysis (WGCNA) and functional enrichment analysis using Mercator-based annotation.

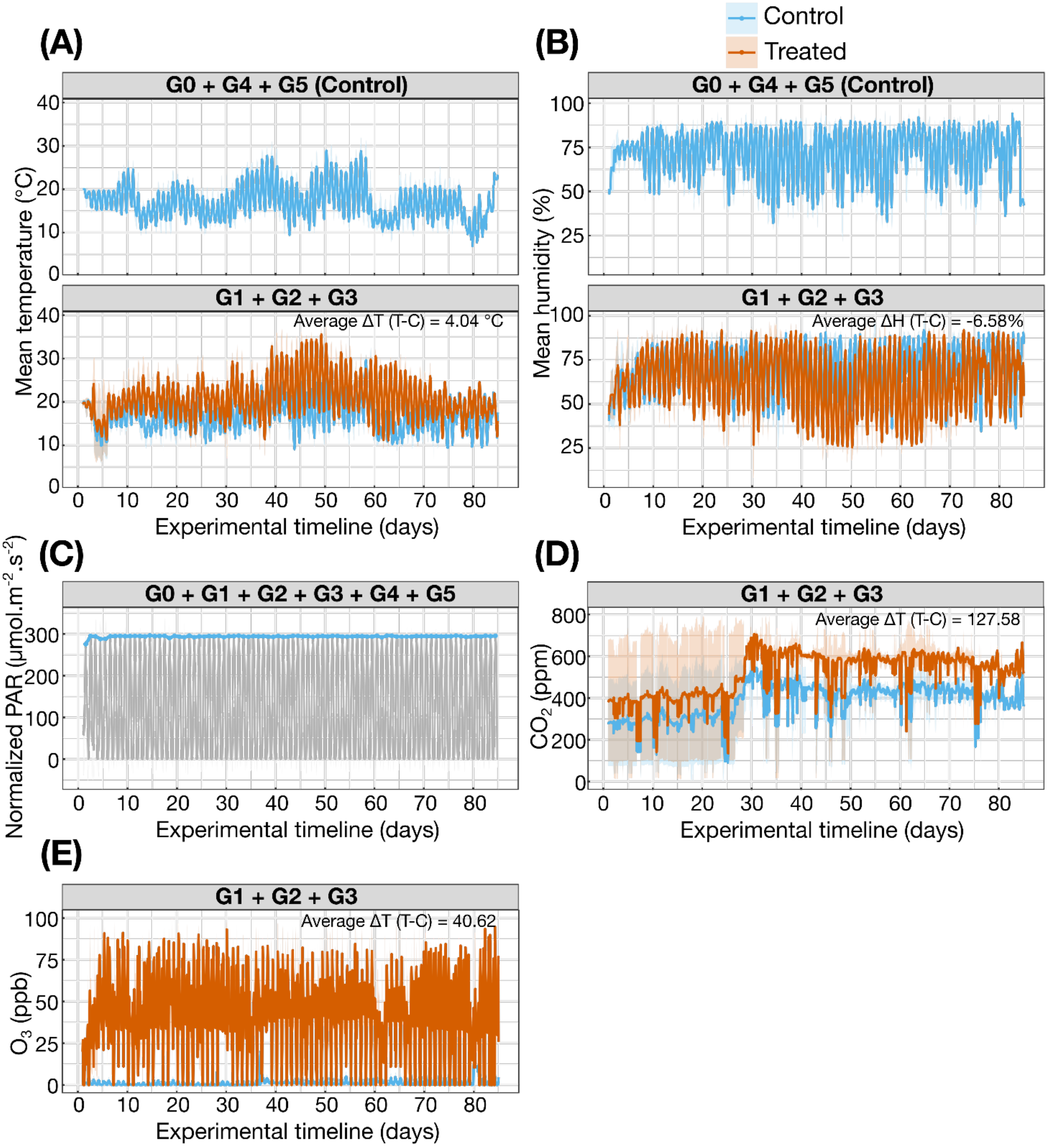


**Supplementary Fig S2. Climate simulation profiles across the multi-generational experiment.**

Environmental conditions applied in control and treated chambers during the five-generation experiment. Control conditions corresponded to G0, G4, and G5, whereas treated conditions were applied during G1-G3. **(A)** Simulated temperature profiles in control and treated chambers across generations. **(B)** Relative humidity profiles in control and treated chambers across generations. **(C)** Photosynthetic Photon Flux Density (PPFD) profile across the experimental timeline. **(D)** $CO_2$ fumigation profile during the treated generations (G1-G3). **(E)** $O_3$ fumigation profile during the treated generations (G1-G3). Blue lines indicate control conditions and orange lines indicate treated conditions. Mean differences between treated and control conditions are shown within the relevant panels. These plots document the environmental separation achieved between control and future-climate treatments across the multi-generational design.

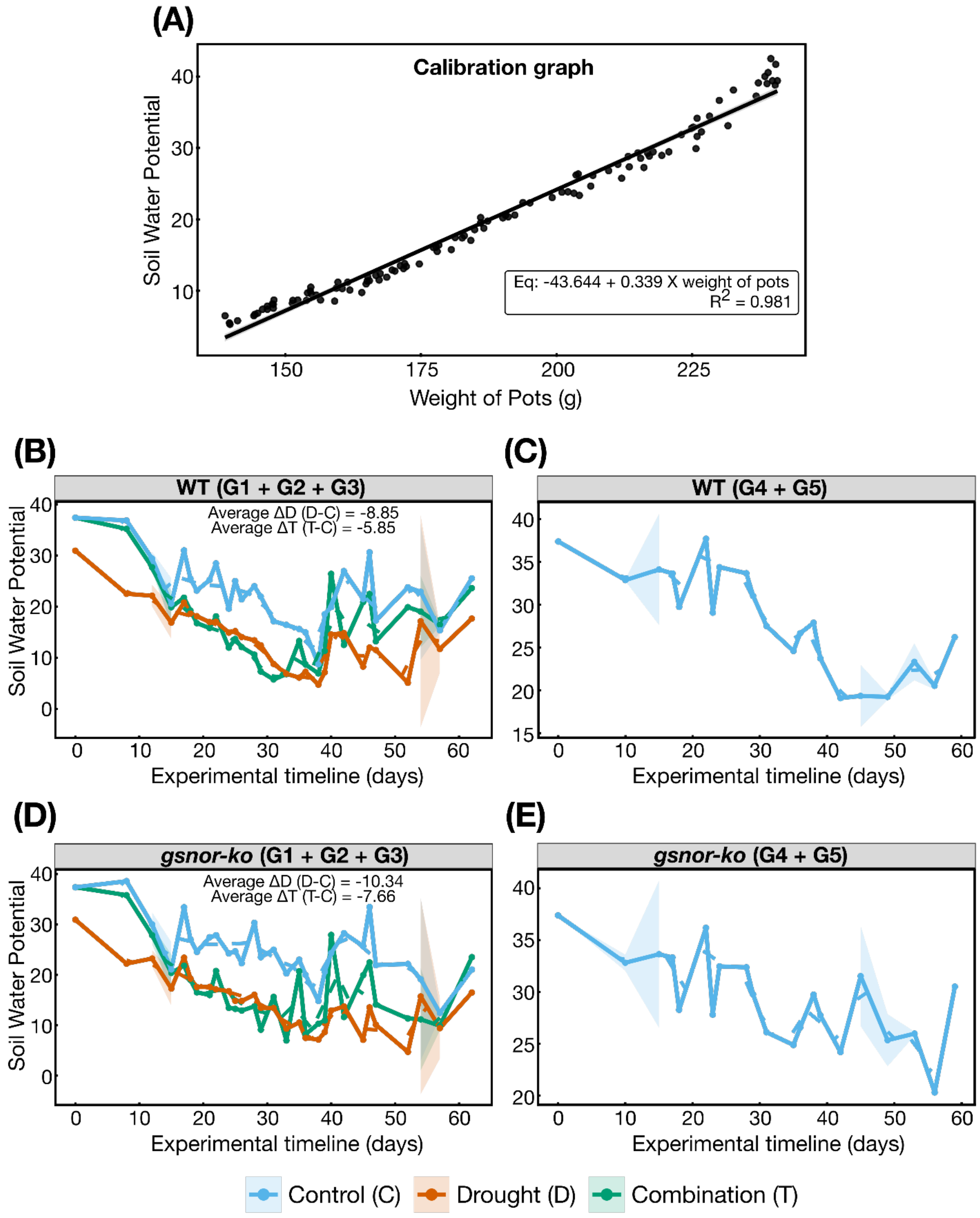


**Supplementary Fig S3. Calibration of pot weight to soil water potential and drought profiles across generations. (A)** Calibration curve relating pot weight to soil water potential used to estimate drought progression during the experiment. The fitted linear regression and coefficient of determination ($R^2$) are shown in the panel. **(B–E)** Estimated soil water potential profiles derived from pot-weight measurements for WT and *gsnor-ko* during the stressed generations (G1+G2+G3) and recovery generations (G4+G5). Separate panels are shown for WT and *gsnor-ko*, illustrating drought-treated and control conditions across the multi-generational experiment. Shaded areas indicate variability among measurements where shown. These plots document the temporal progression of water limitation and recovery in the drought treatment.

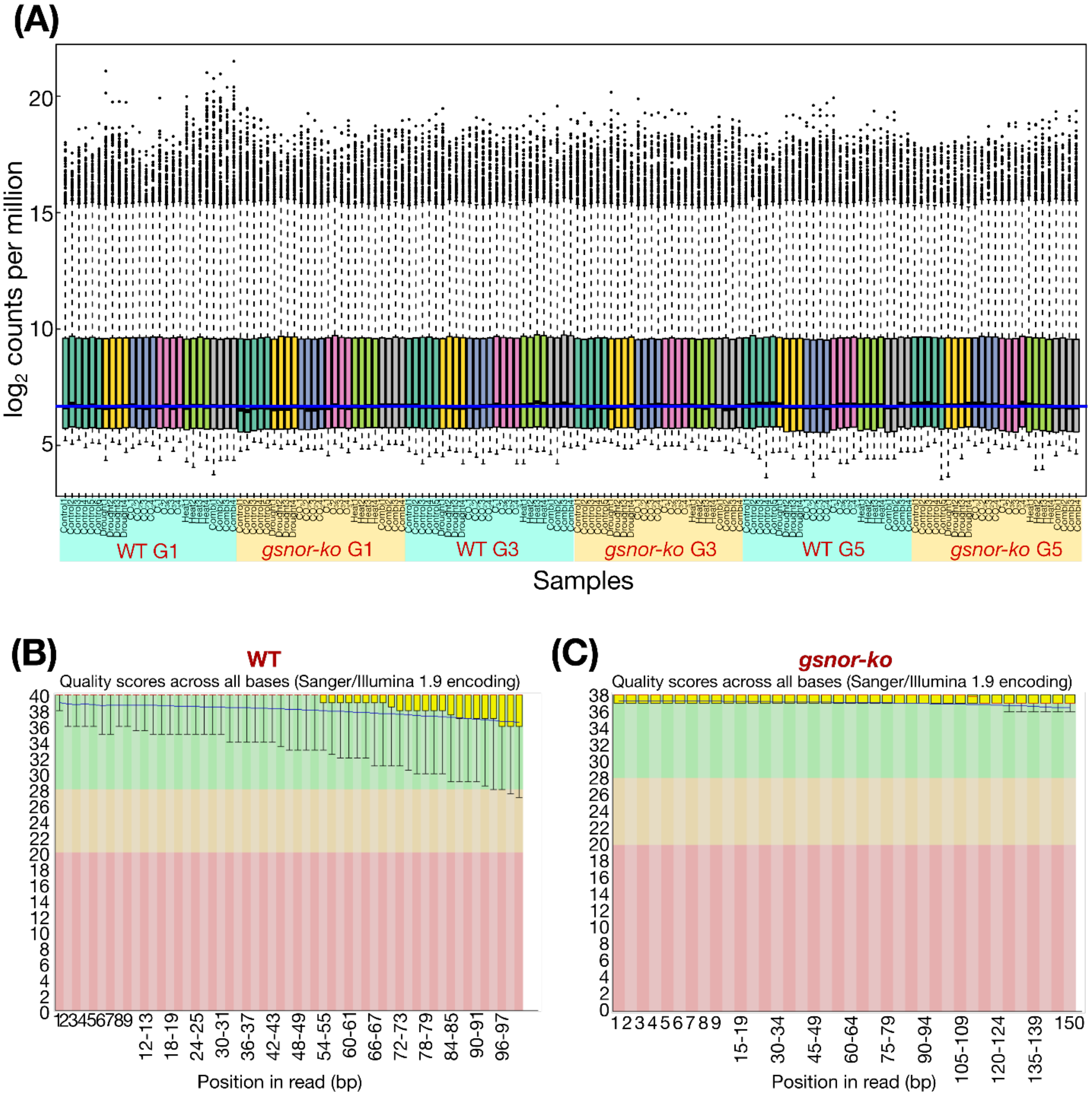


**Supplementary Fig S4. Quality assessment of RNA-seq libraries. (A)** Summary of RNA-seq library quality across all WT and *gsnor-ko* samples from generations G1, G3, and G5 under control and stress treatments. Bars show the percentage of bases with Phred quality scores above the indicated threshold, providing an overview of overall sequencing quality across libraries. **(B)** Representative per-base sequence quality profile for one RNA-seq library, shown separately for WT and *gsnor-ko*. The blue horizontal line indicates the quality threshold, and the coloured background denotes commonly used quality ranges. These plots show that the sequenced libraries were of consistently high quality and suitable for downstream alignment and differential expression analyses.

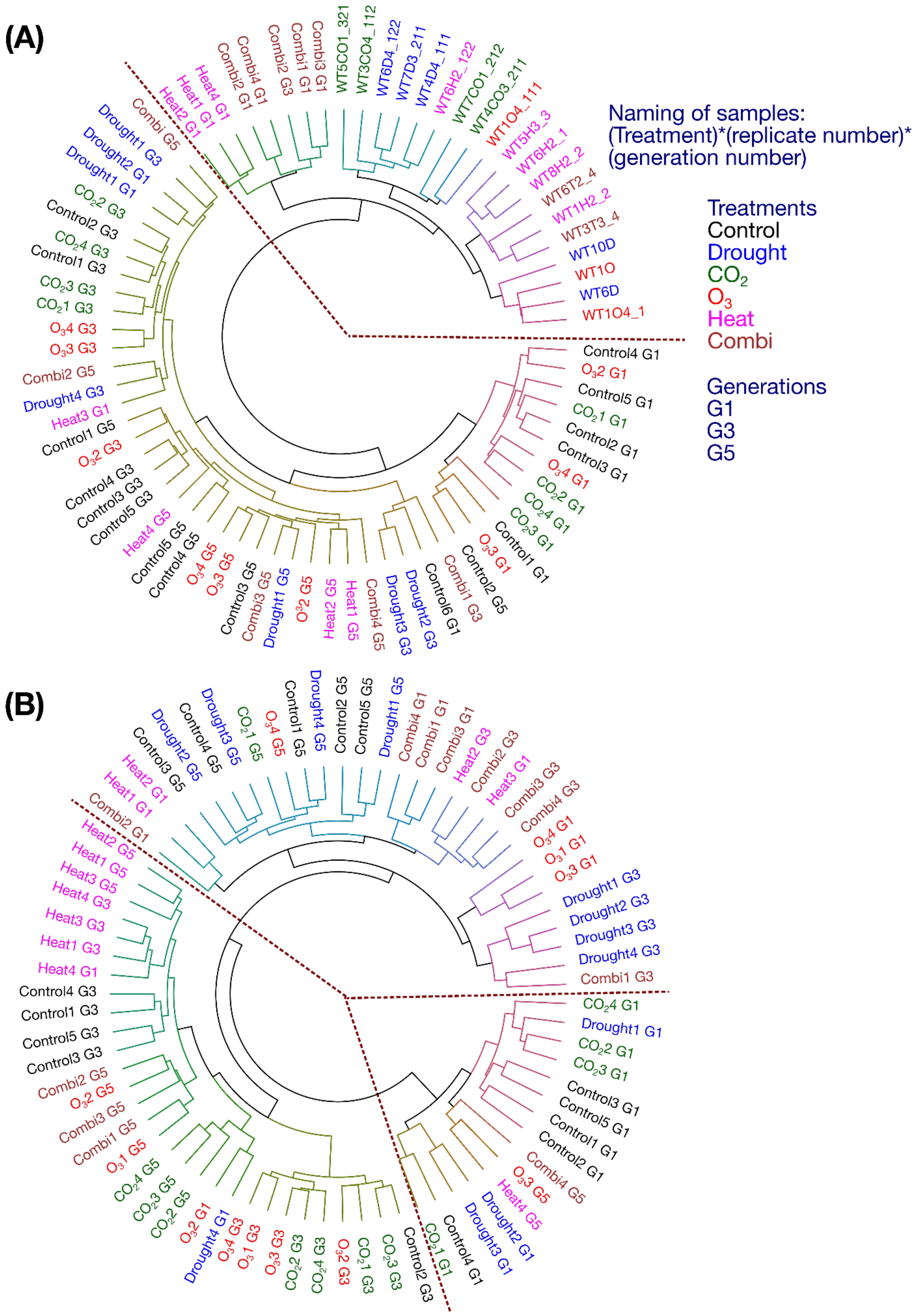

**Supplementary Fig S5. Hierarchical clustering of RNA-seq samples.** Circular dendrograms showing hierarchical clustering of RNA-seq samples from WT and *gsnor-ko* across generations G1, G3, and G5 under Control, Drought, elevated $CO_2$, elevated $O_3$, Heat, and Combination treatments. Sample labels are colour-coded by treatment, and generation is indicated in the sample names. The upper panel shows clustering for WT **(A)**, and the lower panel **(B)** shows clustering for *gsnor-ko*. Red dashed lines indicate the major cluster divisions. These dendrograms illustrate the overall relatedness among samples and the extent to which transcriptomic profiles group by treatment and/or generation.

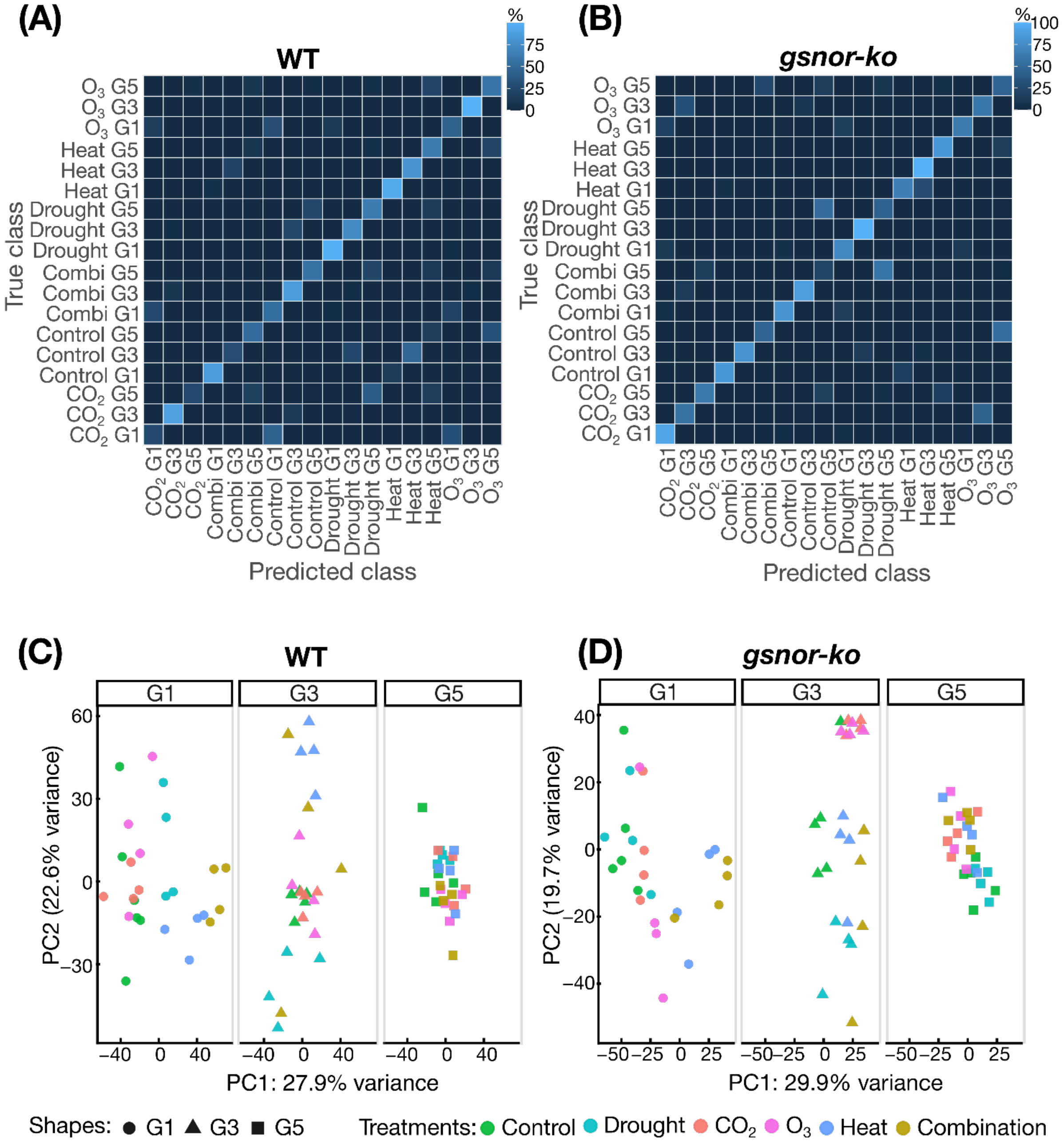


**Supplementary Fig S6. Classification performance and unsupervised transcriptome structure in WT and *gsnor-ko*. (A, B)** Cross-validation confusion matrices for discriminant analysis of principal components (DAPC) performed on WT (**A**) and *gsnor-ko* (**B**) RNA-seq datasets. Values are shown as row-wise percentages, with darker shading indicating lower classification frequency and lighter shading indicating higher classification frequency. Diagonal enrichment indicates correct classification of treatment × generation classes. **(C, D)** Principal component analysis (PCA) of VST-transformed RNA-seq data for WT (**C**) and *gsnor-ko* (**D**), shown separately for generations G1, G3, and G5. Colours indicate treatments (Control, Drought, elevated $CO_2$, elevated $O_3$, Heat, and Combination). Symbols distinguish generations. These plots show the unsupervised distribution of samples in transcriptome space and complement the DAPC-based class separation shown in Fig. 3A.

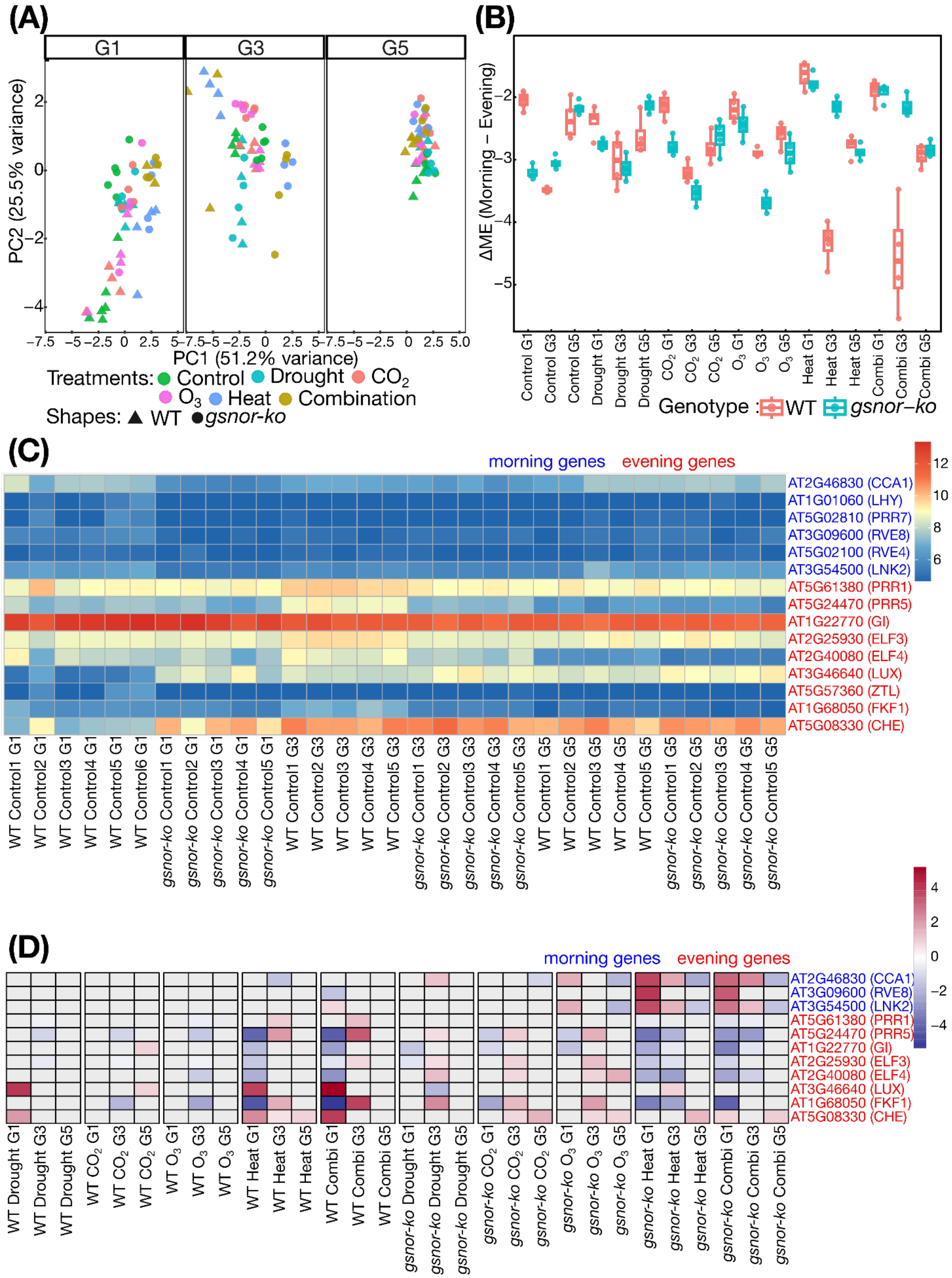


**Supplementary Fig S7. Clock gene analysis across genotypes, treatments, and generations. (A)** Principal component analysis (PCA) based on VST-transformed expression values of selected circadian clock genes in WT and *gsnor-ko* across generations G1, G3, and G5. Colours indicate treatments and symbols indicate treatment classes as shown in the legend.

**(B)** Distribution of the expression metric used to compare morning- and evening-phased clock genes across genotype × generation × treatment combinations. Boxplots show the median, interquartile range, and individual data points for WT and *gsnor-ko*. **(C)** Heatmap of normalised expression values for representative morning- and evening-expressed clock genes across WT and *gsnor-ko* samples from control conditions in G1, G3, and G5. Morning genes are labelled in blue and evening genes in red. **(D)** Heatmap of $\log_2$ fold-change values for selected clock genes across treatments and generations in WT and *gsnor-ko*. Red indicates higher expression and blue indicates lower expression relative to the corresponding control comparison. Together, these analyses show that core clock gene expression patterns remained broadly consistent across generations and support comparable sampling time points for transcriptome analysis.

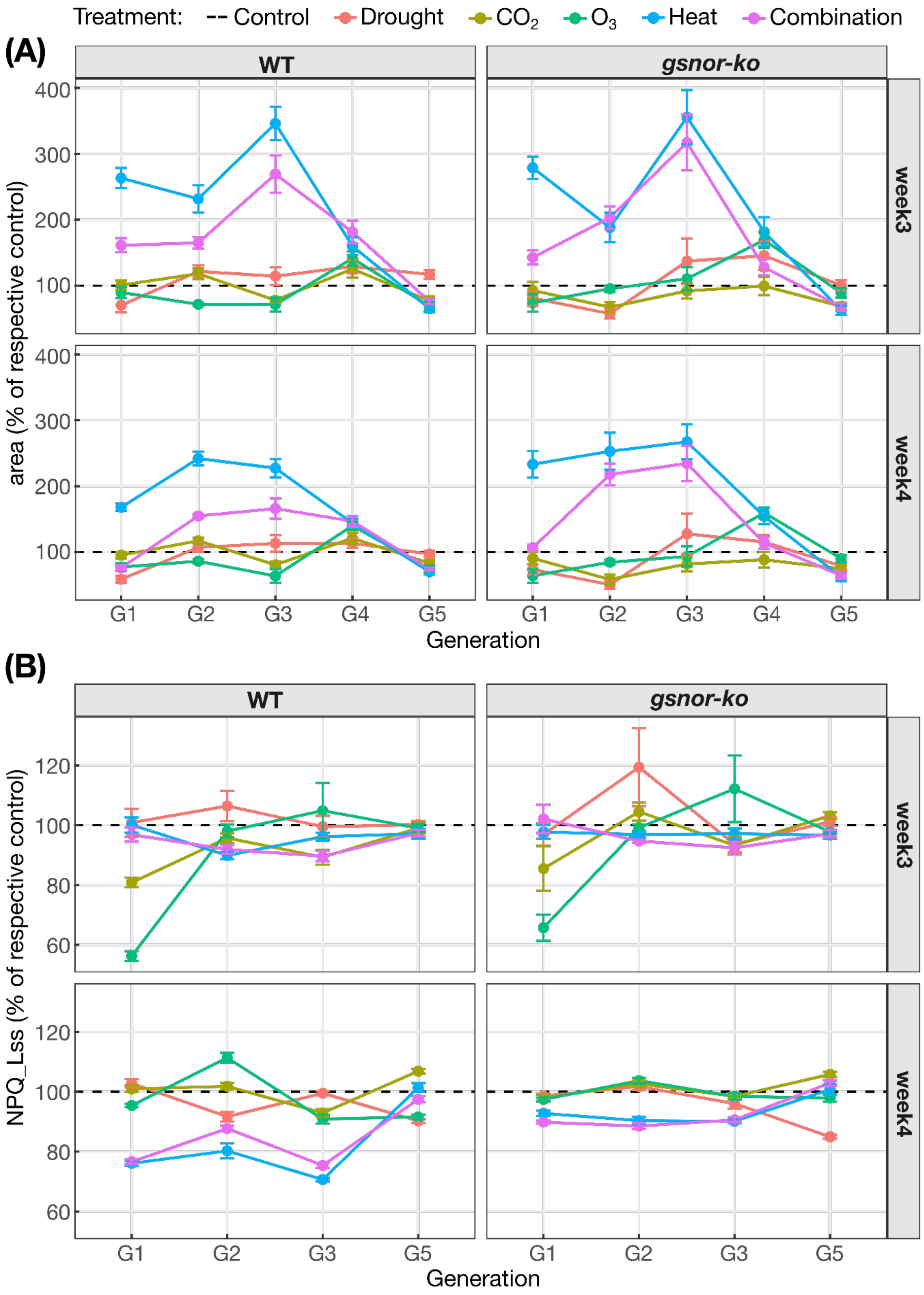


**Supplementary Fig. S8. Normalized growth and photoprotective responses across generations in WT and *gsnor-ko*.**

(A) Rosette area expressed as a percentage of the corresponding same-generation control within each genotype for WT and *gsnor-ko* across G1–G5. (B) NPQ_Lss expressed as a percentage of the corresponding same-generation control within each genotype for WT and *gsnor-ko* across G1, G2, G3, and G5. Values are shown for Drought, $CO_2$, $O_3$, Heat, and

Combination treatments; colours indicate treatment identity. The dashed horizontal line marks 100%, representing the matched control level for each genotype and generation. Points show mean normalized values, and error bars indicate SE. No fluorescence measurements were available for G4 because of technical issues during data acquisition.

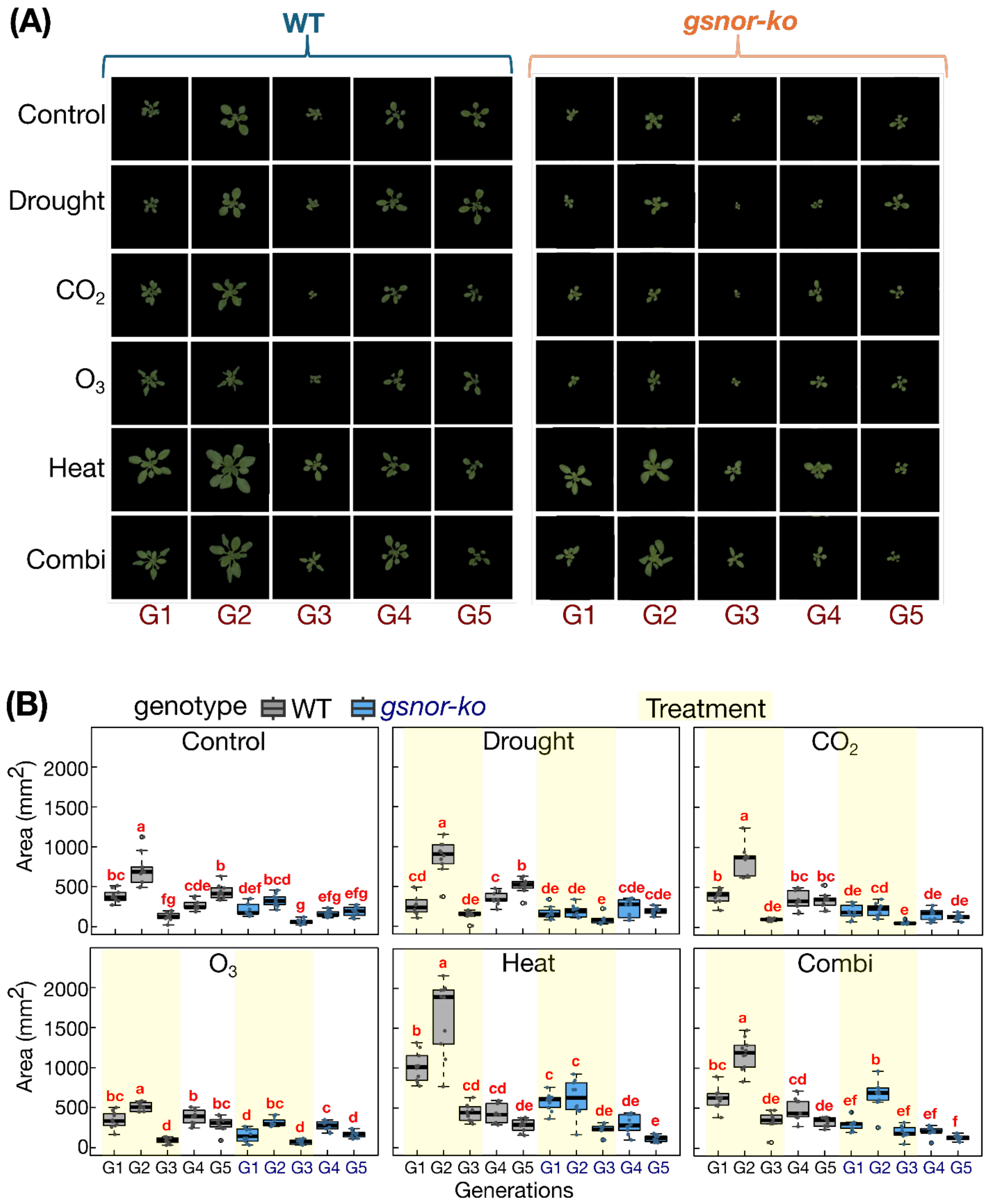


**Supplementary Fig S9. Phenotypic data at week 3. (A)** Representative RGB images of WT and *gsnor-ko* plants grown under Control, Drought, elevated $CO_2$, elevated $O_3$, Heat, and Combination treatments across generations G1-G5 at week 3. **(B)** Rosette area at week 3 for WT and *gsnor-ko* across treatments and generations. Boxplots show the distribution of individual values, with points representing biological replicates. Beige shading indicates generations exposed to stress treatment; G4 and G5 were grown under control conditions after stress withdrawal. Different letters indicate significant differences among genotype × generation groups within each treatment (GLM followed by post hoc multiple comparison, $P < 0.05$).

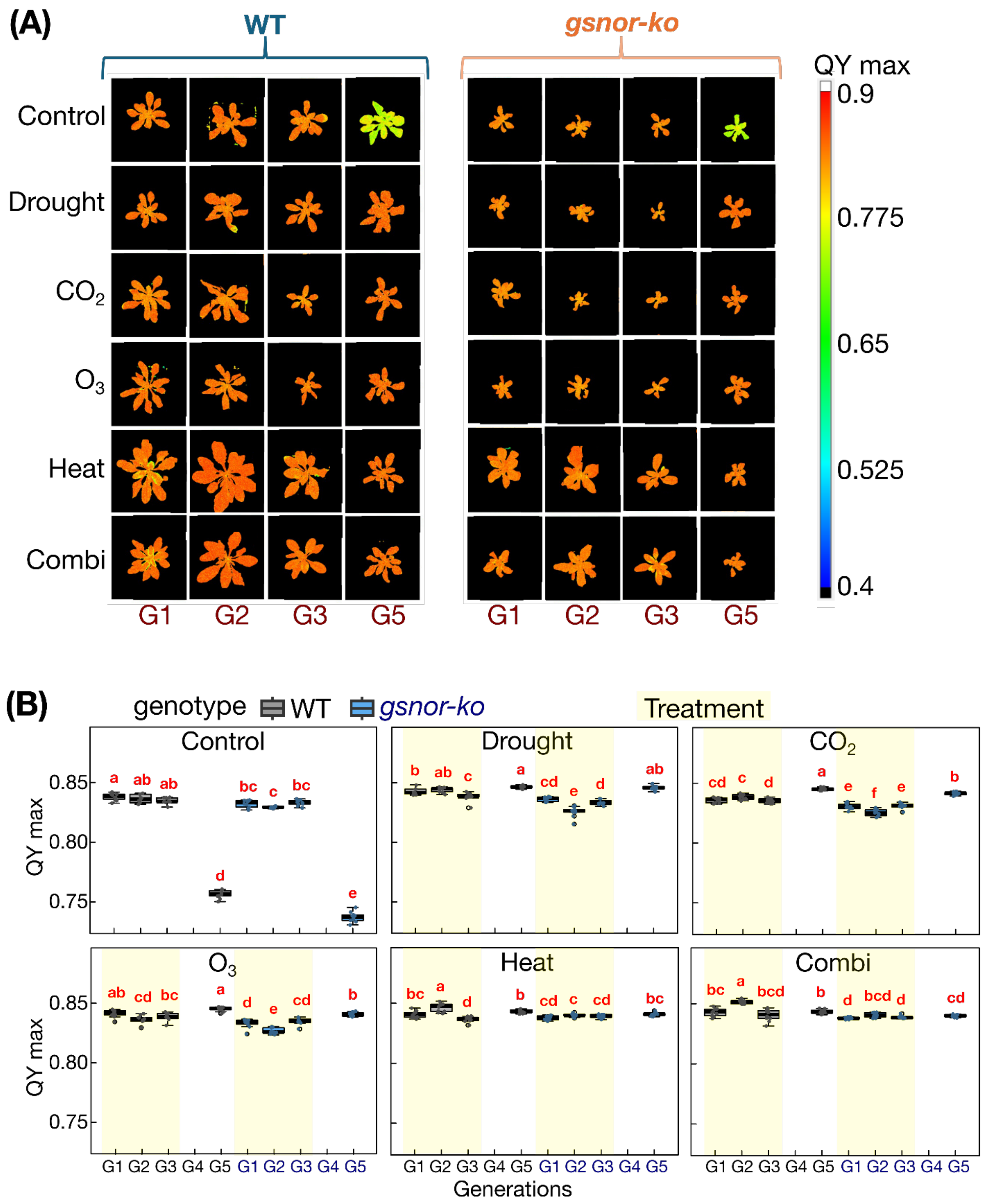


**Supplementary Fig S10. QYmax at week 4 across treatments and generations. (A)** Representative colour images of QYmax in WT and *gsnor-ko* plants grown under Control, Drought, elevated $CO_2$, elevated $O_3$, Heat, and Combination treatments in generations G1, G2, G3, and G5. **(B)** Boxplots showing QYmax values for WT and *gsnor-ko* across treatments and generations at week 4. Points represent biological replicates. Beige shading indicates generations exposed to stress treatment. Different letters indicate significant differences among genotype × generation groups within each treatment (GLM followed by a post hoc multiplecomparison test). $P < 0.05$). QYmax measurements were unavailable for G4 due to technical issues during data acquisition.

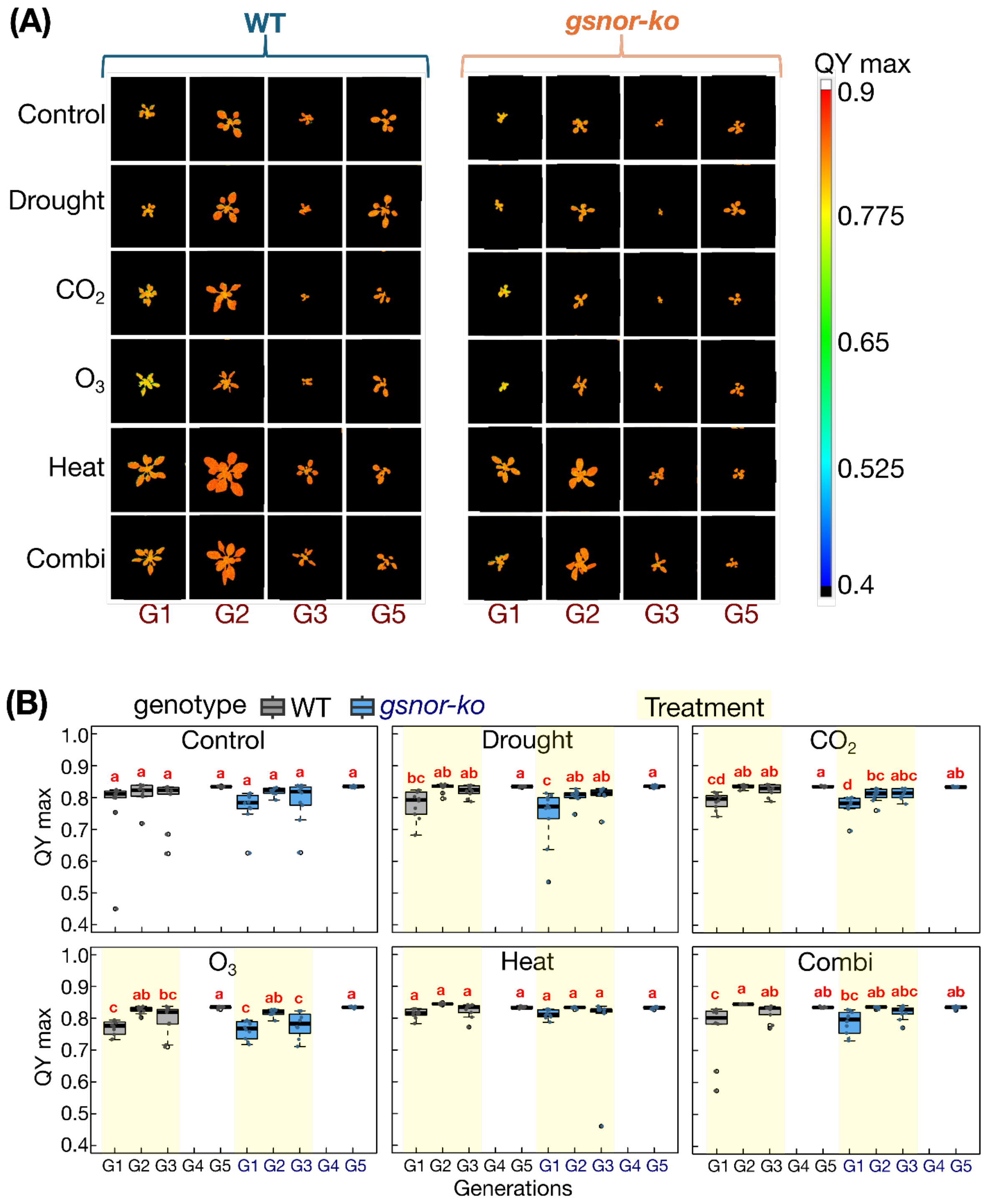


**Supplementary Fig S11. QYmax at week 3 across treatments and generations. (A)** Representative colour images of QYmax in WT and *gsnor-ko* plants grown under Control, Drought, elevated $CO_2$, elevated $O_3$, Heat, and Combination treatments in generations G1, G2, G3, and G5. **(B)** Boxplots showing QYmax values for WT and *gsnor-ko* across treatments and generations at week 3. Points represent biological replicates. Beige shading indicates generations exposed to stress treatment. Different letters indicate significant differences among genotype × generation groups within each treatment (GLM followed by post hoc multiple comparison, $P < 0.05$). QYmax measurements were unavailable for G4 due to technical issues during data acquisition.

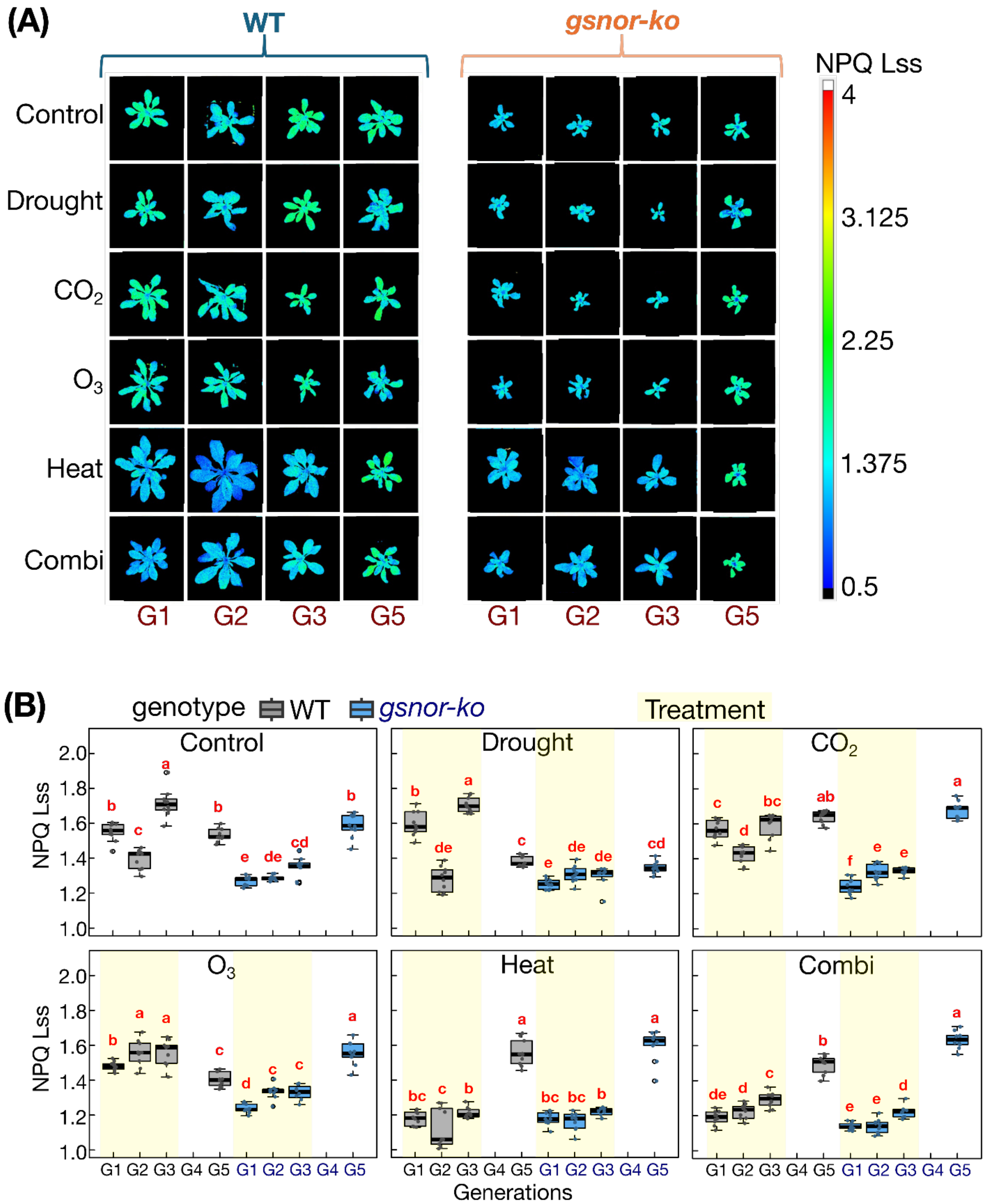


**Supplementary Fig S12. NPQ Lss at week 4 across treatments and generations.** **(A)** Representative colour images of NPQ Lss in WT and *gsnor-ko* plants grown under Control, Drought, elevated $CO_2$, elevated $O_3$, Heat, and Combination treatments in generations G1, G2, G3, and G5. **(B)** Boxplots showing NPQ Lss values for WT and *gsnor-ko* across treatments and generations at week 4. Points represent biological replicates. Beige shading indicates generations exposed to stress treatment. Different letters indicate significant differences among genotype × generation groups within each treatment (GLM followed by post hoc multiple comparison, $P < 0.05$). NPQ Lss measurements were unavailable for G4 due to technical issues during data acquisition.

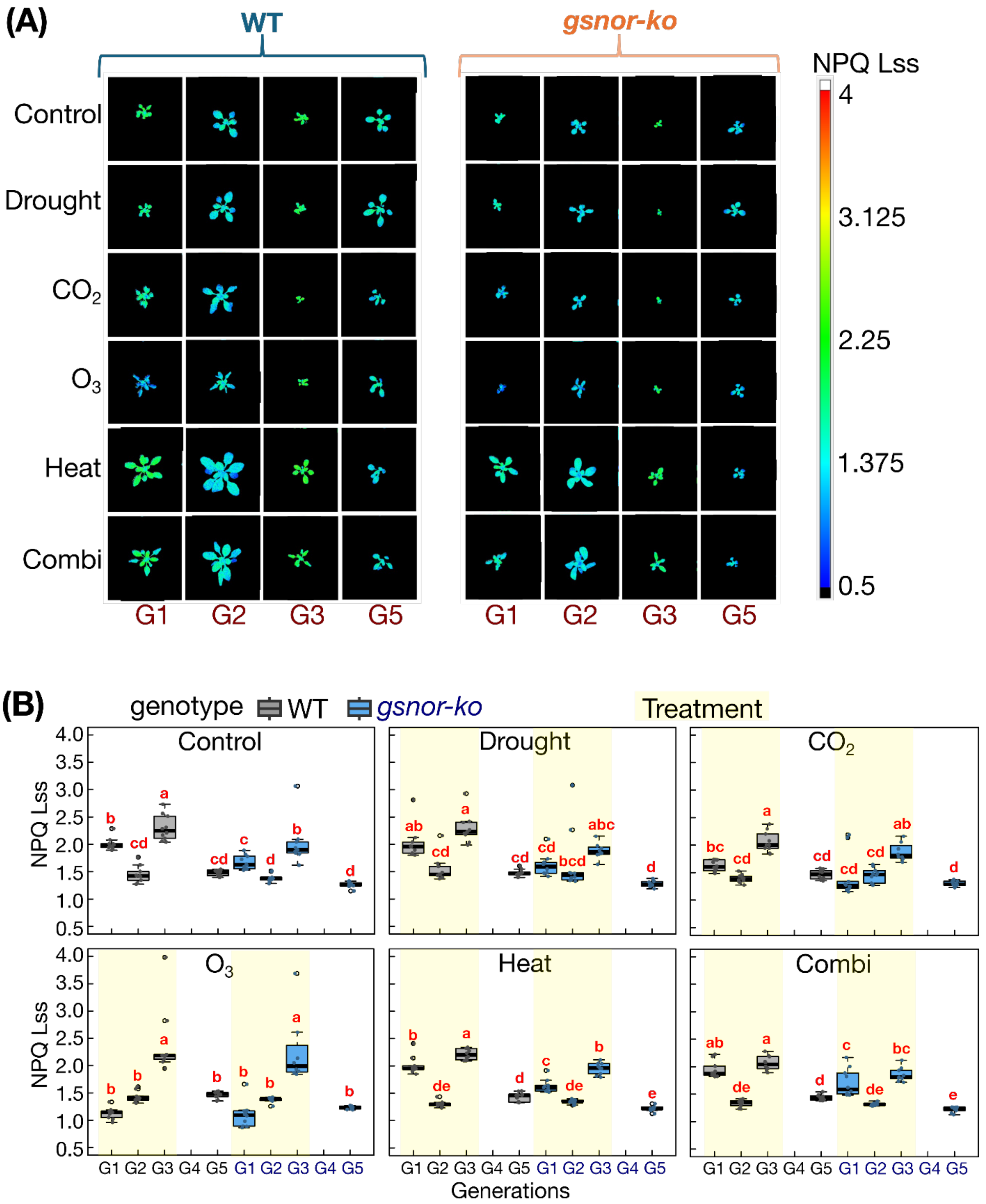


**Supplementary Fig S13. NPQ Lss at week 3 across treatments and generations.** **(A)** Representative colour images of NPQ Lss in WT and *gsnor-ko* plants grown under Control, Drought, elevated $CO_2$, elevated $O_3$, Heat, and Combination treatments in generations G1, G2, G3, and G5. **(B)** Boxplots showing NPQ Lss values for WT and *gsnor-ko* across treatments and generations at week 3. Points represent biological replicates. Beige shading indicates generations exposed to stress treatment. Different letters indicate significant differences among genotype × generation groups within each treatment (GLM followed by post hoc multiple comparison, $P < 0.05$). NPQ Lss measurements were unavailable for G4 due to technical issues during data acquisition.

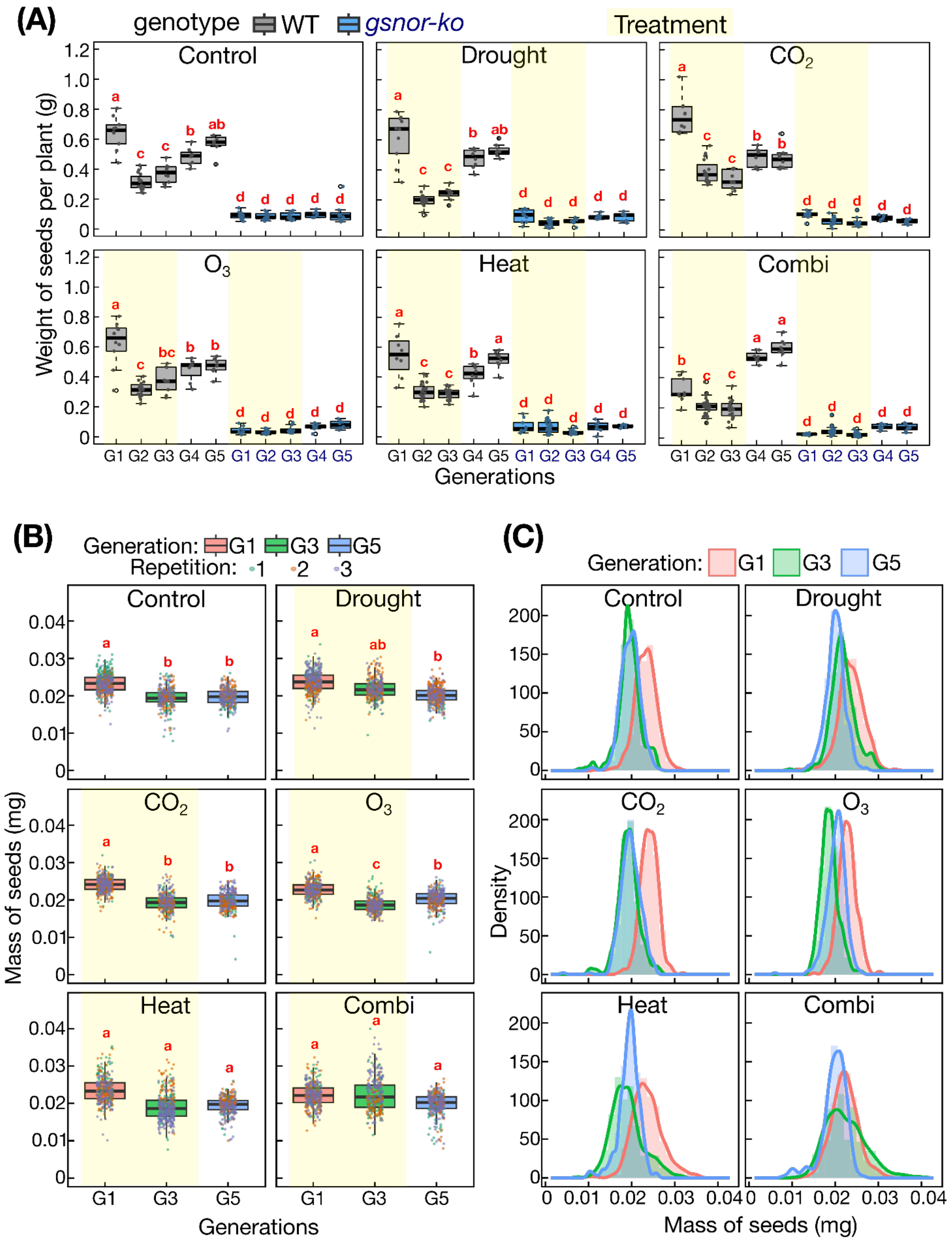


**Supplementary Fig S14. Seed weight and seed phenotyping across treatments and generations. (A)** Seed weight per plant in WT and *gsnor-ko* across Control, Drought, elevated $CO_2$, elevated $O_3$, Heat, and Combination treatments in generations G1-G5. Boxplots show individual biological replicates; WT is shown in grey and *gsnor-ko* in blue. Beige shading indicates generations exposed to stress treatment. Different letters indicate significant differences among genotype × generation groups within each treatment (GLM followed by post hoc multiple comparison, $P < 0.05$). **(B)** Seed mass distribution in WT across treatments and generations G1, G3, and G5. Boxplots show seed mass for individual seeds, with colours indicating generations and points indicating technical repetitions. **(C)** Density distributions of seed mass in WT across treatments and generations G1, G3, and G5. Colours indicate generations. Seed phenotyping in panels **(B, C)** was performed only for WT because insufficient seed material was available for comparable analysis in *gsnor-ko*.

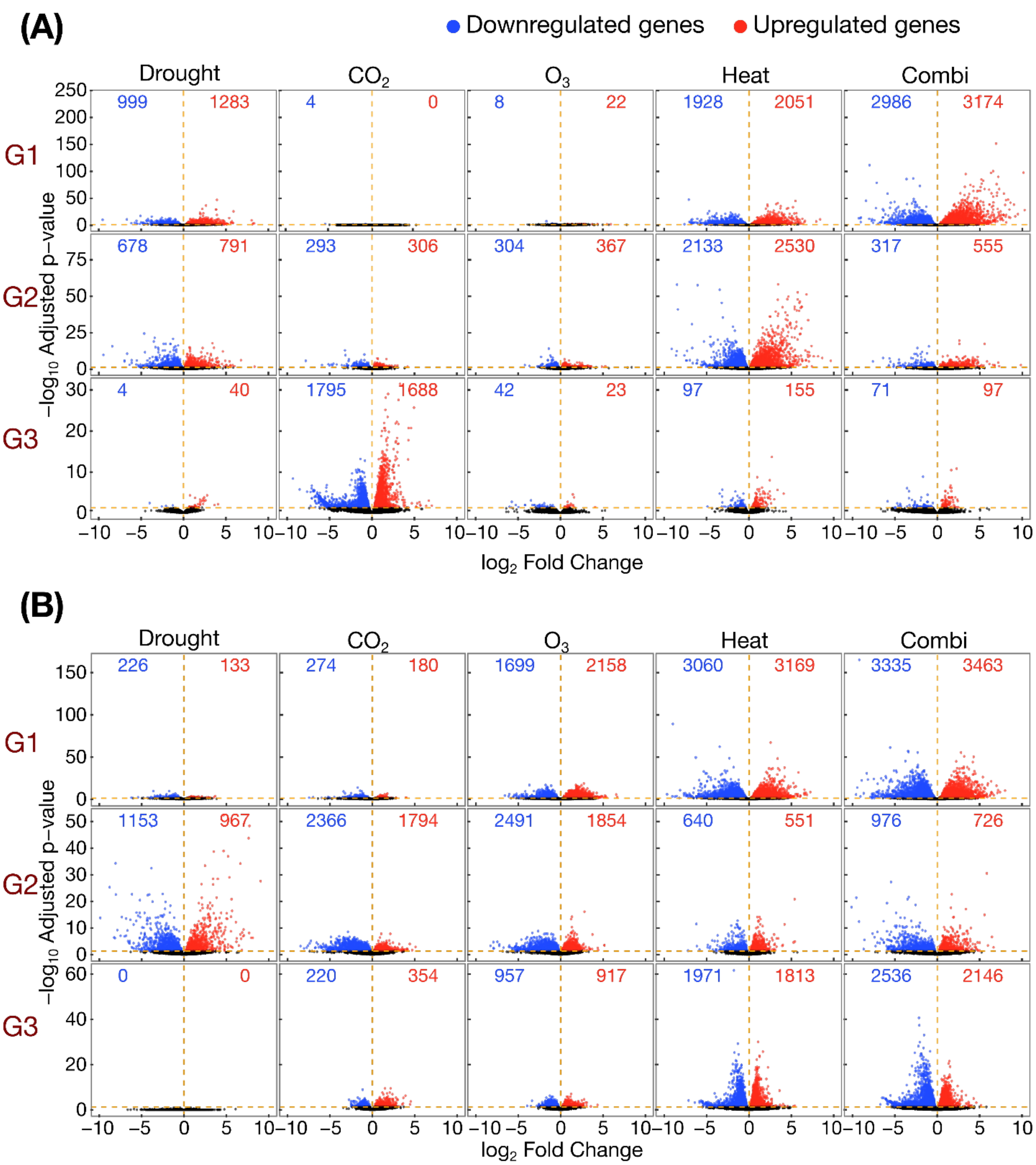


**Supplementary Fig S15. Volcano plots of differential gene expression across treatments and generations in WT and *gsnor-ko*.** Volcano plots showing differentially expressed genes in **(A)** WT and **(B)** *gsnor-ko* for Drought, elevated $CO_2$, elevated $O_3$, Heat, and Combination treatments in generations G1, G3, and G5. Red points indicate up-regulated genes and blue points indicate down-regulated genes. Dashed orange lines indicate the applied significance ($P<0.05$) and fold-change thresholds used to define differentially expressed genes. Numbers in the upper corners of each panel indicate the total numbers of down-regulated (blue) and up-regulated (red) genes for the corresponding contrast.

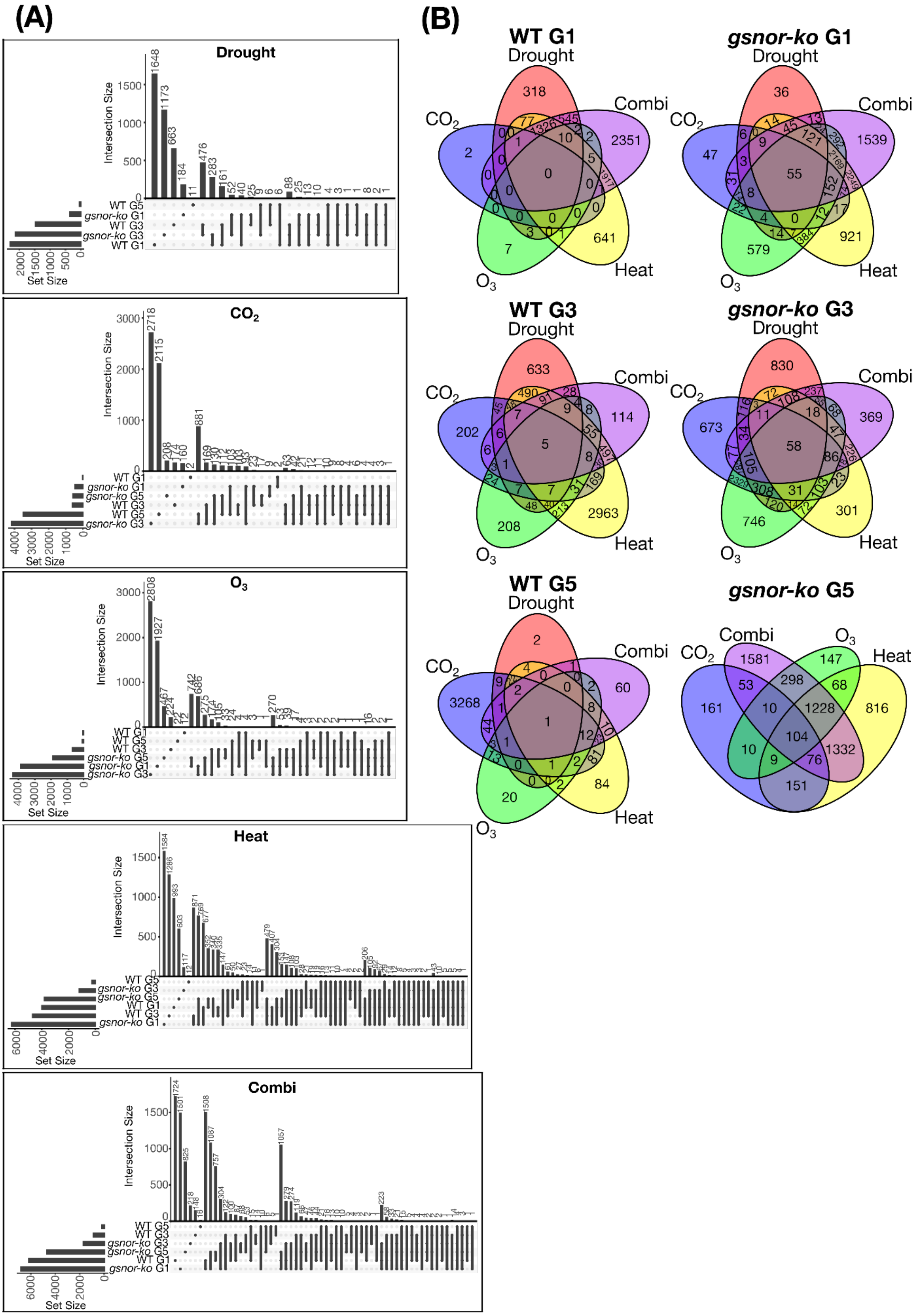

**Supplementary Fig S16. Overlap of differentially expressed genes across treatments, genotypes, and generations. (A)** UpSet plots showing overlap of differentially expressed gene sets between WT and *gsnor-ko* across generations G1, G3, and G5 for each treatment. Separate plots are shown for the elevated $CO_2$, elevated $O_3$, Heat, and Combination treatments. Bar plots indicate intersection sizes and set sizes for the corresponding DEG sets. **(B)** Multi-set Venn diagrams showing overlap of differentially expressed genes among Drought, elevated $CO_2$, elevated $O_3$, Heat, and Combination treatments within each genotype and generation. Separate diagrams are shown for WT G1, *gsnor-ko* G1, WT G3, *gsnor-ko* G3, WT G5, and *gsnor-ko* G5. Numbers indicate unique and shared DEGs among treatments.

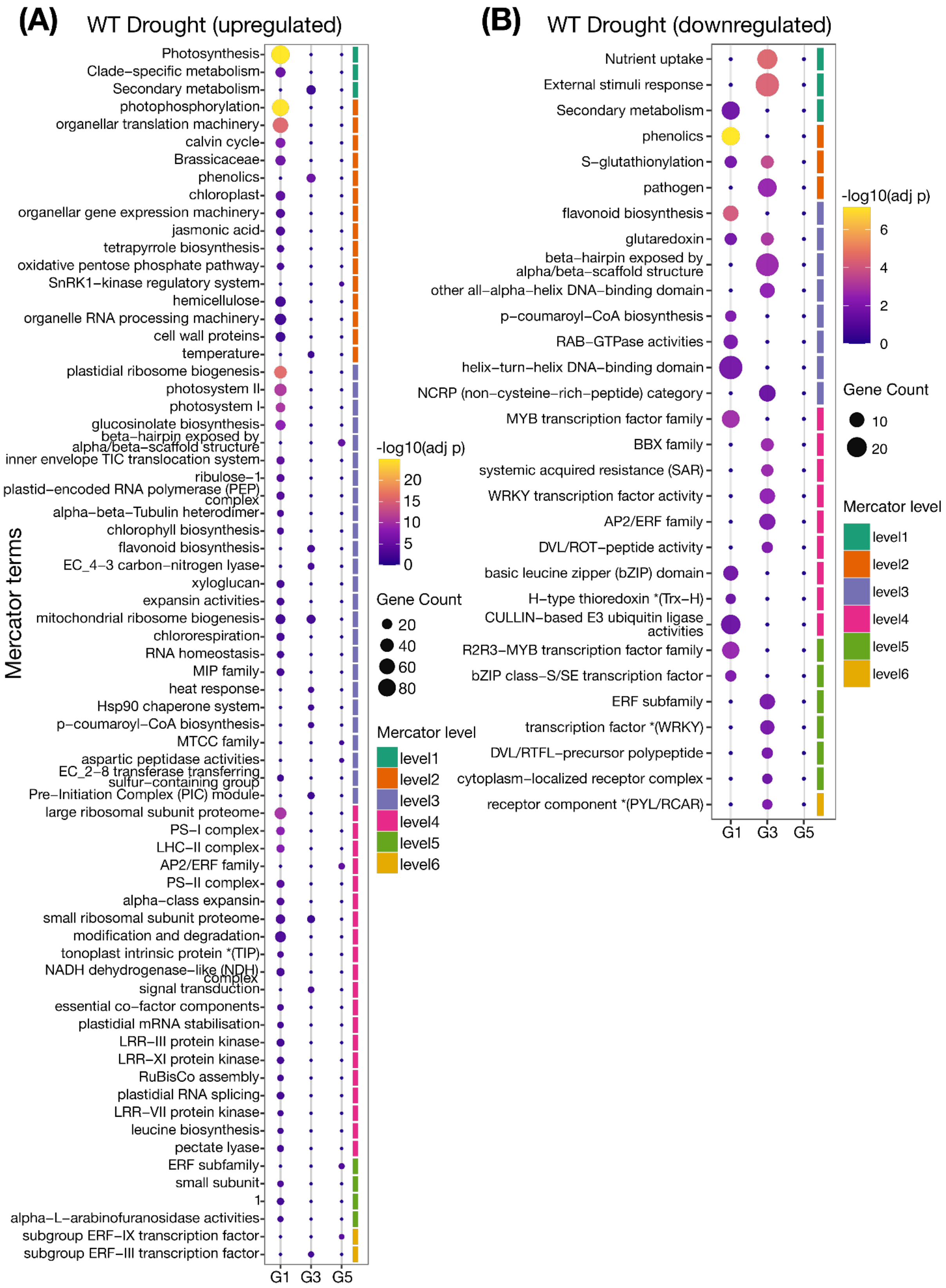

(A)
WT Drought (upregulated)
Mercator terms
Photosynthesis
Clade−specific metabolism
Secondary metabolism
photophosphorylation
organellar translation machinery
calvin cycle
Brassicaceae
phenolics
chloroplast
organellar gene expression machinery
jasmonic acid
tetrapyrrole biosynthesis
oxidative pentose phosphate pathway
SnRK1−kinase regulatory system
hemicellulose
organelle RNA processing machinery
cell wall proteins
temperature
plastidial ribosome biogenesis
photosystem II
photosystem I
glucosinolate biosynthesis
beta−hairpin exposed by alpha/beta−scaffold structure
inner envelope TIC translocation system
ribulose−1
plastid−encoded RNA polymerase (PEP) complex
alpha−beta−Tubulin heterodimer
chlorophyll biosynthesis
flavonoid biosynthesis
EC_4−3 carbon−nitrogen lyase
xyloglucan
expansin activities
mitochondrial ribosome biogenesis
chlororespiration
RNA homeostasis
MIP family
heat response
Hsp90 chaperone system
p−coumaroyl−CoA biosynthesis
MTCC family
aspartic peptidase activities
EC_2−8 transferase transferring sulfur−containing group
Pre−Initiation Complex (PIC) module
large ribosomal subunit proteome
PS−I complex
LHC−II complex
AP2/ERF family
PS−II complex
alpha−class expansin
small ribosomal subunit proteome
modification and degradation
tonoplast intrinsic protein *(TIP)
NADH dehydrogenase−like (NDH) complex
signal transduction
essential co−factor components
plastidial mRNA stabilisation
LRR−III protein kinase
LRR−XI protein kinase
RuBisCo assembly
plastidial RNA splicing
LRR−VII protein kinase
leucine biosynthesis
pectate lyase
ERF subfamily
small subunit
1
alpha−L−arabinofuranosidase activities
subgroup ERF−IX transcription factor
subgroup ERF−III transcription factor
G1 G3 G5
-log10(adj p)
20
15
10
5
0
Gene Count
20
40
60
80
Mercator level
level1
level2
level3
level4
level5
level6
(B)
WT Drought (downregulated)
Nutrient uptake
External stimuli response
Secondary metabolism
phenolics
S−glutathionylation
pathogen
flavonoid biosynthesis
glutaredoxin
beta−hairpin exposed by alpha/beta−scaffold structure
other all−alpha−helix DNA−binding domain
p−coumaroyl−CoA biosynthesis
RAB−GTPase activities
helix−turn−helix DNA−binding domain
NCRP (non−cysteine−rich−peptide) category
MYB transcription factor family
BBX family
systemic acquired resistance (SAR)
WRKY transcription factor activity
AP2/ERF family
DVL/ROT−peptide activity
basic leucine zipper (bZIP) domain
H−type thioredoxin *(Trx−H)
CULLIN−based E3 ubiquitin ligase activities
R2R3−MYB transcription factor family
bZIP class−S/SE transcription factor
ERF subfamily
transcription factor *(WRKY)
DVL/RTFL−precursor polypeptide
cytoplasm−localized receptor complex
receptor component *(PYL/RCAR)
G1 G3 G5
-log10(adj p)
6
4
2
0
Gene Count
10
20
Mercator level
level1
level2
level3
level4
level5
level6

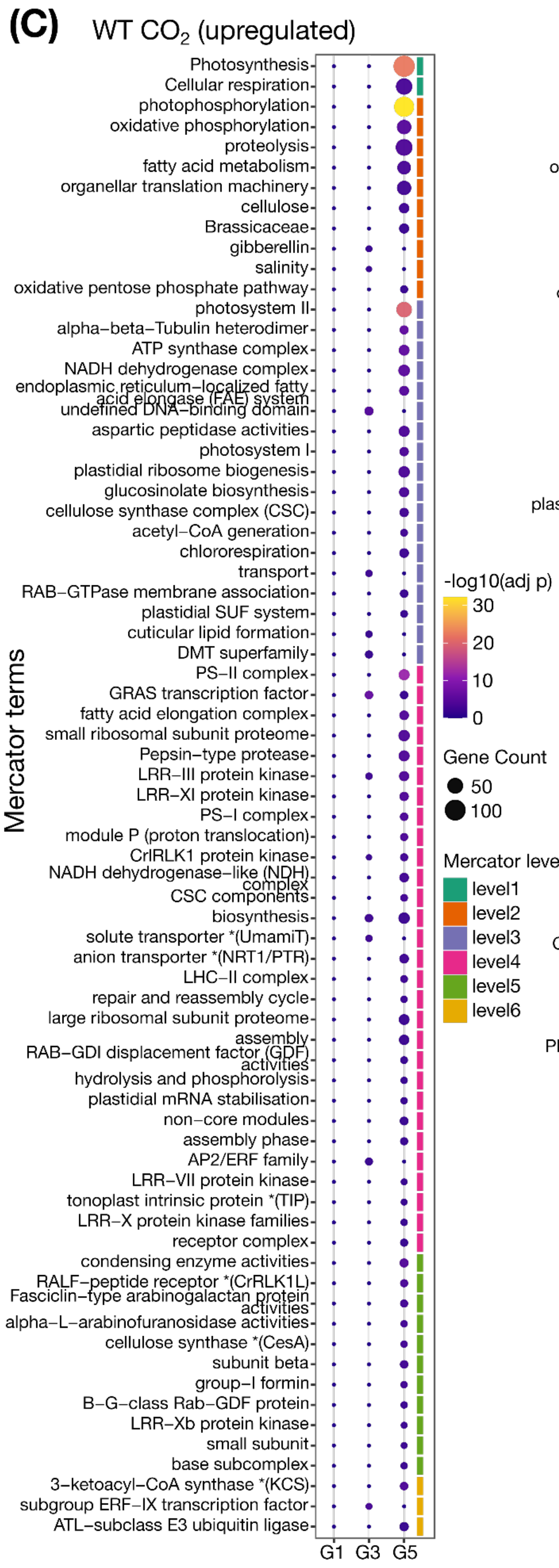

(C)
WT $CO_2$ (upregulated)
Mercator terms
Photosynthesis
Cellular respiration
photophosphorylation
oxidative phosphorylation
proteolysis
fatty acid metabolism
organellar translation machinery
cellulose
Brassicaceae
gibberellin
salinity
oxidative pentose phosphate pathway
photosystem II
alpha−beta−Tubulin heterodimer
ATP synthase complex
NADH dehydrogenase complex
endoplasmic reticulum−localized fatty acid elongase (FAE) system
undefined DNA−binding domain
aspartic peptidase activities
photosystem I
plastidial ribosome biogenesis
glucosinolate biosynthesis
cellulose synthase complex (CSC)
acetyl−CoA generation
chlororespiration
transport
RAB−GTPase membrane association
plastidial SUF system
cuticular lipid formation
DMT superfamily
PS−II complex
GRAS transcription factor
fatty acid elongation complex
small ribosomal subunit proteome
Pepsin−type protease
LRR−III protein kinase
LRR−XI protein kinase
PS−I complex
module P (proton translocation)
CrRLK1 protein kinase
NADH dehydrogenase−like (NDH) complex
CSC components
biosynthesis
solute transporter *(UmamiT)
anion transporter *(NRT1/PTR)
LHC−II complex
repair and reassembly cycle
large ribosomal subunit proteome
assembly
RAB−GDI displacement factor (GDF) activities
hydrolysis and phosphorolysis
plastidial mRNA stabilisation
non−core modules
assembly phase
AP2/ERF family
LRR−VII protein kinase
tonoplast intrinsic protein *(TIP)
LRR−X protein kinase families
receptor complex
condensing enzyme activities
RALF−peptide receptor *(CrRLK1L)
Fasciclin−type arabinogalactan protein activities
alpha−L−arabinofuranosidase activities
cellulose synthase *(CesA)
subunit beta
group−I formin
B−G−class Rab−GDF protein
LRR−Xb protein kinase
small subunit
base subcomplex
3−ketoacyl−CoA synthase *(KCS)
subgroup ERF−IX transcription factor
ATL−subclass E3 ubiquitin ligase
G1
G3
G5
-log10(adj p)
30
20
10
0
Gene Count
50
100
Mercator level
level1
level2
level3
level4
level5
level6


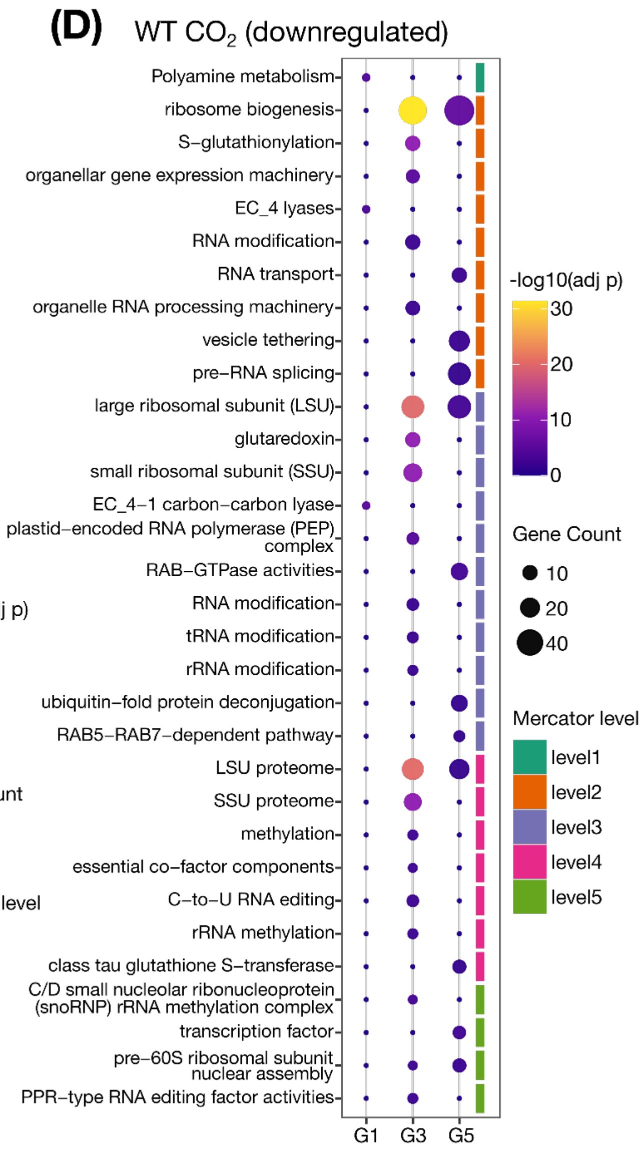

(D)
WT $CO_2$ (downregulated)
Polyamine metabolism
ribosome biogenesis
S−glutathionylation
organellar gene expression machinery
EC_4 lyases
RNA modification
RNA transport
organelle RNA processing machinery
vesicle tethering
pre−RNA splicing
large ribosomal subunit (LSU)
glutaredoxin
small ribosomal subunit (SSU)
EC_4−1 carbon−carbon lyase
plastid−encoded RNA polymerase (PEP) complex
RAB−GTPase activities
RNA modification
tRNA modification
rRNA modification
ubiquitin−fold protein deconjugation
RAB5−RAB7−dependent pathway
LSU proteome
SSU proteome
methylation
essential co−factor components
C−to−U RNA editing
rRNA methylation
class tau glutathione S−transferase
C/D small nucleolar ribonucleoprotein (snoRNP) rRNA methylation complex
transcription factor
pre−60S ribosomal subunit nuclear assembly
PPR−type RNA editing factor activities
G1
G3
G5
-log10(adj p)
30
20
10
0
Gene Count
10
20
40
Mercator level
level1
level2
level3
level4
level5

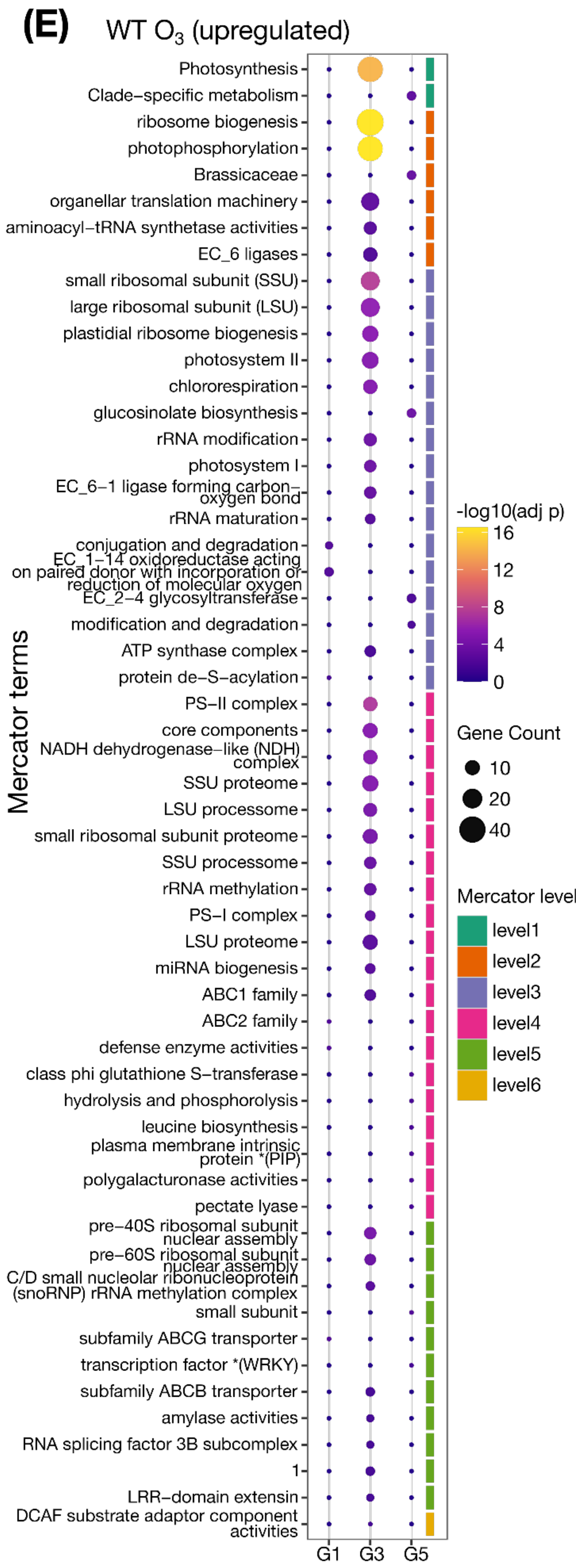

(E)
WT $O_3$ (upregulated)
Mercator terms
Photosynthesis
Clade–specific metabolism
ribosome biogenesis
photophosphorylation
Brassicaceae
organellar translation machinery
aminoacyl–tRNA synthetase activities
EC_6 ligases
small ribosomal subunit (SSU)
large ribosomal subunit (LSU)
plastidial ribosome biogenesis
photosystem II
chlororespiration
glucosinolate biosynthesis
rRNA modification
photosystem I
EC_6–1 ligase forming carbon–oxygen bond
rRNA maturation
conjugation and degradation
EC_1–14 oxidoreductase acting on paired donor with incorporation or reduction of molecular oxygen
EC_2–4 glycosyltransferase
modification and degradation
ATP synthase complex
protein de–S–acylation
PS–II complex
core components
NADH dehydrogenase–like (NDH) complex
SSU proteome
LSU processome
small ribosomal subunit proteome
SSU processome
rRNA methylation
PS–I complex
LSU proteome
miRNA biogenesis
ABC1 family
ABC2 family
defense enzyme activities
class phi glutathione S–transferase
hydrolysis and phosphorolysis
leucine biosynthesis
plasma membrane intrinsic protein *(PIP)
polygalacturonase activities
pectate lyase
pre–40S ribosomal subunit nuclear assembly
pre–60S ribosomal subunit nuclear assembly
C/D small nucleolar ribonucleoprotein (snoRNP) rRNA methylation complex
small subunit
subfamily ABCG transporter
transcription factor *(WRKY)
subfamily ABCB transporter
amylase activities
RNA splicing factor 3B subcomplex
1
LRR–domain extensin
DCAF substrate adaptor component activities
G1
G3
G5
-log10(adj p)
16
12
8
4
0
Gene Count
10
20
40
Mercator level
level1
level2
level3
level4
level5
level6


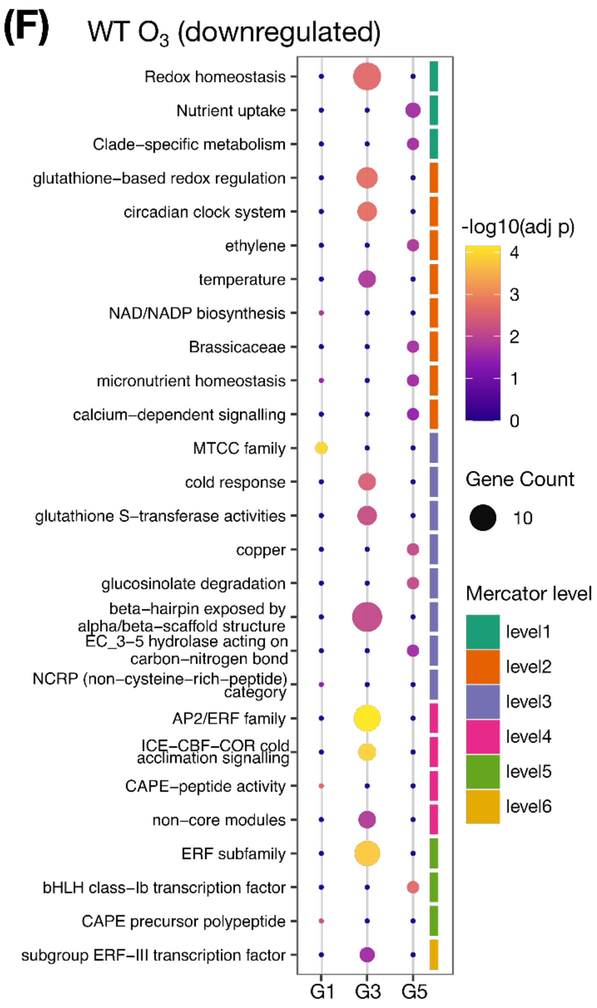

(F)
WT $O_3$ (downregulated)
Redox homeostasis
Nutrient uptake
Clade–specific metabolism
glutathione–based redox regulation
circadian clock system
ethylene
temperature
NAD/NADP biosynthesis
Brassicaceae
micronutrient homeostasis
calcium–dependent signalling
MTCC family
cold response
glutathione S–transferase activities
copper
glucosinolate degradation
beta–hairpin exposed by alpha/beta–scaffold structure
EC_3–5 hydrolase acting on carbon–nitrogen bond
NCRP (non–cysteine–rich–peptide) category
AP2/ERF family
ICE–CBF–COR cold acclimation signalling
CAPE–peptide activity
non–core modules
ERF subfamily
bHLH class–Ib transcription factor
CAPE precursor polypeptide
subgroup ERF–III transcription factor
G1
G3
G5
-log10(adj p)
4
3
2
1
0
Gene Count
10
Mercator level
level1
level2
level3
level4
level5
level6

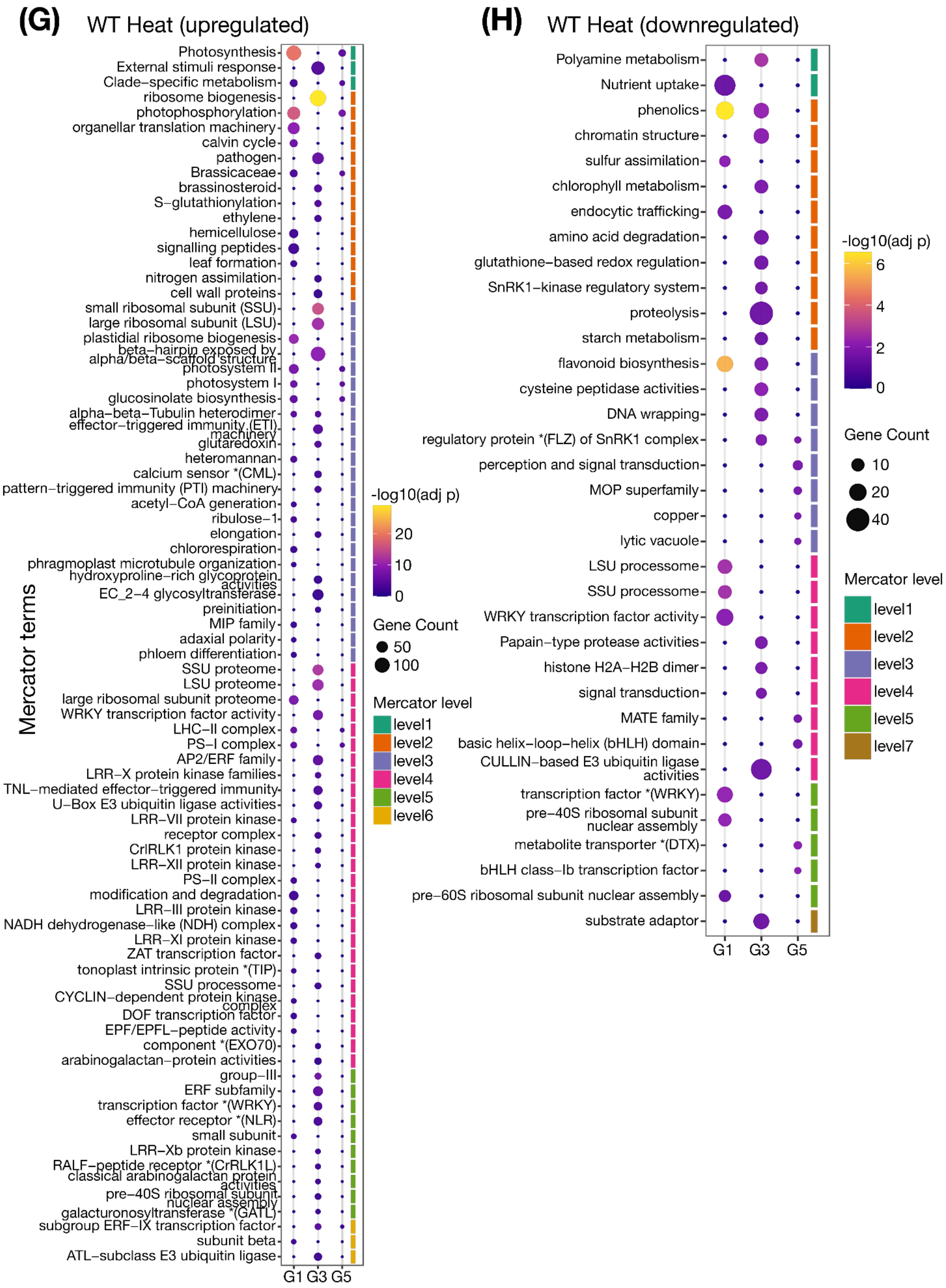

(G)
WT Heat (upregulated)
Mercator terms
Photosynthesis
External stimuli response
Clade−specific metabolism
ribosome biogenesis
photophosphorylation
organellar translation machinery
calvin cycle
pathogen
Brassicaceae
brassinosteroid
S−glutathionylation
ethylene
hemicellulose
signalling peptides
leaf formation
nitrogen assimilation
cell wall proteins
small ribosomal subunit (SSU)
large ribosomal subunit (LSU)
plastidial ribosome biogenesis
beta−hairpin exposed by alpha/beta−scaffold structure
photosystem II
photosystem I
glucosinolate biosynthesis
alpha−beta−Tubulin heterodimer
effector−triggered immunity (ETI) machinery
glutaredoxin
heteromannan
calcium sensor *(CML)
pattern−triggered immunity (PTI) machinery
acetyl−CoA generation
ribulose−1
elongation
chlororespiration
phragmoplast microtubule organization
hydroxyproline−rich glycoprotein activities
EC_2−4 glycosyltransferase
preinitiation
MIP family
adaxial polarity
phloem differentiation
SSU proteome
LSU proteome
large ribosomal subunit proteome
WRKY transcription factor activity
LHC−II complex
PS−I complex
AP2/ERF family
LRR−X protein kinase families
TNL−mediated effector−triggered immunity
U−Box E3 ubiquitin ligase activities
LRR−VII protein kinase
receptor complex
CrRLK1 protein kinase
LRR−XII protein kinase
PS−II complex
modification and degradation
LRR−III protein kinase
NADH dehydrogenase−like (NDH) complex
LRR−XI protein kinase
ZAT transcription factor
tonoplast intrinsic protein *(TIP)
SSU processome
CYCLIN−dependent protein kinase complex
DOF transcription factor
EPF/EPFL−peptide activity
component *(EXO70)
arabinogalactan−protein activities
group−III
ERF subfamily
transcription factor *(WRKY)
effector receptor *(NLR)
small subunit
LRR−Xb protein kinase
RALF−peptide receptor *(CrRLK1L)
classical arabinogalactan protein activities
pre−40S ribosomal subunit nuclear assembly
galacturonosyltransferase *(GATL)
subgroup ERF−IX transcription factor
subunit beta
ATL−subclass E3 ubiquitin ligase
G1 G3 G5
-log10(adj p)
20
10
0
Gene Count
50
100
Mercator level
level1
level2
level3
level4
level5
level6
(H)
WT Heat (downregulated)
Polyamine metabolism
Nutrient uptake
phenolics
chromatin structure
sulfur assimilation
chlorophyll metabolism
endocytic trafficking
amino acid degradation
glutathione−based redox regulation
SnRK1−kinase regulatory system
proteolysis
starch metabolism
flavonoid biosynthesis
cysteine peptidase activities
DNA wrapping
regulatory protein *(FLZ) of SnRK1 complex
perception and signal transduction
MOP superfamily
copper
lytic vacuole
LSU processome
SSU processome
WRKY transcription factor activity
Papain−type protease activities
histone H2A−H2B dimer
signal transduction
MATE family
basic helix−loop−helix (bHLH) domain
CULLIN−based E3 ubiquitin ligase activities
transcription factor *(WRKY)
pre−40S ribosomal subunit nuclear assembly
metabolite transporter *(DTX)
bHLH class−Ib transcription factor
pre−60S ribosomal subunit nuclear assembly
substrate adaptor
G1 G3 G5
-log10(adj p)
6
4
2
0
Gene Count
10
20
40
Mercator level
level1
level2
level3
level4
level5
level7

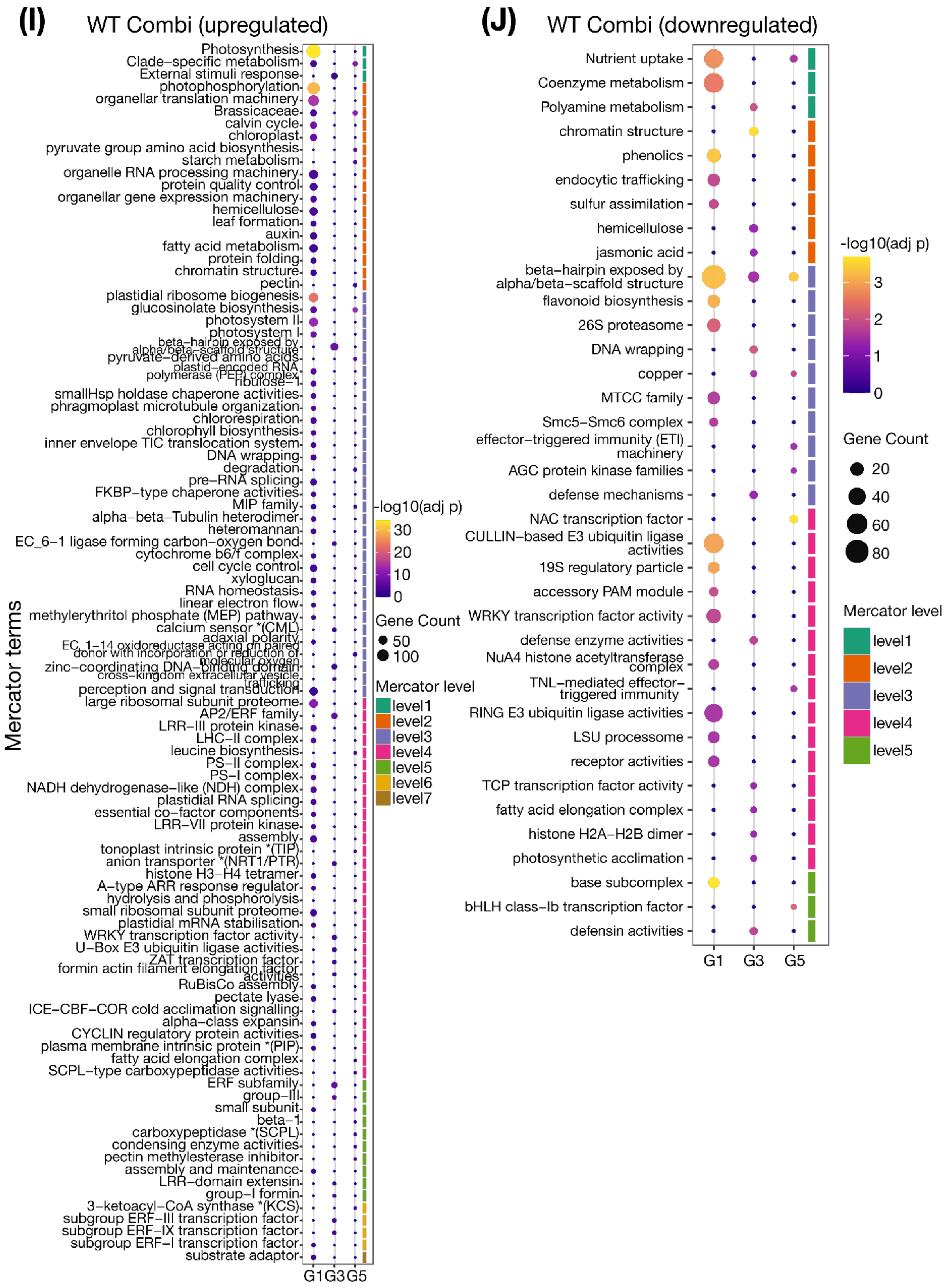

(I)
WT Combi (upregulated)
Mercator terms
Photosynthesis
Clade−specific metabolism
External stimuli response
photophosphorylation
organellar translation machinery
Brassicaceae
calvin cycle
chloroplast
pyruvate group amino acid biosynthesis
starch metabolism
organelle RNA processing machinery
protein quality control
organellar gene expression machinery
hemicellulose
leaf formation
auxin
fatty acid metabolism
protein folding
chromatin structure
pectin
plastidial ribosome biogenesis
glucosinolate biosynthesis
photosystem II
photosystem I
ribulose−1
smallHsp holdase chaperone activities
phragmoplast microtubule organization
chlororespiration
chlorophyll biosynthesis
inner envelope TIC translocation system
DNA wrapping
degradation
pre−RNA splicing
FKBP−type chaperone activities
MIP family
alpha−beta−Tubulin heterodimer
heteromannan
EC_6−1 ligase forming carbon−oxygen bond
cytochrome b6/f complex
cell cycle control
xyloglucan
RNA homeostasis
linear electron flow
methylerythritol phosphate (MEP) pathway
calcium sensor *(CML)
perception and signal transduction
large ribosomal subunit proteome
AP2/ERF family
LRR−III protein kinase
LHC−II complex
leucine biosynthesis
PS−II complex
PS−I complex
NADH dehydrogenase−like (NDH) complex
plastidial RNA splicing
essential co−factor components
LRR−VII protein kinase
assembly
tonoplast intrinsic protein *(TIP)
anion transporter *(NRT1/PTR)
histone H3−H4 tetramer
A−type ARR response regulator
hydrolysis and phosphorolysis
small ribosomal subunit proteome
plastidial mRNA stabilisation
WRKY transcription factor activity
U−Box E3 ubiquitin ligase activities
RuBisCo assembly
pectate lyase
ICE−CBF−COR cold acclimation signalling
alpha−class expansin
CYCLIN regulatory protein activities
plasma membrane intrinsic protein *(PIP)
fatty acid elongation complex
SCPL−type carboxypeptidase activities
ERF subfamily
group−III
small subunit
beta−1
carboxypeptidase *(SCPL)
condensing enzyme activities
pectin methylesterase inhibitor
assembly and maintenance
LRR−domain extensin
group−I formin
3−ketoacyl−CoA synthase *(KCS)
subgroup ERF−III transcription factor
subgroup ERF−IX transcription factor
subgroup ERF−I transcription factor
substrate adaptor
G1 G3 G5
-log10(adj p)
30
20
10
0
Gene Count
50
100
Mercator level
level1
level2
level3
level4
level5
level6
level7
(J)
WT Combi (downregulated)
Nutrient uptake
Coenzyme metabolism
Polyamine metabolism
chromatin structure
phenolics
endocytic trafficking
sulfur assimilation
hemicellulose
jasmonic acid
beta−hairpin exposed by alpha/beta−scaffold structure
flavonoid biosynthesis
26S proteasome
DNA wrapping
copper
MTCC family
Smc5−Smc6 complex
effector−triggered immunity (ETI) machinery
AGC protein kinase families
defense mechanisms
NAC transcription factor
CULLIN−based E3 ubiquitin ligase activities
19S regulatory particle
accessory PAM module
WRKY transcription factor activity
defense enzyme activities
NuA4 histone acetyltransferase complex
TNL−mediated effector−triggered immunity
RING E3 ubiquitin ligase activities
LSU processome
receptor activities
TCP transcription factor activity
fatty acid elongation complex
histone H2A−H2B dimer
photosynthetic acclimation
base subcomplex
bHLH class−Ib transcription factor
defensin activities
G1 G3 G5
-log10(adj p)
3
2
1
0
Gene Count
20
40
60
80
Mercator level
level1
level2
level3
level4
level5

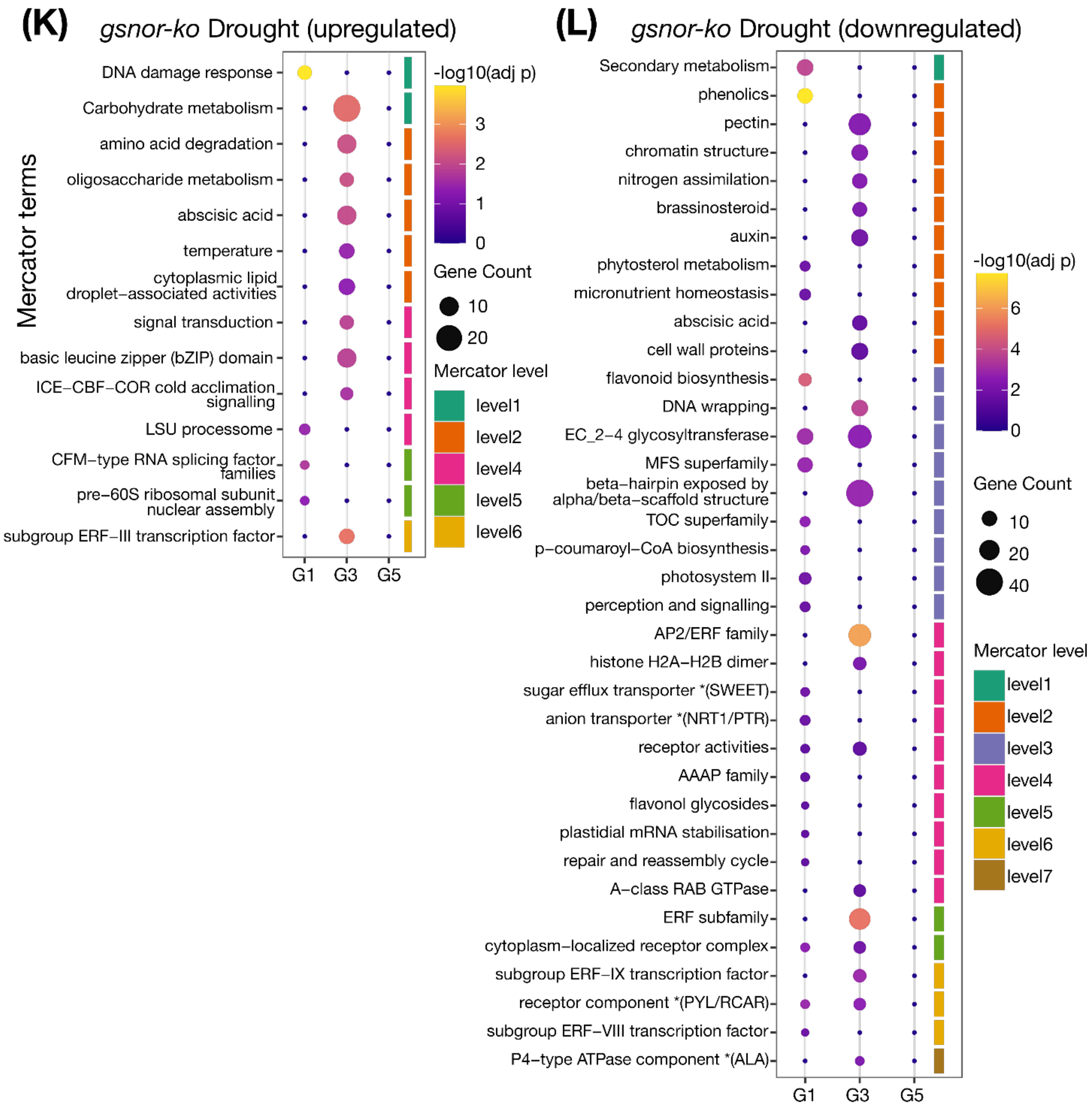
(K) gsnor-ko Drought (upregulated)
Mercator terms
DNA damage response
Carbohydrate metabolism
amino acid degradation
oligosaccharide metabolism
abscisic acid
temperature
cytoplasmic lipid droplet–associated activities
signal transduction
basic leucine zipper (bZIP) domain
ICE–CBF–COR cold acclimation signalling
LSU processome
CFM–type RNA splicing factor families
pre–60S ribosomal subunit nuclear assembly
subgroup ERF–III transcription factor
G1
G3
G5
-log10(adj p)
3
2
1
0
Gene Count
10
20
Mercator level
level1
level2
level4
level5
level6
(L) gsnor-ko Drought (downregulated)
Secondary metabolism
phenolics
pectin
chromatin structure
nitrogen assimilation
brassinosteroid
auxin
phytosterol metabolism
micronutrient homeostasis
abscisic acid
cell wall proteins
flavonoid biosynthesis
DNA wrapping
EC_2–4 glycosyltransferase
MFS superfamily
beta–hairpin exposed by alpha/beta–scaffold structure
TOC superfamily
p–coumaroyl–CoA biosynthesis
photosystem II
perception and signalling
AP2/ERF family
histone H2A–H2B dimer
sugar efflux transporter *(SWEET)
anion transporter *(NRT1/PTR)
receptor activities
AAAP family
flavonol glycosides
plastidial mRNA stabilisation
repair and reassembly cycle
A–class RAB GTPase
ERF subfamily
cytoplasm–localized receptor complex
subgroup ERF–IX transcription factor
receptor component *(PYL/RCAR)
subgroup ERF–VIII transcription factor
P4–type ATPase component *(ALA)
G1
G3
G5
-log10(adj p)
6
4
2
0
Gene Count
10
20
40
Mercator level
level1
level2
level3
level4
level5
level6
level7

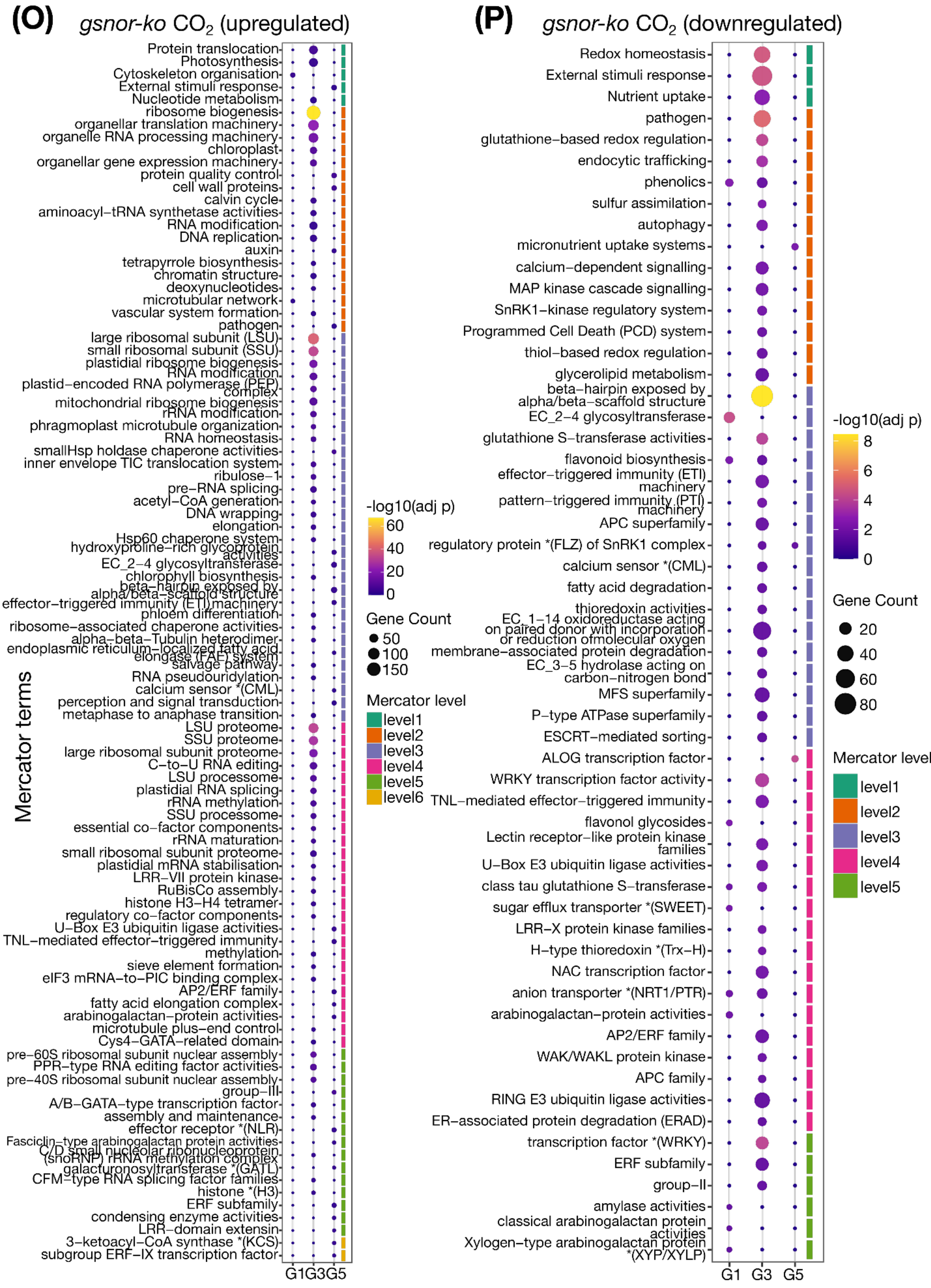
(O) gsnor-ko CO2 (upregulated)
Mercator terms
Protein translocation
Photosynthesis
Cytoskeleton organisation
External stimuli response
Nucleotide metabolism
ribosome biogenesis
organellar translation machinery
organelle RNA processing machinery
chloroplast
organellar gene expression machinery
protein quality control
cell wall proteins
calvin cycle
aminoacyl−tRNA synthetase activities
RNA modification
DNA replication
auxin
tetrapyrrole biosynthesis
chromatin structure
deoxynucleotides
microtubular network
vascular system formation
pathogen
large ribosomal subunit (LSU)
small ribosomal subunit (SSU)
plastidial ribosome biogenesis
RNA modification
plastid−encoded RNA polymerase (PEP) complex
mitochondrial ribosome biogenesis
rRNA modification
phragmoplast microtubule organization
RNA homeostasis
smallHsp holdase chaperone activities
inner envelope TIC translocation system
ribulose−1
pre−RNA splicing
acetyl−CoA generation
DNA wrapping
elongation
Hsp60 chaperone system
hydroxyproline−rich glycoprotein activities
EC_2−4 glycosyltransferase
chlorophyll biosynthesis
beta−hairpin exposed by alpha/beta−scaffold structure
effector−triggered immunity (ETI) machinery
phloem differentiation
ribosome−associated chaperone activities
alpha−beta−Tubulin heterodimer
endoplasmic reticulum−localized fatty acid elongase (FAE) system
salvage pathway
RNA pseudouridylation
calcium sensor *(CML)
perception and signal transduction
metaphase to anaphase transition
LSU proteome
SSU proteome
large ribosomal subunit proteome
C−to−U RNA editing
LSU processome
plastidial RNA splicing
rRNA methylation
SSU processome
essential co−factor components
rRNA maturation
small ribosomal subunit proteome
plastidial mRNA stabilisation
LRR−VII protein kinase
RuBisCo assembly
histone H3−H4 tetramer
regulatory co−factor components
U−Box E3 ubiquitin ligase activities
TNL−mediated effector−triggered immunity
methylation
sieve element formation
eIF3 mRNA−to−PIC binding complex
AP2/ERF family
fatty acid elongation complex
arabinogalactan−protein activities
microtubule plus−end control
Cys4−GATA−related domain
pre−60S ribosomal subunit nuclear assembly
PPR−type RNA editing factor activities
pre−40S ribosomal subunit nuclear assembly
group−III
A/B−GATA−type transcription factor
assembly and maintenance
effector receptor *(NLR)
Fasciclin−type arabinogalactan protein activities
C/D small nucleolar ribonucleoprotein (snoRNP) rRNA methylation complex
galacturonosyltransferase *(GATL)
CFM−type RNA splicing factor families
histone *(H3)
ERF subfamily
condensing enzyme activities
LRR−domain extensin
3−ketoacyl−CoA synthase *(KCS)
subgroup ERF−IX transcription factor
G1 G3 G5
-log10(adj p)
60
40
20
0
Gene Count
50
100
150
Mercator level
level1
level2
level3
level4
level5
level6
(P) gsnor-ko CO2 (downregulated)
Redox homeostasis
External stimuli response
Nutrient uptake
pathogen
glutathione−based redox regulation
endocytic trafficking
phenolics
sulfur assimilation
autophagy
micronutrient uptake systems
calcium−dependent signalling
MAP kinase cascade signalling
SnRK1−kinase regulatory system
Programmed Cell Death (PCD) system
thiol−based redox regulation
glycerolipid metabolism
beta−hairpin exposed by alpha/beta−scaffold structure
EC_2−4 glycosyltransferase
glutathione S−transferase activities
flavonoid biosynthesis
effector−triggered immunity (ETI) machinery
pattern−triggered immunity (PTI) machinery
APC superfamily
regulatory protein *(FLZ) of SnRK1 complex
calcium sensor *(CML)
fatty acid degradation
thioredoxin activities
EC_1−14 oxidoreductase acting on paired donor with incorporation or reduction ofmolecular oxygen
membrane−associated protein degradation
EC_3−5 hydrolase acting on carbon−nitrogen bond
MFS superfamily
P−type ATPase superfamily
ESCRT−mediated sorting
ALOG transcription factor
WRKY transcription factor activity
TNL−mediated effector−triggered immunity
flavonol glycosides
Lectin receptor−like protein kinase families
U−Box E3 ubiquitin ligase activities
class tau glutathione S−transferase
sugar efflux transporter *(SWEET)
LRR−X protein kinase families
H−type thioredoxin *(Trx−H)
NAC transcription factor
anion transporter *(NRT1/PTR)
arabinogalactan−protein activities
AP2/ERF family
WAK/WAKL protein kinase
APC family
RING E3 ubiquitin ligase activities
ER−associated protein degradation (ERAD)
transcription factor *(WRKY)
ERF subfamily
group−II
amylase activities
classical arabinogalactan protein activities
Xylogen−type arabinogalactan protein *(XYP/XYLP)
G1 G3 G5
-log10(adj p)
8
6
4
2
0
Gene Count
20
40
60
80
Mercator level
level1
level2
level3
level4
level5

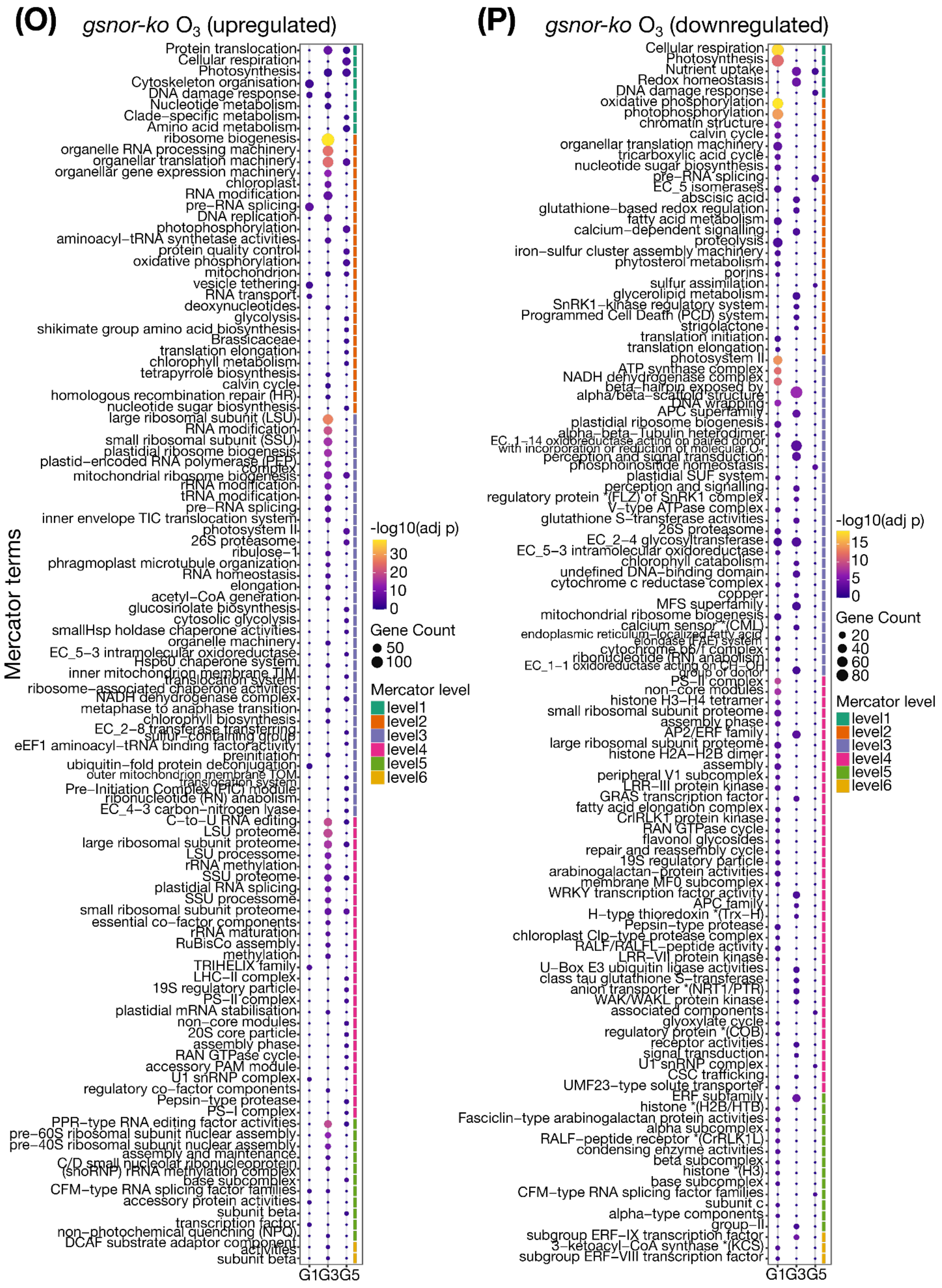

(O)
gsnor-ko O3 (upregulated)
Mercator terms
Protein translocation
Cellular respiration
Photosynthesis
Cytoskeleton organisation
DNA damage response
Nucleotide metabolism
Clade-specific metabolism
Amino acid metabolism
ribosome biogenesis
organelle RNA processing machinery
organellar translation machinery
organellar gene expression machinery
chloroplast
RNA modification
pre-RNA splicing
DNA replication
photophosphorylation
aminoacyl-tRNA synthetase activities
protein quality control
oxidative phosphorylation
mitochondrion
vesicle tethering
RNA transport
deoxynucleotides
glycolysis
shikimate group amino acid biosynthesis
Brassicaceae
translation elongation
chlorophyll metabolism
tetrapyrrole biosynthesis
calvin cycle
homologous recombination repair (HR)
nucleotide sugar biosynthesis
large ribosomal subunit (LSU)
RNA modification
small ribosomal subunit (SSU)
plastidial ribosome biogenesis
plastid-encoded RNA polymerase (PEP) complex
mitochondrial ribosome biogenesis
rRNA modification
tRNA modification
pre-RNA splicing
inner envelope TIC translocation system
photosystem II
26S proteasome
ribulose-1
phragmoplast microtubule organization
RNA homeostasis
elongation
acetyl-CoA generation
glucosinolate biosynthesis
cytosolic glycolysis
smallHsp holdase chaperone activities
organelle machinery
EC_5-3 intramolecular oxidoreductase
Hsp60 chaperone system
inner mitochondrion membrane TIM translocation system
ribosome-associated chaperone activities
NADH dehydrogenase complex
metaphase to anaphase transition
chlorophyll biosynthesis
EC_2-8 transferase transferring sulfur-containing group
eEF1 aminoacyl-tRNA binding factor activity
preinitiation
ubiquitin-fold protein deconjugation
outer mitochondrion membrane TOM translocation system
Pre-Initiation Complex (PIC) module
ribonucleotide (RN) anabolism
EC_4-3 carbon-nitrogen lyase
C-to-U RNA editing
LSU proteome
large ribosomal subunit proteome
LSU processome
rRNA methylation
SSU proteome
plastidial RNA splicing
SSU processome
small ribosomal subunit proteome
essential co-factor components
rRNA maturation
RuBisCo assembly
methylation
TRIHELIX family
LHC-II complex
19S regulatory particle
PS-II complex
plastidial mRNA stabilisation
non-core modules
20S core particle
assembly phase
RAN GTPase cycle
accessory PAM module
U1 snRNP complex
regulatory co-factor components
Pepsin-type protease
PS-I complex
PPR-type RNA editing factor activities
pre-60S ribosomal subunit nuclear assembly
pre-40S ribosomal subunit nuclear assembly
assembly and maintenance
C/D small nucleolar ribonucleoprotein (snoRNP) rRNA methylation complex
base subcomplex
CFM-type RNA splicing factor families
accessory protein activities
subunit beta
transcription factor
non-photochemical quenching (NPQ)
DCAF substrate adaptor component activities
subunit beta
G1 G3 G5
-log10(adj p)
30
20
10
0
Gene Count
50
100
Mercator level
level1
level2
level3
level4
level5
level6
(P)
gsnor-ko O3 (downregulated)
Cellular respiration
Photosynthesis
Nutrient uptake
Redox homeostasis
DNA damage response
oxidative phosphorylation
photophosphorylation
chromatin structure
calvin cycle
organellar translation machinery
tricarboxylic acid cycle
nucleotide sugar biosynthesis
pre-RNA splicing
EC_5 isomerases
abscisic acid
glutathione-based redox regulation
fatty acid metabolism
calcium-dependent signalling
proteolysis
iron-sulfur cluster assembly machinery
phytosterol metabolism
porins
sulfur assimilation
glycerolipid metabolism
SnRK1-kinase regulatory system
Programmed Cell Death (PCD) system
strigolactone
translation initiation
translation elongation
photosystem II
ATP synthase complex
NADH dehydrogenase complex
beta-hairpin exposed by alpha/beta-scaffold structure
DNA wrapping
APC superfamily
plastidial ribosome biogenesis
alpha-beta-Tubulin heterodimer
EC_1-14 oxidoreductase acting on paired donor with incorporation or reduction of molecular O2
perception and signal transduction
phosphoinositide homeostasis
plastidial SUF system
perception and signalling
regulatory protein *(FLZ) of SnRK1 complex
V-type ATPase complex
glutathione S-transferase activities
26S proteasome
EC_2-4 glycosyltransferase
EC_5-3 intramolecular oxidoreductase
chlorophyll catabolism
undefined DNA-binding domain
cytochrome c reductase complex
copper
MFS superfamily
mitochondrial ribosome biogenesis
calcium sensor *(CML)
endoplasmic reticulum-localized fatty acid elongase (FAE) system
cytochrome b6/f complex
ribonucleotide (RN) anabolism
EC_1-1 oxidoreductase acting on CH-OH group of donor
PS-II complex
non-core modules
histone H3-H4 tetramer
small ribosomal subunit proteome
assembly phase
AP2/ERF family
large ribosomal subunit proteome
histone H2A-H2B dimer
assembly
peripheral V1 subcomplex
LRR-III protein kinase
GRAS transcription factor
fatty acid elongation complex
CrRLK1 protein kinase
RAN GTPase cycle
flavonol glycosides
repair and reassembly cycle
19S regulatory particle
arabinogalactan-protein activities
membrane MF0 subcomplex
WRKY transcription factor activity
APC family
H-type thioredoxin *(Trx-H)
Pepsin-type protease
chloroplast Clp-type protease complex
RALF/RALFL-peptide activity
LRR-VII protein kinase
U-Box E3 ubiquitin ligase activities
class tau glutathione S-transferase
anion transporter *(NRT1/PTR)
WAK/WAKL protein kinase
associated components
glyoxylate cycle
regulatory protein *(COB)
receptor activities
signal transduction
U1 snRNP complex
CSC trafficking
UMF23-type solute transporter
ERF subfamily
histone *(H2B/HTB)
Fasciclin-type arabinogalactan protein activities
alpha subcomplex
RALF-peptide receptor *(CrRLK1L)
condensing enzyme activities
beta subcomplex
histone *(H3)
base subcomplex
CFM-type RNA splicing factor families
subunit c
alpha-type components
group-II
subgroup ERF-IX transcription factor
3-ketoacyl-CoA synthase *(KCS)
subgroup ERF-VIII transcription factor
G1 G3 G5
-log10(adj p)
15
10
5
0
Gene Count
20
40
60
80
Mercator level
level1
level2
level3
level4
level5
level6

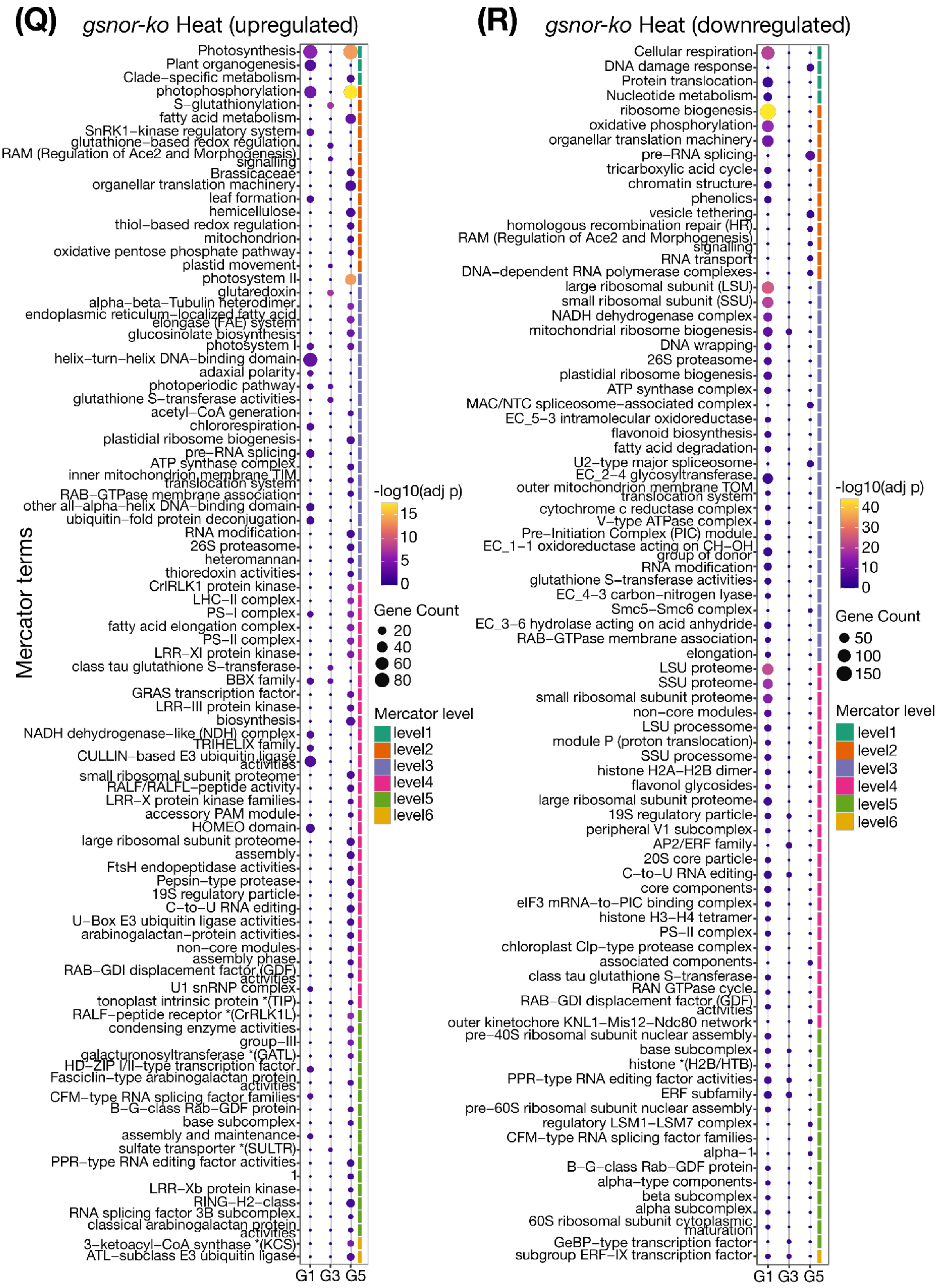
(Q) gsnor-ko Heat (upregulated)
Mercator terms
Photosynthesis
Plant organogenesis
Clade−specific metabolism
photophosphorylation
S−glutathionylation
fatty acid metabolism
SnRK1−kinase regulatory system
glutathione−based redox regulation
RAM (Regulation of Ace2 and Morphogenesis) signalling
Brassicaceae
organellar translation machinery
leaf formation
hemicellulose
thiol−based redox regulation
mitochondrion
oxidative pentose phosphate pathway
plastid movement
photosystem II
glutaredoxin
alpha−beta−Tubulin heterodimer
endoplasmic reticulum−localized fatty acid elongase (FAE) system
glucosinolate biosynthesis
photosystem I
helix−turn−helix DNA−binding domain
adaxial polarity
photoperiodic pathway
glutathione S−transferase activities
acetyl−CoA generation
chlororespiration
plastidial ribosome biogenesis
pre−RNA splicing
ATP synthase complex
inner mitochondrion membrane TIM translocation system
RAB−GTPase membrane association
other all−alpha−helix DNA−binding domain
ubiquitin−fold protein deconjugation
RNA modification
26S proteasome
heteromannan
thioredoxin activities
CrRLK1 protein kinase
LHC−II complex
PS−I complex
fatty acid elongation complex
PS−II complex
LRR−XI protein kinase
class tau glutathione S−transferase
BBX family
GRAS transcription factor
LRR−III protein kinase
biosynthesis
NADH dehydrogenase−like (NDH) complex
TRIHELIX family
CULLIN−based E3 ubiquitin ligase activities
small ribosomal subunit proteome
RALF/RALFL−peptide activity
LRR−X protein kinase families
accessory PAM module
HOMEO domain
large ribosomal subunit proteome
assembly
FtsH endopeptidase activities
Pepsin−type protease
19S regulatory particle
C−to−U RNA editing
U−Box E3 ubiquitin ligase activities
arabinogalactan−protein activities
non−core modules
assembly phase
RAB−GDI displacement factor (GDF) activities
U1 snRNP complex
tonoplast intrinsic protein *(TIP)
RALF−peptide receptor *(CrRLK1L)
condensing enzyme activities
group−III
galacturonosyltransferase *(GATL)
HD−ZIP I/II−type transcription factor
Fasciclin−type arabinogalactan protein activities
CFM−type RNA splicing factor families
B−G−class Rab−GDF protein
base subcomplex
assembly and maintenance
sulfate transporter *(SULTR)
PPR−type RNA editing factor activities
1
LRR−Xb protein kinase
RING−H2−class
RNA splicing factor 3B subcomplex
classical arabinogalactan protein activities
3−ketoacyl−CoA synthase *(KCS)
ATL−subclass E3 ubiquitin ligase
G1 G3 G5
-log10(adj p)
15
10
5
0
Gene Count
20
40
60
80
Mercator level
level1
level2
level3
level4
level5
level6
(R) gsnor-ko Heat (downregulated)
Cellular respiration
DNA damage response
Protein translocation
Nucleotide metabolism
ribosome biogenesis
oxidative phosphorylation
organellar translation machinery
pre−RNA splicing
tricarboxylic acid cycle
chromatin structure
phenolics
vesicle tethering
homologous recombination repair (HR)
RAM (Regulation of Ace2 and Morphogenesis) signalling
RNA transport
DNA−dependent RNA polymerase complexes
large ribosomal subunit (LSU)
small ribosomal subunit (SSU)
NADH dehydrogenase complex
mitochondrial ribosome biogenesis
DNA wrapping
26S proteasome
plastidial ribosome biogenesis
ATP synthase complex
MAC/NTC spliceosome−associated complex
EC_5−3 intramolecular oxidoreductase
flavonoid biosynthesis
fatty acid degradation
U2−type major spliceosome
EC_2−4 glycosyltransferase
outer mitochondrion membrane TOM translocation system
cytochrome c reductase complex
V−type ATPase complex
Pre−Initiation Complex (PIC) module
EC_1−1 oxidoreductase acting on CH−OH group of donor
RNA modification
glutathione S−transferase activities
EC_4−3 carbon−nitrogen lyase
Smc5−Smc6 complex
EC_3−6 hydrolase acting on acid anhydride
RAB−GTPase membrane association
elongation
LSU proteome
SSU proteome
small ribosomal subunit proteome
non−core modules
LSU processome
module P (proton translocation)
SSU processome
histone H2A−H2B dimer
flavonol glycosides
large ribosomal subunit proteome
19S regulatory particle
peripheral V1 subcomplex
AP2/ERF family
20S core particle
C−to−U RNA editing
core components
eIF3 mRNA−to−PIC binding complex
histone H3−H4 tetramer
PS−II complex
chloroplast Clp−type protease complex
associated components
class tau glutathione S−transferase
RAN GTPase cycle
RAB−GDI displacement factor (GDF) activities
outer kinetochore KNL1−Mis12−Ndc80 network
pre−40S ribosomal subunit nuclear assembly
base subcomplex
histone *(H2B/HTB)
PPR−type RNA editing factor activities
ERF subfamily
pre−60S ribosomal subunit nuclear assembly
regulatory LSM1−LSM7 complex
CFM−type RNA splicing factor families
alpha−1
B−G−class Rab−GDF protein
alpha−type components
beta subcomplex
alpha subcomplex
60S ribosomal subunit cytoplasmic maturation
GeBP−type transcription factor
subgroup ERF−IX transcription factor
G1 G3 G5
-log10(adj p)
40
30
20
10
0
Gene Count
50
100
150
Mercator level
level1
level2
level3
level4
level5
level6

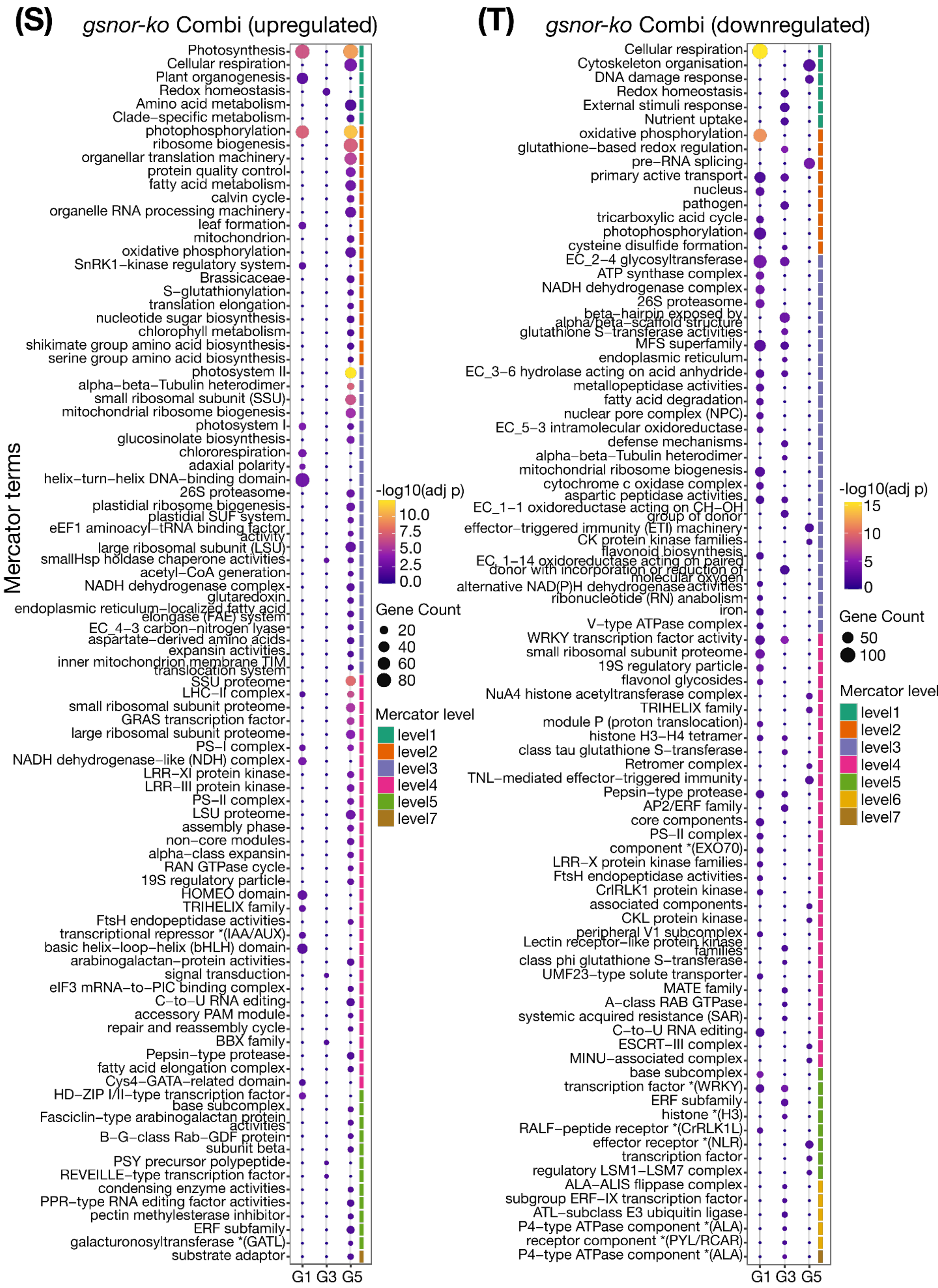


**Supplementary Fig S17. Mercator-based functional enrichment analysis across treatments, generations, and genotypes. (A-T)** Bubble plots showing significantly enriched Mercator categories for differentially expressed genes (DEGs) identified in WT and *gsnor-ko* plants under Drought, elevated $CO_2$, elevated $O_3$, Heat, and Combination treatments across generations G1, G3, and G5. Separate panels represent individual genotype × treatment × DEG-direction subsets. The x-axis indicates generation, and the y-axis shows enriched Mercator terms. Bubble size corresponds to the number of genes assigned to each term, and bubble colour indicates enrichment significance as $-\log_{10}$ adjusted *P*-value. The coloured bars at the right indicate Mercator hierarchical levels.

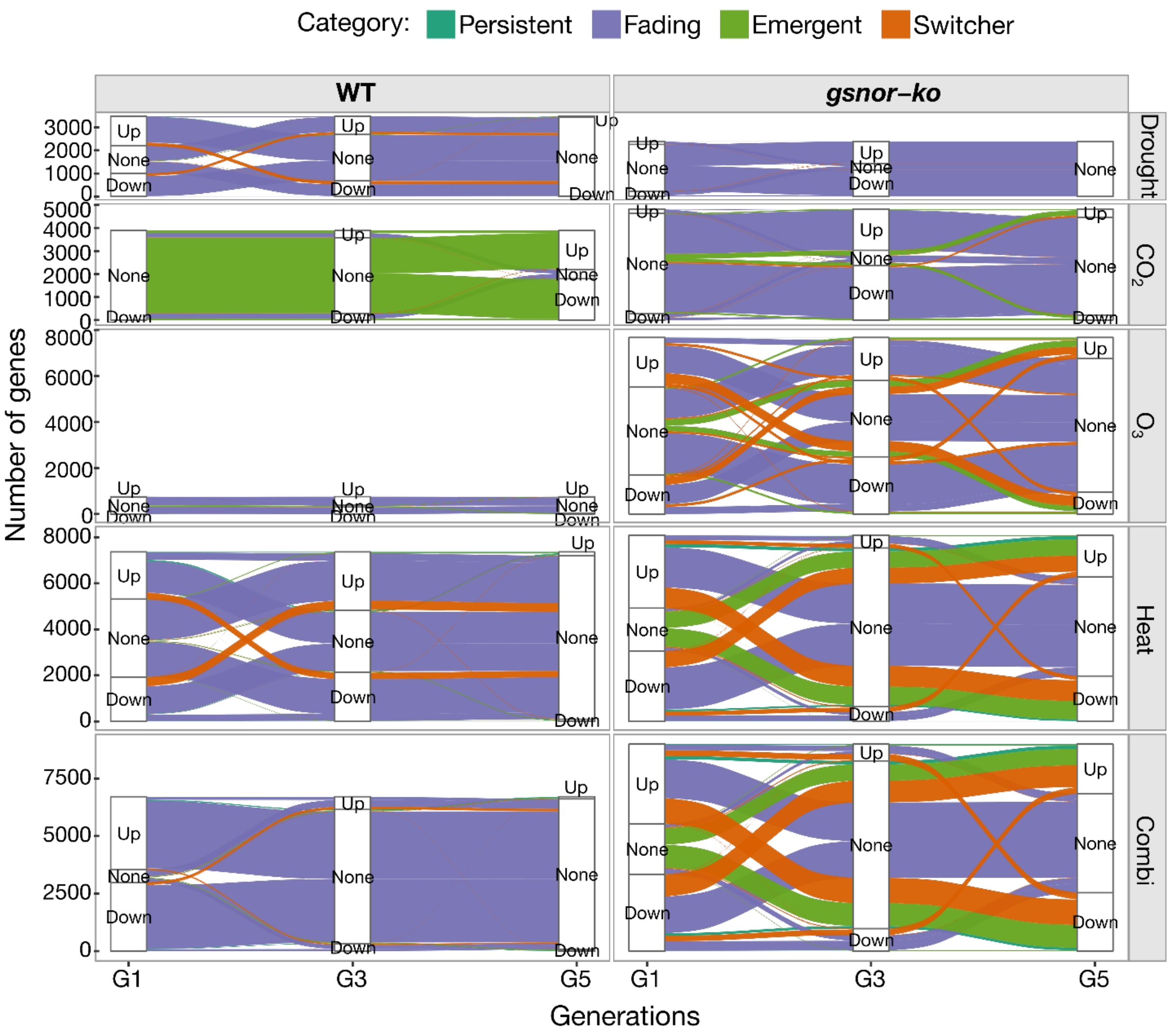


**Supplementary Fig S18. Alluvial plots showing multigenerational transitions in DEG states across treatments in WT and *gsnor-ko*.** Alluvial plots showing the distribution and transitions of differential expression states across generations G1, G3, and G5 for WT and *gsnor-ko* plants under Drought, elevated $CO_2$, $O_3$, Heat, and Combination treatments. For each treatment, genes were classified at each generation as Up, Down, or None (not significantly differentially expressed relative to the corresponding control). The height of each stratum represents the number of genes in a given state, and the connecting flows indicate how genes changed or maintained their expression state across generations. Panels highlight treatment-specific differences in transcriptomic persistence, resetting, and state switching between WT and *gsnor-ko*.

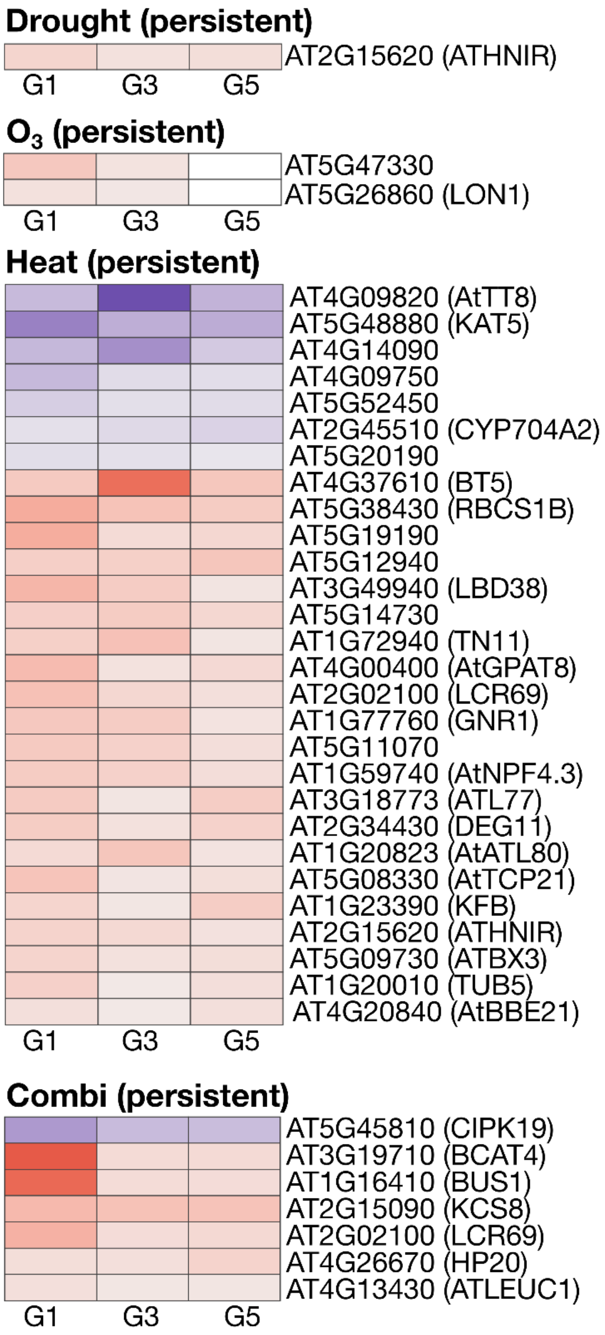


| Treatment | 3 generations | 2 generations |
|---|---|---|
| WT Drought | ✓ (plotted) | ✓ |
| WT $CO_2$ | ✗ | ✓ (plotted) |
| WT $O_3$ | ✗ | ✓ (plotted) |
| WT Heat | ✓ (plotted) | ✓ |
| WT Combi | ✓ (plotted) | ✓ |

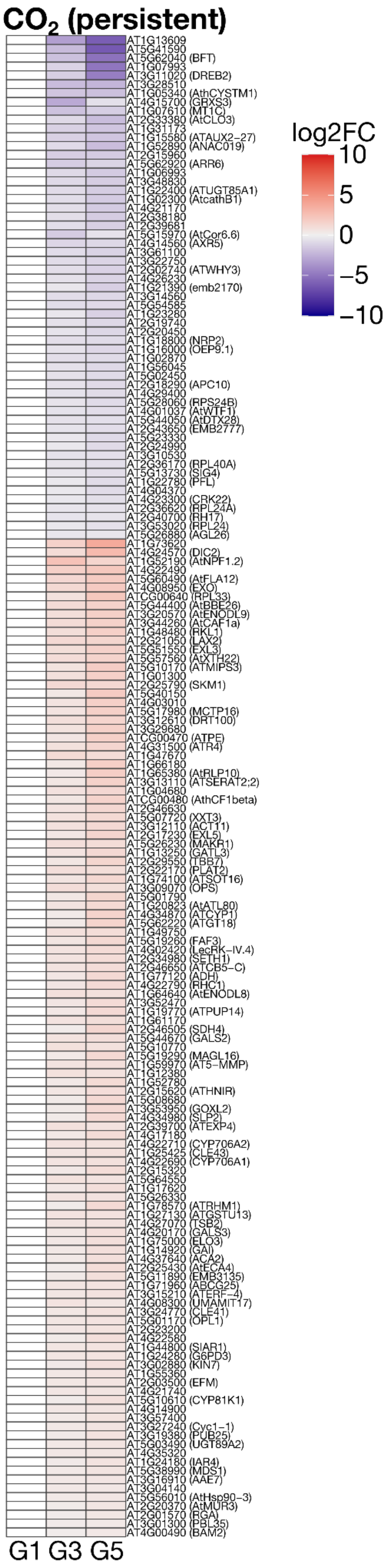


**Supplementary Fig S19. Persistent DEG signatures across generations in WT.** Heatmaps showing persistent differentially expressed genes (DEGs) in WT across generations under the indicated climate treatments. Persistent genes were defined as genes that remained differentially expressed in the same direction across either three generations (G1, G3, and G5) or two generations. Separate panels summarise treatment-specific persistent gene sets, and the table indicates which treatment categories yielded genes in each persistence class. The large heatmap summarises the combined persistent gene set across treatments. Columns represent generations and rows represent genes. Colour scale indicates relative expression change, with red indicating higher and blue indicating lower expression.

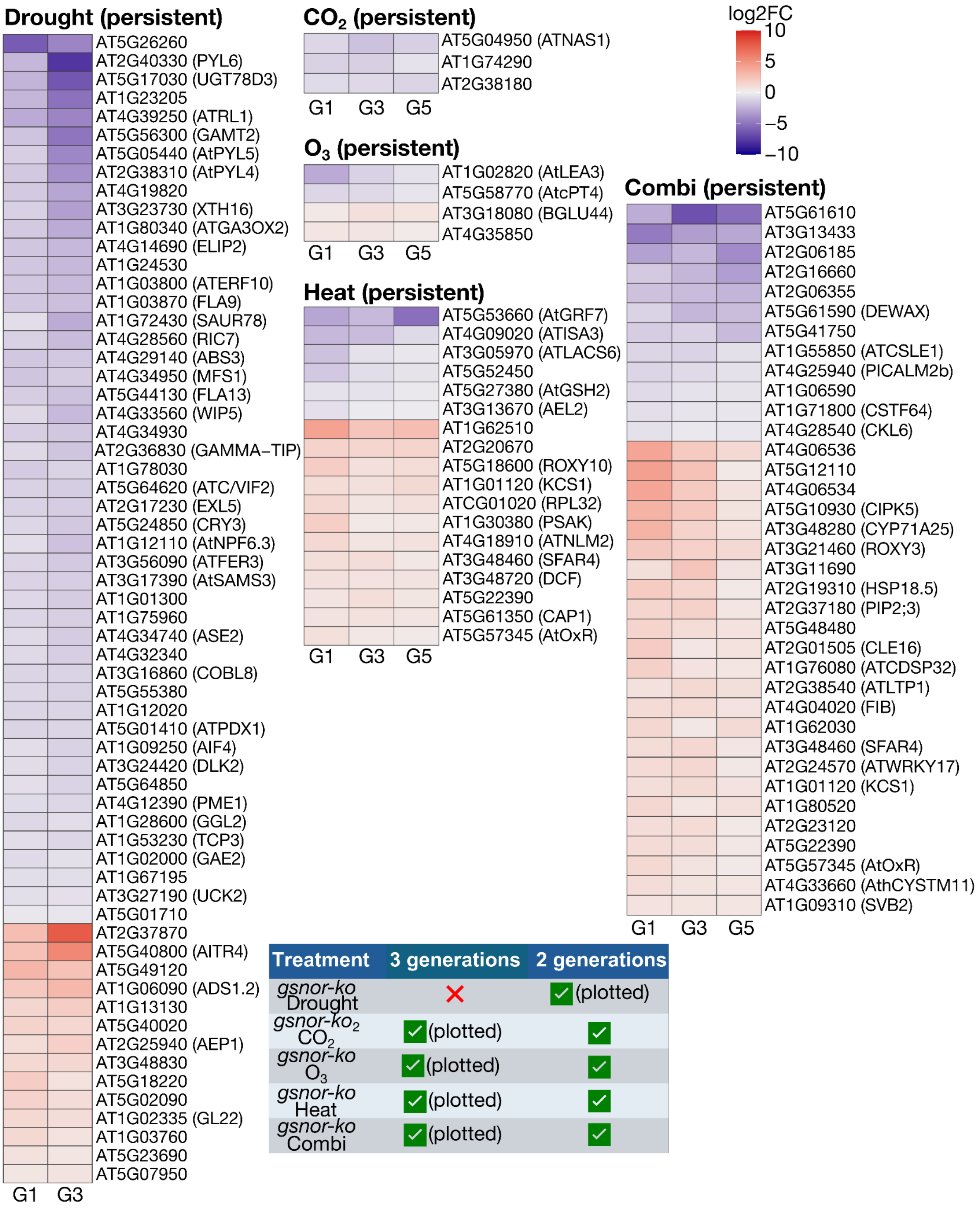

**Supplementary Fig S20. Persistent DEG signatures across generations in *gsnor-ko*.** Heatmaps showing persistent differentially expressed genes (DEGs) in *gsnor-ko* across generations under the indicated climate treatments. Persistent genes were defined as genes that remained differentially expressed in the same direction across either three generations (G1, G3, and G5) or two generations. Separate panels show treatment-specific persistent gene sets, and the summary table indicates which treatments yielded persistent genes. The large heatmap summarises the full set of persistent genes detected in *gsnor-ko*. Columns represent generations and rows represent genes. Colour scale indicates relative expression change, with red indicating higher and blue indicating lower expression.

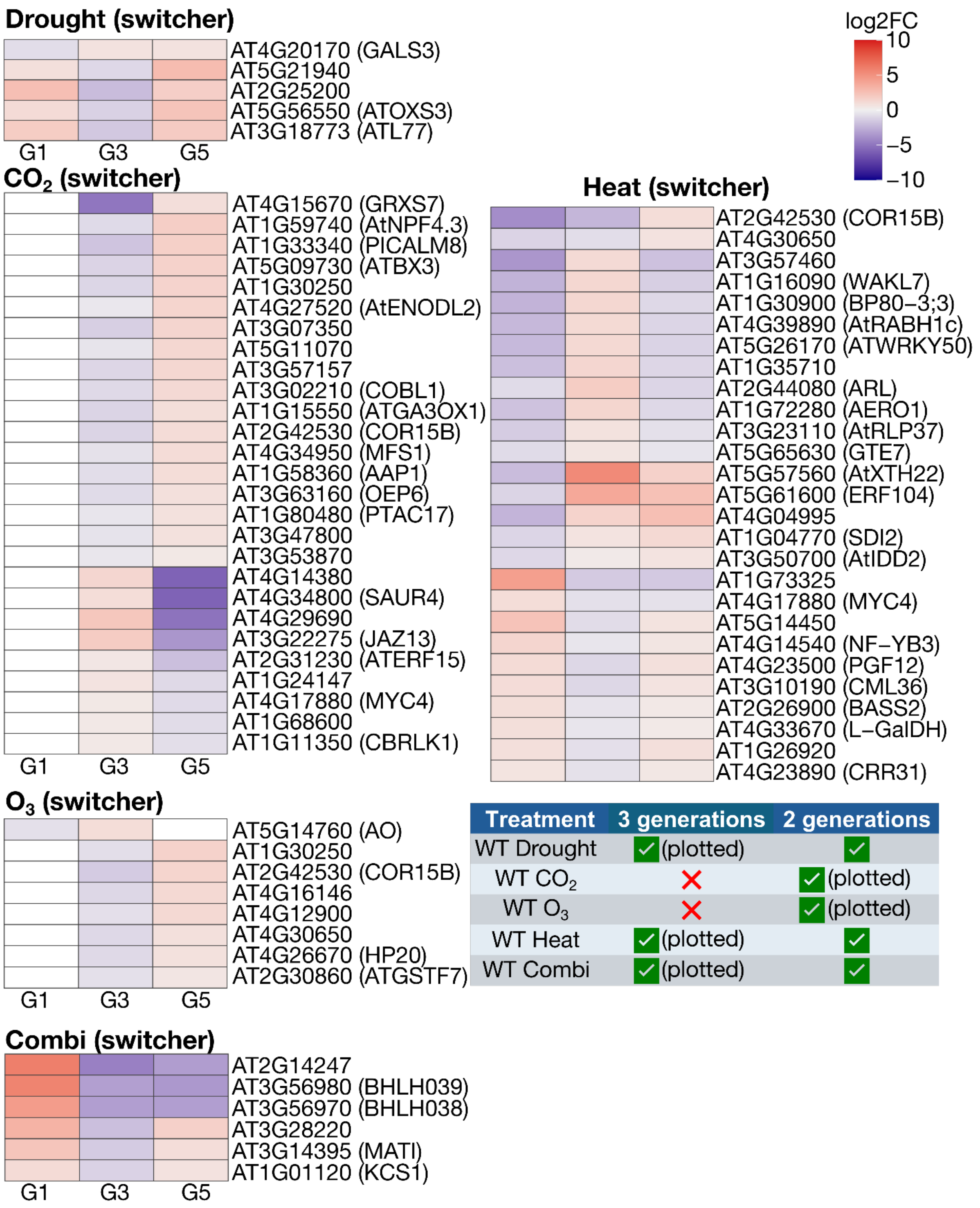


| Treatment | 3 generations | 2 generations |
|---|---|---|
| WT Drought | ✓ (plotted) | ✓ |
| WT $CO_2$ | ✗ | ✓ (plotted) |
| WT $O_3$ | ✗ | ✓ (plotted) |
| WT Heat | ✓ (plotted) | ✓ |
| WT Combi | ✓ (plotted) | ✓ |

**Supplementary Fig S21. Switcher genes across generations in WT.** Heatmaps showing switcher genes in WT, defined as genes that were differentially expressed across generations and changed the direction of regulation (e.g., from up-regulated to down-regulated, or vice versa). Separate panels show treatment-specific switcher-gene sets, and the accompanying table summarises whether switcher genes were identified across three generations or two generations for each treatment. The large heatmap summarises the combined switcher-gene set in WT. Columns represent generations and rows represent genes. Colour scale indicates relative expression change, with red indicating higher and blue indicating lower expression.

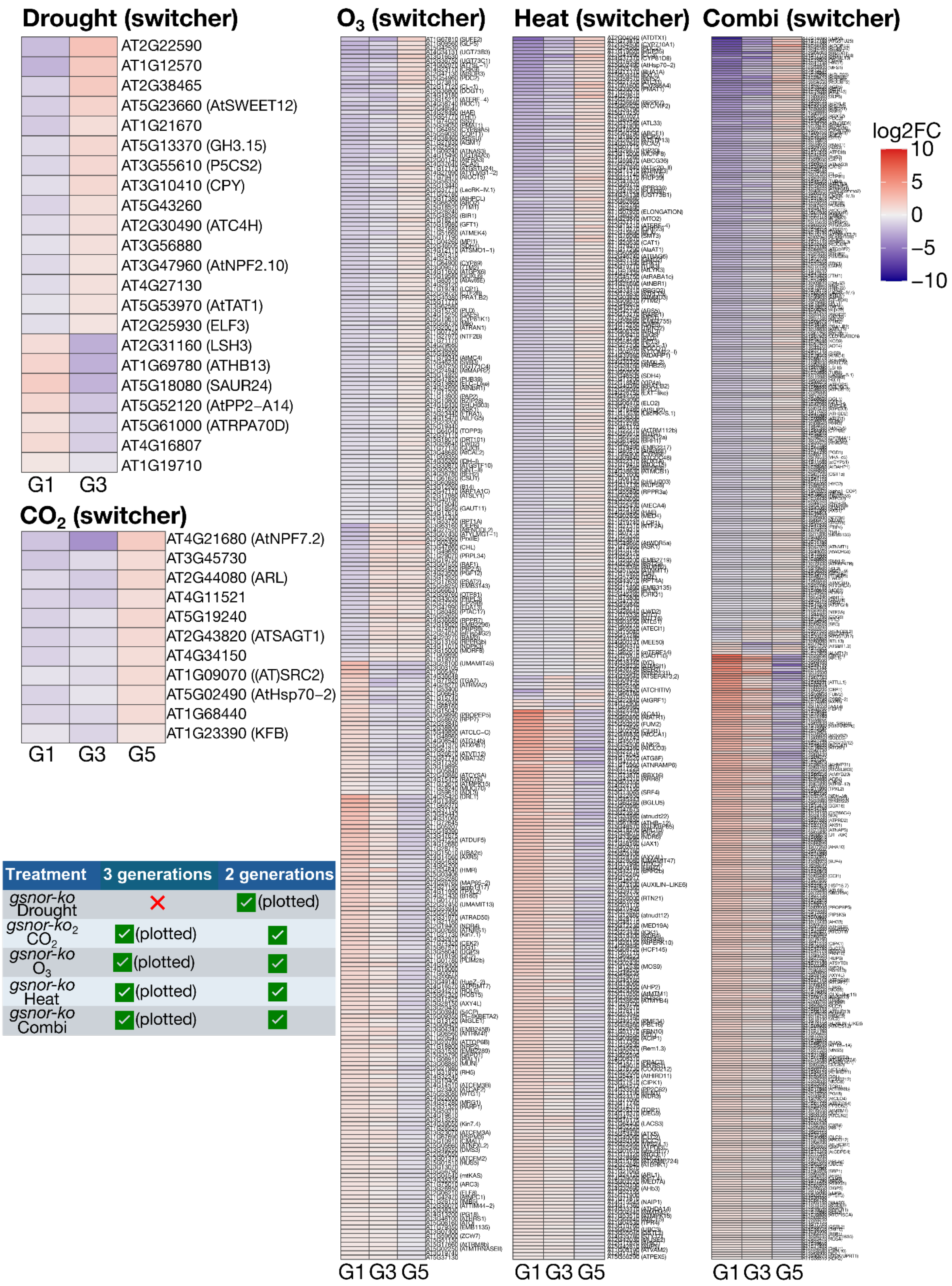


**Supplementary Fig S22. Switcher genes across generations in *gsnor-ko*.** Heatmaps showing switcher genes in *gsnor-ko*, defined as genes that were differentially expressed across generations and reversed the direction of regulation. Separate panels show treatment-specific switcher-gene sets, and the accompanying table summarises whether switcher genes were identified across three generations or two generations for each treatment. The large heatmap summarises the combined switcher-gene set in *gsnor-ko*. Columns represent generations and rows represent genes. Colour scale indicates relative expression change, with red indicating higher and blue indicating lower expression.

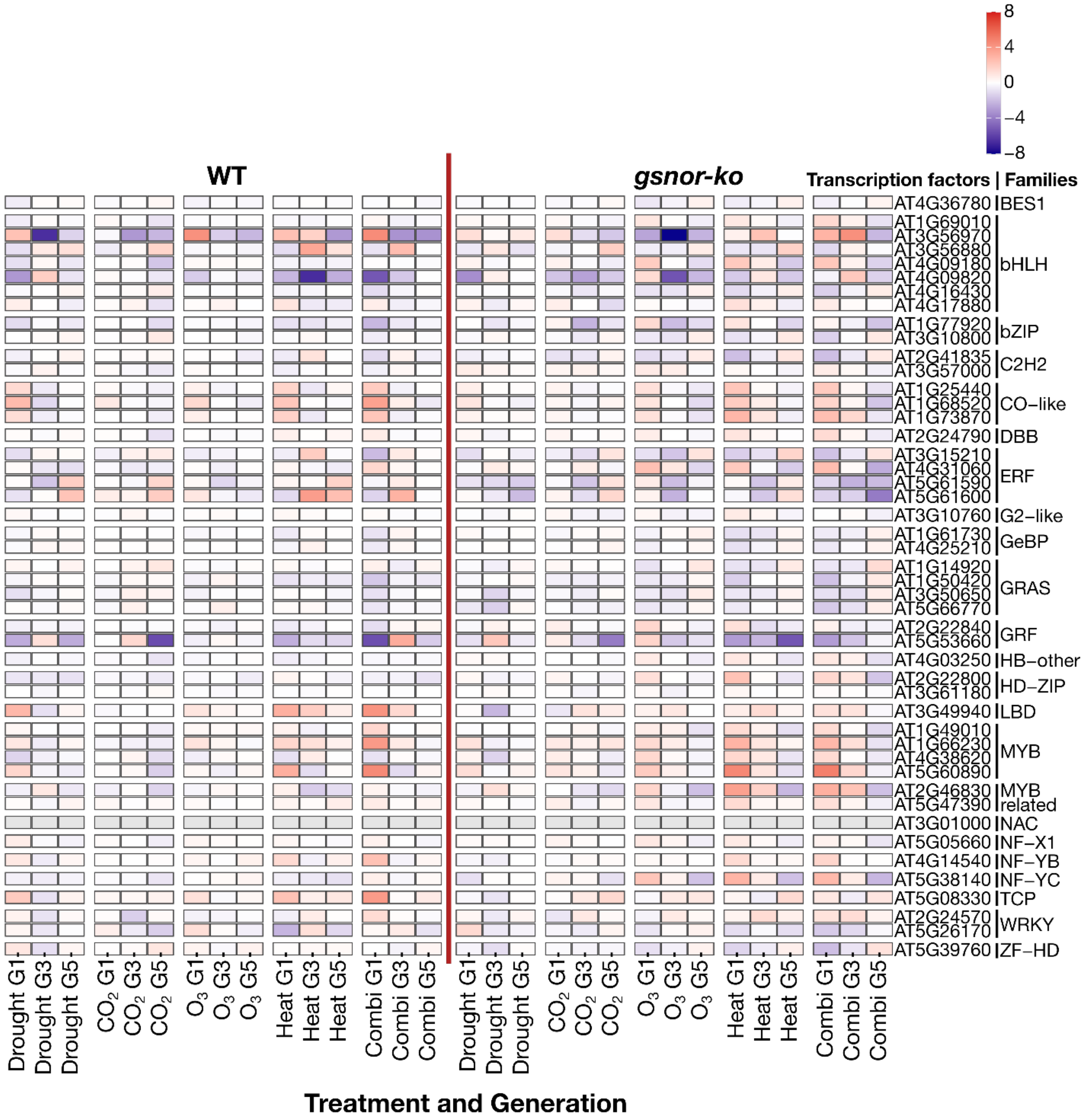


**Supplementary Fig S23. Common transcription factors shared across generations and treatments in WT and *gsnor-ko*.** Heatmap showing transcription factors (TFs) identified as common across multiple generations and treatment conditions in WT and *gsnor-ko* plants. Columns represent treatment × generation combinations, arranged by genotype, with WT shown on the left and *gsnor-ko* on the right; the red dashed line indicates the separation between the two genotypes. Rows represent shared TF genes. Colour scale indicates relative expression change, with red representing higher expression and blue representing lower expression. This overview highlights recurrent TF-associated regulatory signatures that were maintained across the multigenerational climate-stress design and reveals both shared and genotype-dependent patterns of transcriptional regulation.

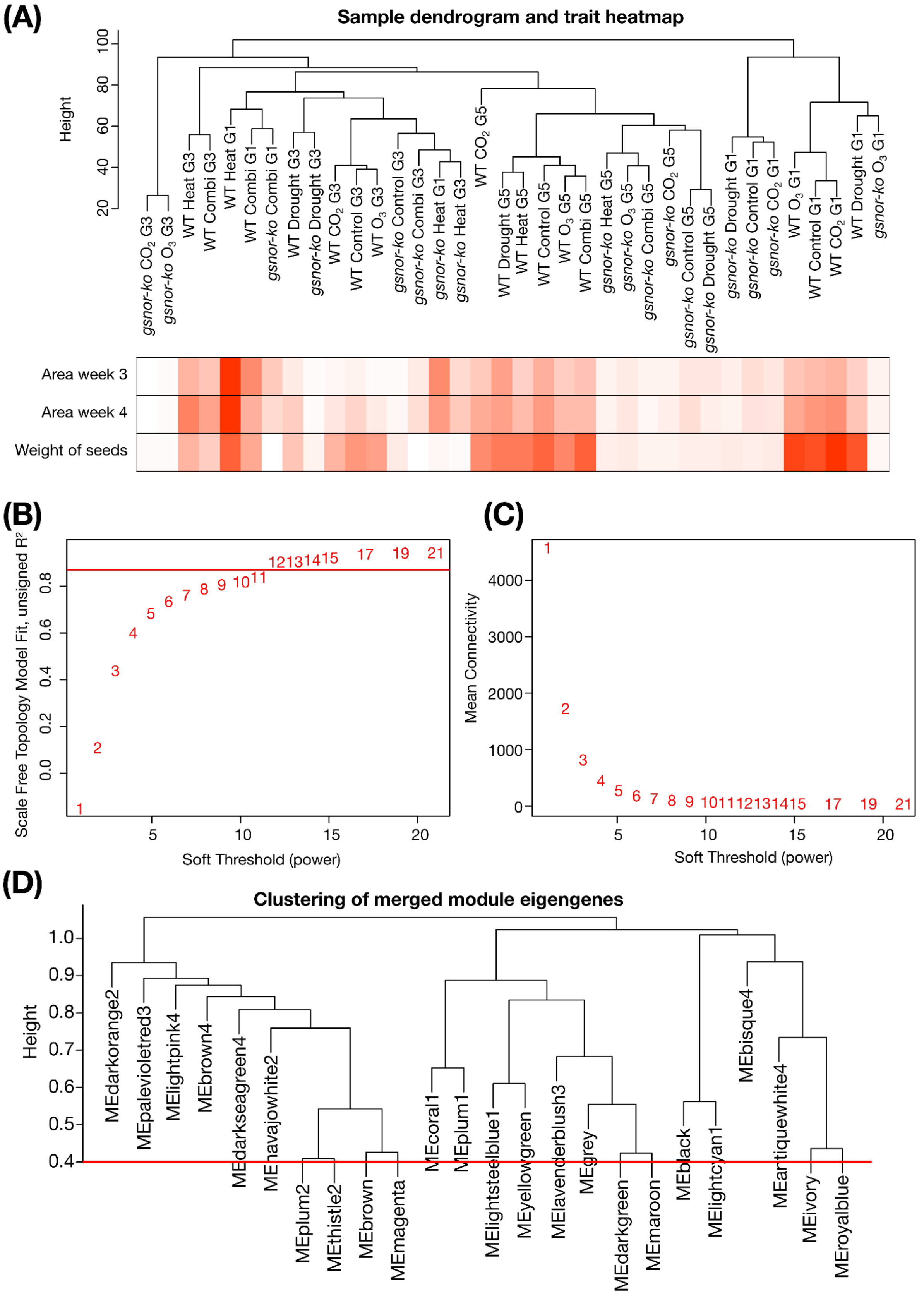


**Supplementary Fig S24. WGCNA network construction and module eigengene relationships. (A)** Dendrogram of all RNA-seq libraries included in the WGCNA, shown together with a trait heatmap summarising week 3 rosette area, week 4 rosette area, and seed weight across samples. **(B)** Scale-free topology fit index across candidate soft-thresholding powers. **(C)** Mean connectivity across the same range of powers. **(D)** Hierarchical clustering of module eigengenes; the red line indicates the threshold used for module merging.

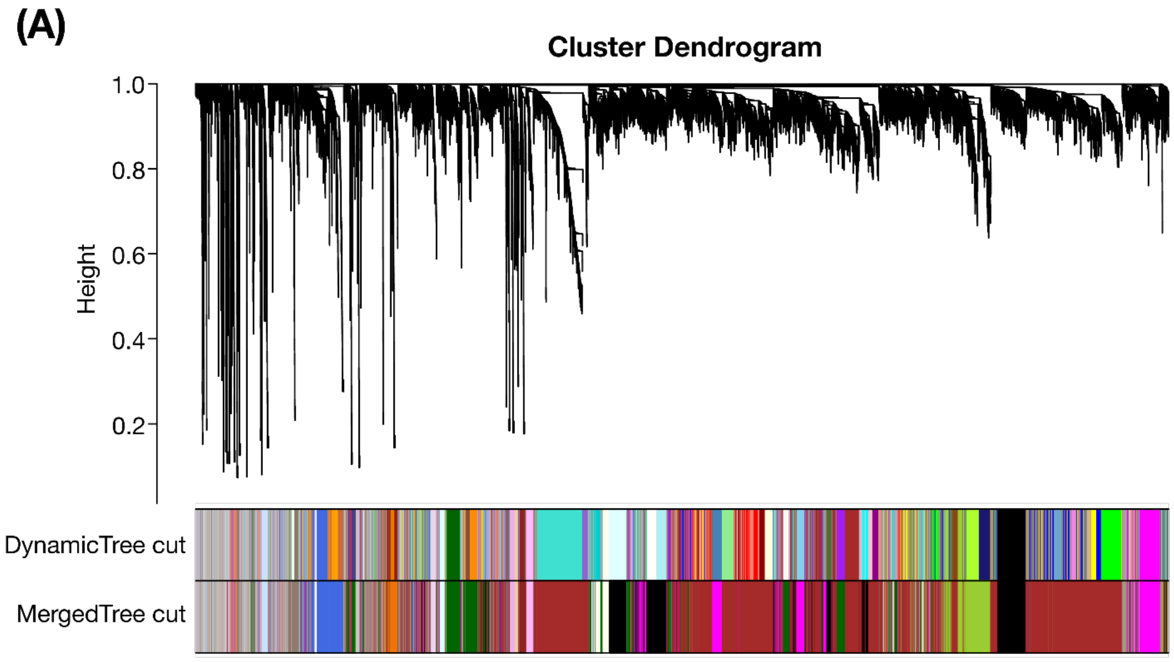


**(B)**

| Module | Putative role | Hub gene | Function |
|---|---|---|---|
| darkgreen | Genotype-associated module | AT2G44930 | Putative DUF247-containing transmembrane protein; currently uncharacterized |
| lightcyan1 | Genotype-associated module | AT1G59930 | MADS-box-family / imprinted gene; possible role in epigenetic or developmental regulation |
| magenta | Generation-trend module | AT5G58920 | Putative homeobox/prospero-like protein; functional evidence limited/unclear |
| ivory | Treatment-responsive module | AT5G25250 | Membrane microdomain protein involved in clathrin-independent endocytosis, seedling development, and stress-related membrane trafficking |

**Supplementary Fig S25. Gene clustering and representative WGCNA modules selected for downstream analysis. (A)** Hierarchical clustering of genes used for WGCNA, with module assignment shown before and after module merging (DynamicTree cut and Merged Tree cut). **(B)** Summary table of representative modules selected for downstream analysis, including their putative role, hub gene, and annotated function. Highlighted modules include darkgreen and lightcyan1 as genotype-associated modules, magenta as a generation-trend module, and ivory as a treatment-responsive module.

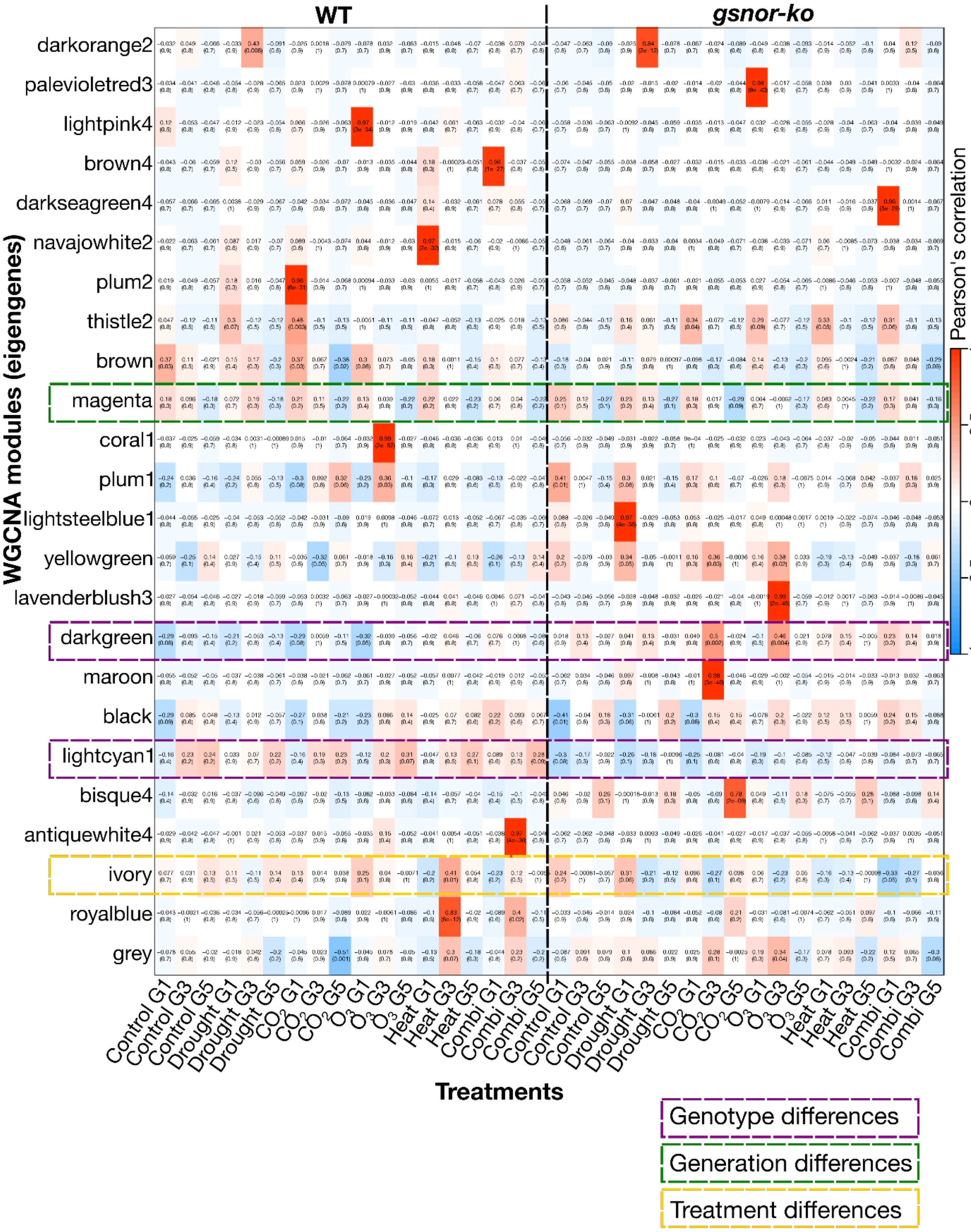


**Supplementary Fig S26. Module eigengene correlations across combined treatment groups.** Heatmap showing Pearson correlations between WGCNA module eigengenes and treatment × generation combinations in WT and *gsnor-ko*. Columns represent Control, Drought, $CO_2$, $O_3$, Heat, and Combination at G1, G3, and G5 for each genotype; rows represent individual co-expression modules. Values in cells indicate correlation coefficients, with *P*-values shown in parentheses. Dashed boxes highlight representative modules predominantly associated with genotype, generation, or treatment differences.

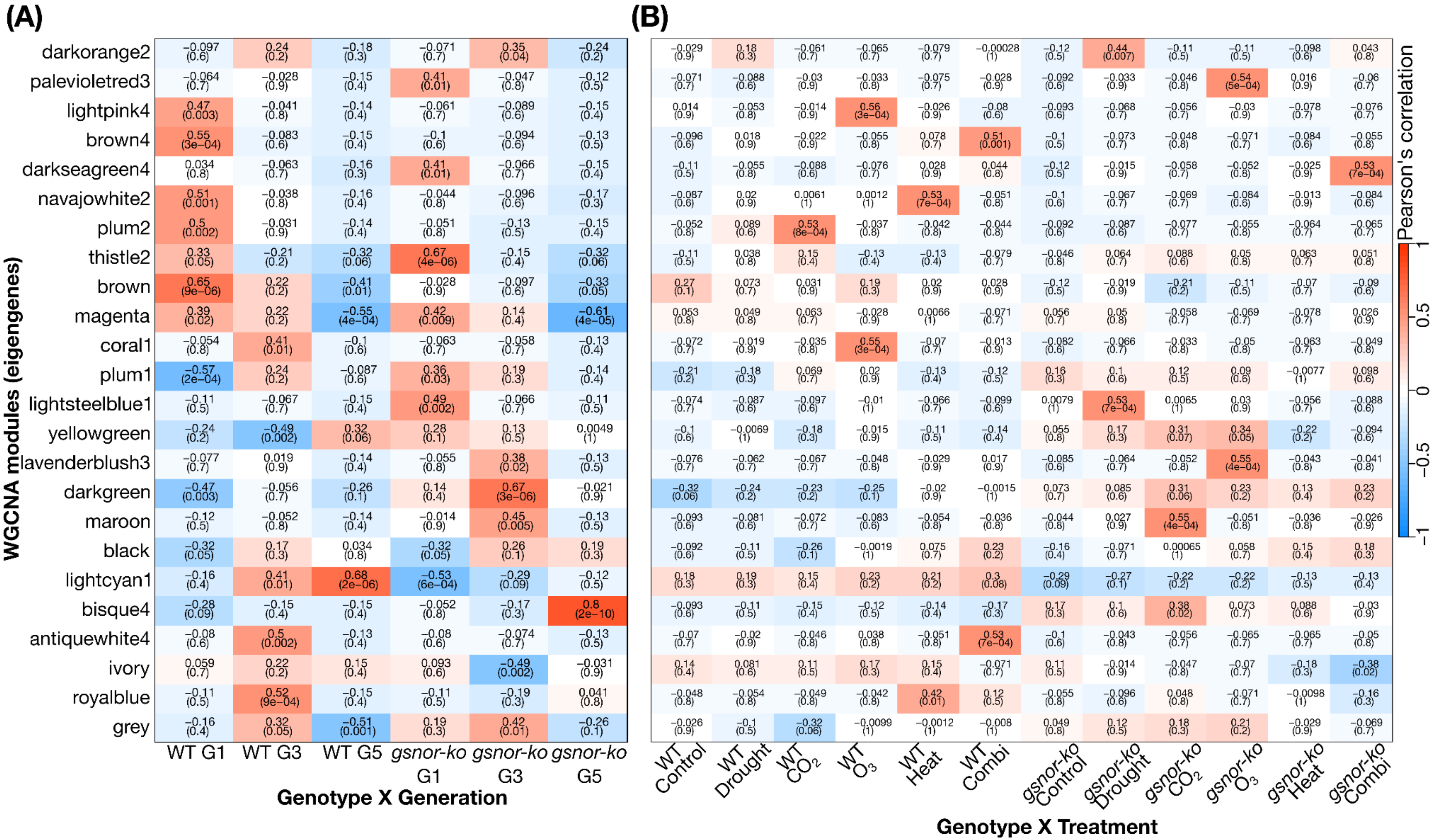
(A)
(B)
WGCNA modules (eigengenes)
darkorange2
palevioletred3
lightpink4
brown4
darkseagreen4
navajowhite2
plum2
thistle2
brown
magenta
coral1
plum1
lightsteelblue1
yellowgreen
lavenderblush3
darkgreen
maroon
black
lightcyan1
bisque4
antiquewhite4
ivory
royalblue
grey
WT G1
WT G3
WT G5
gsnor-ko G1
gsnor-ko G3
gsnor-ko G5
Genotype X Generation
WT Control
WT Drought
WT CO2
WT O3
WT Heat
WT Combi
gsnor-ko Control
gsnor-ko Drought
gsnor-ko CO2
gsnor-ko O3
gsnor-ko Heat
gsnor-ko Combi
Genotype X Treatment
Pearson's correlation
1
0.5
0
−0.5
−1

**Supplementary Fig S27. Genotype- and treatment-stratified module eigengene correlations. (A)** Heatmap of Pearson correlations between WGCNA module eigengenes and genotype × generation classes (WT G1, WT G3, WT G5, *gsnor-ko* G1, *gsnor-ko* G3, *gsnor-ko* G5). **(B)** Heatmap of Pearson correlations between module eigengenes and genotype × treatment classes in WT and *gsnor-ko*. Cell values show correlation coefficients, with *P*-values given in parentheses. These plots highlight modules whose behaviour is more strongly associated with genotype background and generation than with treatment alone.

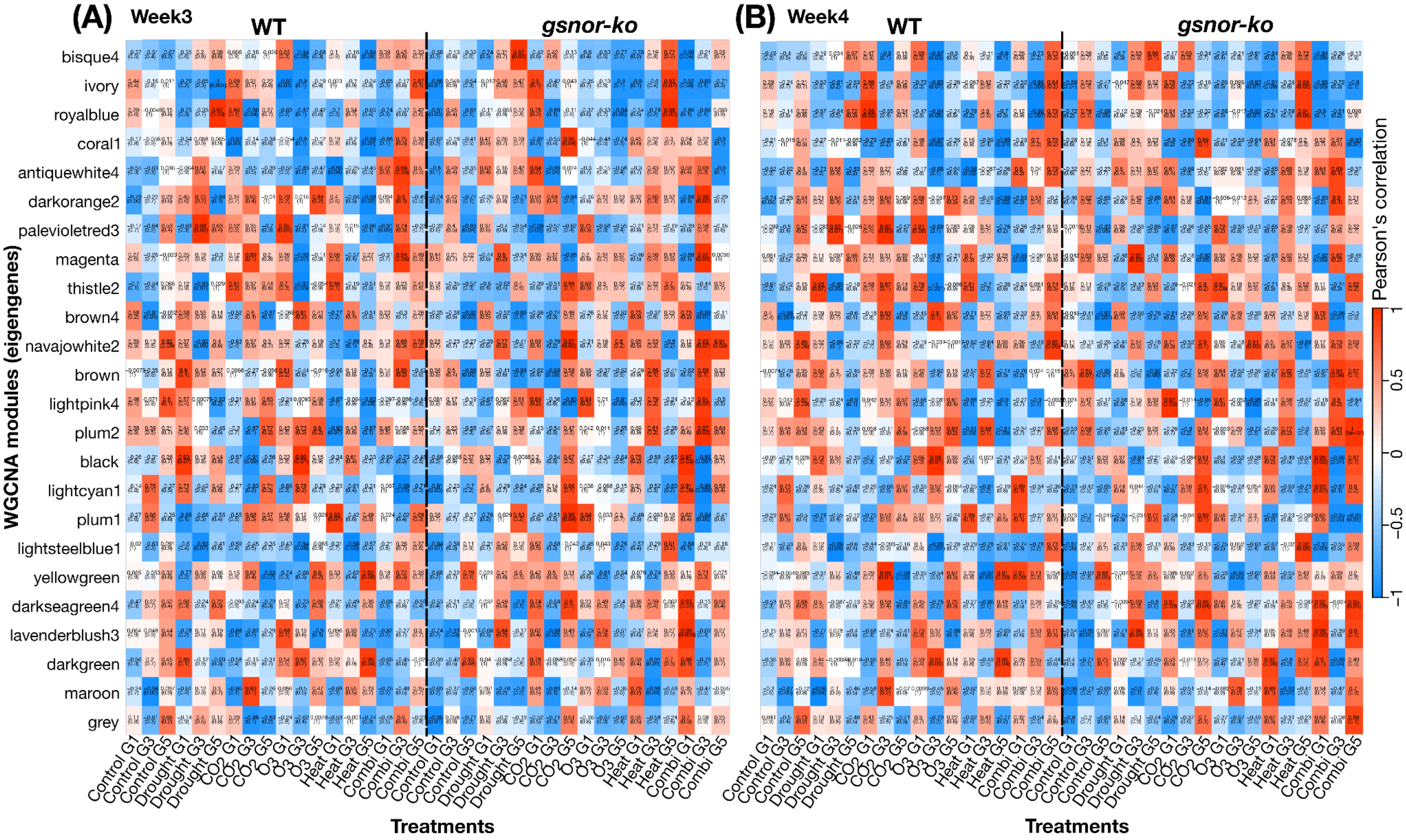

(A)
Week3
WT
gsnor-ko
(B)
Week4
WT
gsnor-ko
WGCNA modules (eigengenes)
bisque4
ivory
royalblue
coral1
antiquewhite4
darkorange2
palevioletred3
magenta
thistle2
brown4
navajowhite2
brown
lightpink4
plum2
black
lightcyan1
plum1
lightsteelblue1
yellowgreen
darkseagreen4
lavenderblush3
darkgreen
maroon
grey
Control G1
Control G3
Control G5
Drought G1
Drought G3
Drought G5
CO2 G1
CO2 G3
CO2 G5
O3 G1
O3 G3
O3 G5
Heat G1
Heat G3
Heat G5
Combi G1
Combi G3
Combi G5
Treatments
Pearson's correlation
1
0.5
0
−0.5
−1

**Supplementary Fig S28. Within-group correlations between WGCNA modules and phenotypic traits.** Heatmaps showing Pearson correlations between WGCNA module eigengenes and phenotypic measurements calculated within experimental groups. Rows represent co-expression modules and columns represent treatment × generation combinations in WT and *gsnor-ko*. Cell values indicate correlation coefficients, with *P*-values shown in parentheses. Upper and lower panels summarise separate within-group module–trait correlation analyses across the phenotypic dataset: **(A)** leaf area week 3; **(B)** leaf area week 4.

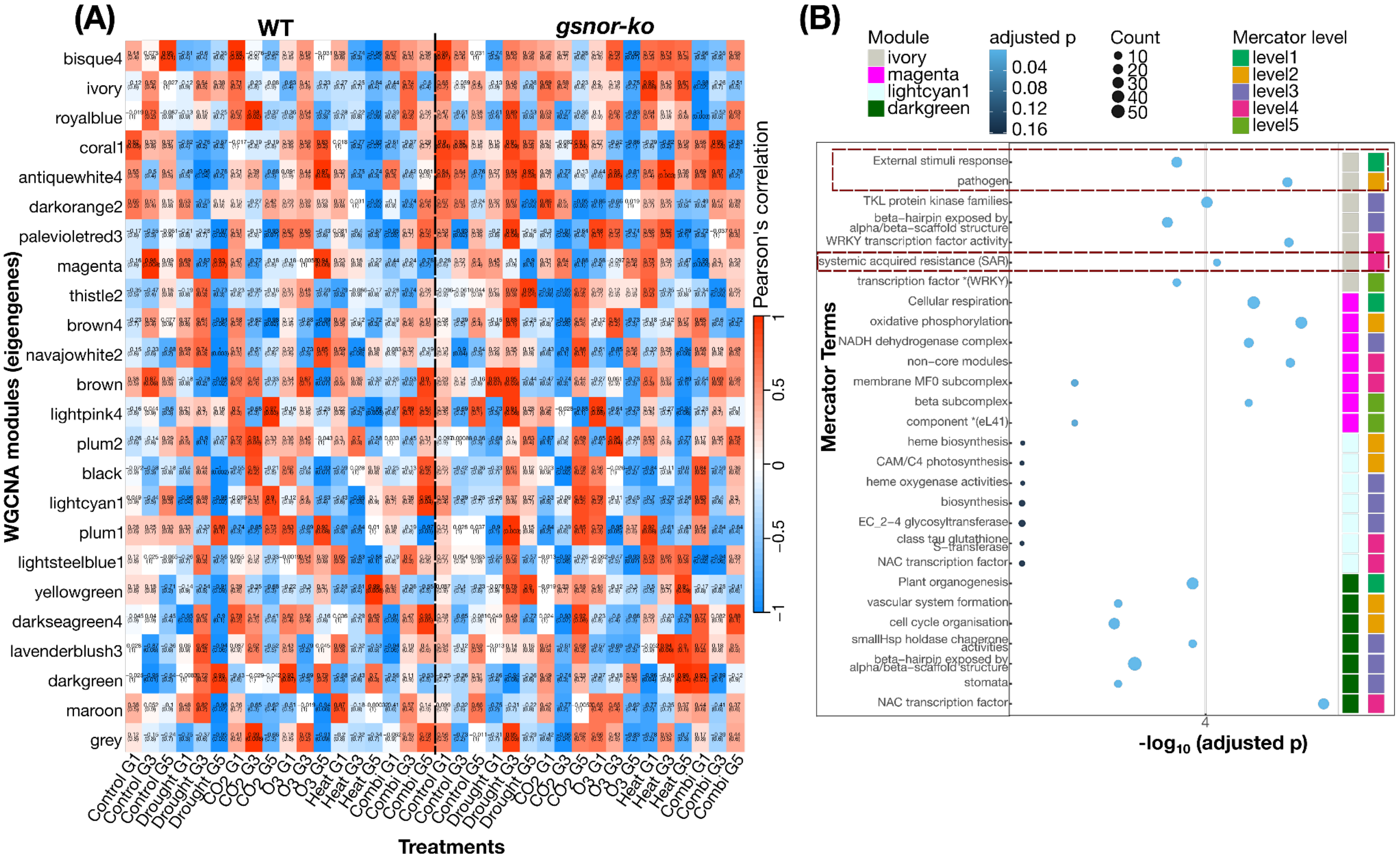

(A)
WT
gsnor-ko
WGCNA modules (eigengenes)
bisque4
ivory
royalblue
coral1
antiquewhite4
darkorange2
palevioletred3
magenta
thistle2
brown4
navajowhite2
brown
lightpink4
plum2
black
lightcyan1
plum1
lightsteelblue1
yellowgreen
darkseagreen4
lavenderblush3
darkgreen
maroon
grey
Pearson's correlation
1
0.5
0
−0.5
−1
Control G1
Control G3
Control G5
Drought G1
Drought G3
Drought G5
CO2 G1
CO2 G3
CO2 G5
O3 G1
O3 G3
O3 G5
Heat G1
Heat G3
Heat G5
Combi G1
Combi G3
Combi G5
Treatments
(B)
Module
ivory
magenta
lightcyan1
darkgreen
adjusted p
0.04
0.08
0.12
0.16
Count
10
20
30
40
50
Mercator level
level1
level2
level3
level4
level5
Mercator Terms
External stimuli response
pathogen
TKL protein kinase families
beta−hairpin exposed by alpha/beta−scaffold structure
WRKY transcription factor activity
systemic acquired resistance (SAR)
transcription factor *(WRKY)
Cellular respiration
oxidative phosphorylation
NADH dehydrogenase complex
non−core modules
membrane MF0 subcomplex
beta subcomplex
component *(eL41)
heme biosynthesis
CAM/C4 photosynthesis
heme oxygenase activities
biosynthesis
EC_2−4 glycosyltransferase
class tau glutathione S−transferase
NAC transcription factor
Plant organogenesis
vascular system formation
cell cycle organisation
smallHsp holdase chaperone activities
beta−hairpin exposed by alpha/beta−scaffold structure
stomata
NAC transcription factor
4
-log10 (adjusted p)

**Supplementary Fig S29. Module–trait relationships and Mercator enrichment of selected WGCNA modules. (A)** Heatmaps showing Pearson correlations between WGCNA module eigengenes and phenotypic measurements calculated within experimental groups. Rows represent co-expression modules and columns represent treatment × generation combinations in WT and *gsnor-ko*. Cell values indicate correlation coefficients, with *P*-values shown in parentheses. Upper and lower panels summarise separate within-group module–trait correlation analyses across the phenotypic dataset: weight of seeds **(B),** Mercator-based enrichment analysis of the selected modules darkgreen, lightcyan1, magenta, and ivory. Bubble size indicates the number of genes assigned to each term, and colour indicates adjusted *P*-value. Side annotation bars indicate module identity and Mercator functional level.

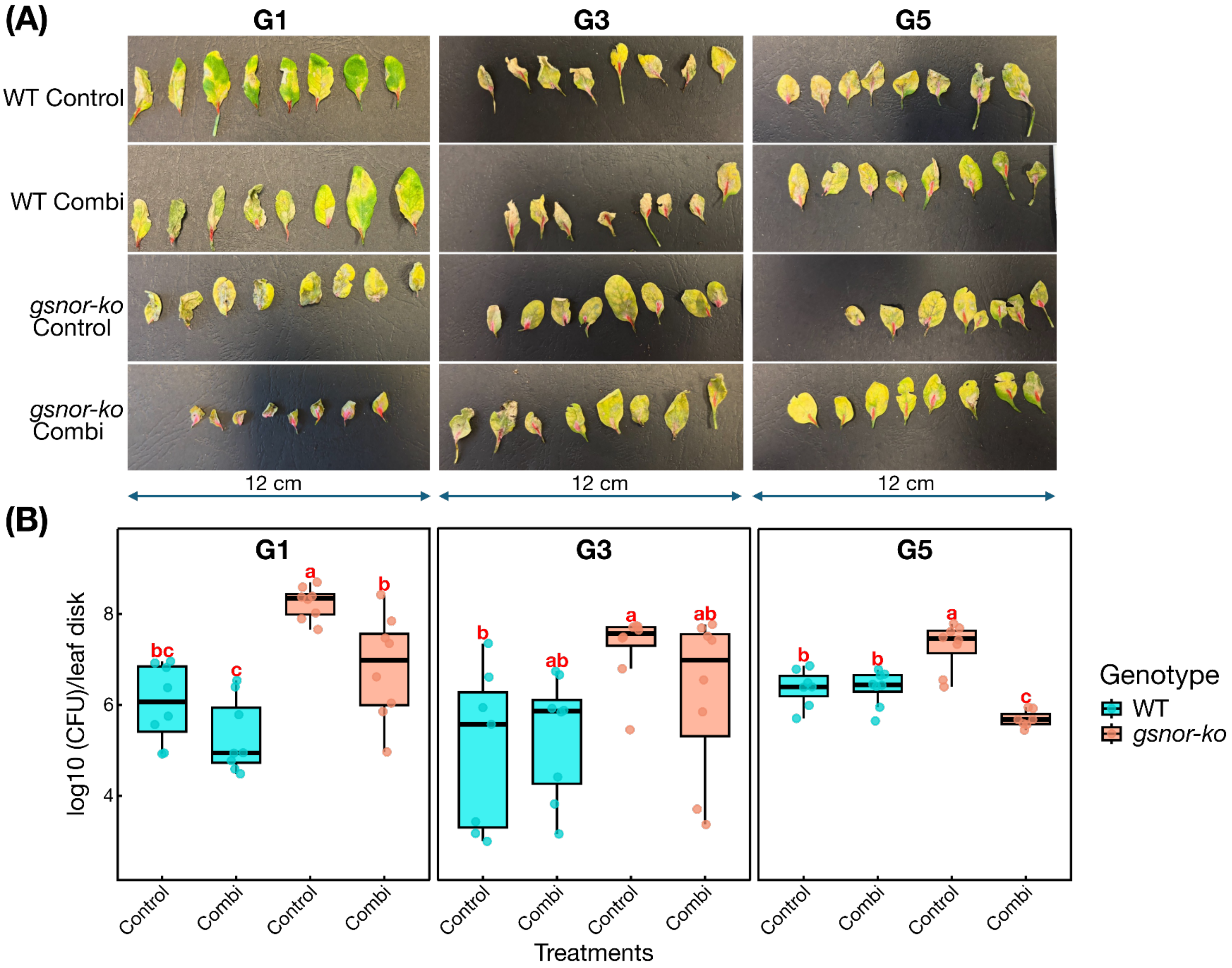

(A)
G1
G3
G5
WT Control
WT Combi
gsnor-ko Control
gsnor-ko Combi
12 cm
12 cm
12 cm
(B)
G1
G3
G5
log10 (CFU)/leaf disk
8
6
4
bc
c
a
b
b
ab
a
ab
b
b
a
c
Control
Combi
Control
Combi
Control
Combi
Control
Combi
Control
Combi
Control
Combi
Treatments
Genotype
WT
gsnor-ko

**Supplementary Fig S30. Disease assay in WT and *gsnor-ko* under control and combination treatments across generations. (A)** Representative detached leaves from WT and *gsnor-ko* plants grown under control conditions (combination having treatment history) and assayed at G1, G3, and G5. Leaves are shown after pathogen challenge, with visible differences in lesion development and tissue collapse between genotypes and treatment histories. Scale bars indicate 12 cm. **(B)** Quantification of bacterial growth in infected leaves, expressed as $log_{10}$(CFU) per leaf disk, for WT and *gsnor-ko* under control and combination conditions at G1, G3, and G5. Boxplots show the distribution of values across biological replicates; points represent individual samples. Different letters indicate significant differences among groups within each generation.

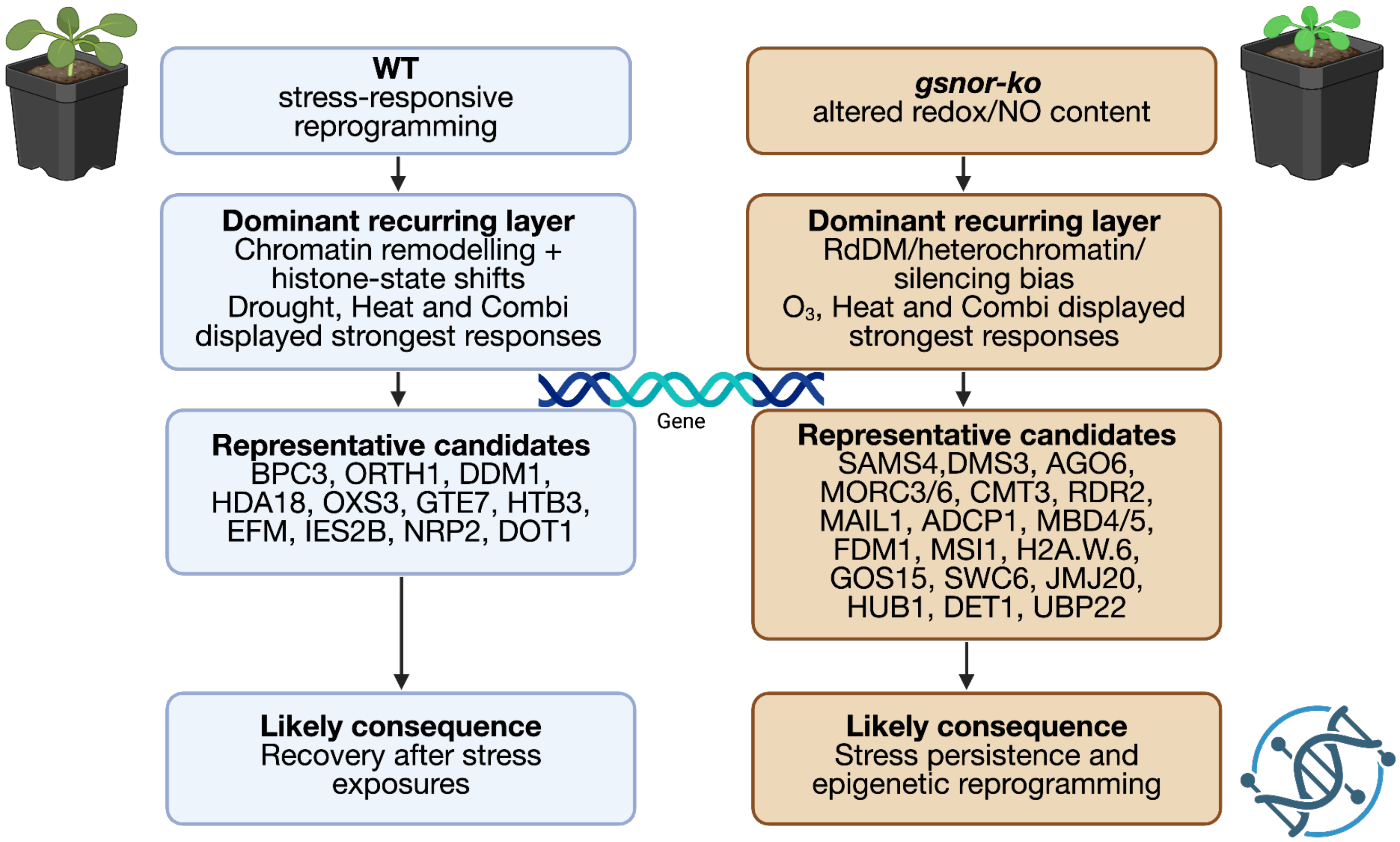


**Supplementary Fig S31. Outlook model summarising the proposed divergence between WT and *gsnor-ko* under repeated climate stress.**
